# Unveiling the Microbial Realm with VEBA 2.0: A modular bioinformatics suite for end-to-end genome-resolved prokaryotic, (micro)eukaryotic, and viral multi-omics from either short- or long-read sequencing

**DOI:** 10.1101/2024.03.08.583560

**Authors:** Josh L. Espinoza, Allan Phillips, Melanie B. Prentice, Gene S. Tan, Pauline L. Kamath, Karen G. Lloyd, Chris L. Dupont

## Abstract

The microbiome is a complex community of microorganisms, encompassing prokaryotic (bacterial and archaeal), eukaryotic, and viral entities. This microbial ensemble plays a pivotal role in influencing the health and productivity of diverse ecosystems while shaping the web of life. However, many software suites developed to study microbiomes analyze only the prokaryotic community and provide limited to no support for viruses and microeukaryotes. Previously, we introduced the *Viral Eukaryotic Bacterial Archaeal* (VEBA) open-source software suite to address this critical gap in microbiome research by extending genome-resolved analysis beyond prokaryotes to encompass the understudied realms of eukaryotes and viruses. Here we present *VEBA 2.0* with key updates including a comprehensive clustered microeukaryotic protein database, rapid genome/protein-level clustering, bioprospecting, non-coding/organelle gene modeling, genome-resolved taxonomic/pathway profiling, long-read support, and containerization. We demonstrate *VEBA’s* versatile application through the analysis of diverse case studies including marine water, Siberian permafrost, and white-tailed deer lung tissues with the latter showcasing how to identify integrated viruses. *VEBA* represents a crucial advancement in microbiome research, offering a powerful and accessible platform that bridges the gap between genomics and biotechnological solutions.

## Introduction

The microbiome is a complex community of microorganisms, encompassing prokaryotic, eukaryotic, and viral entities. This ensemble plays a pivotal role in influencing the health and productivity of diverse ecosystems while shaping the web of life. The influence of microbial activity from their interconnected metabolic processes and competition for resources propagate across trophic levels; an ecological trend that is universal from human microbiomes to extreme environmental systems.

Recently, there have been several large-scale microbiome studies where researchers have recovered tens-to-hundreds of thousands of biome-specific metagenome-assembled genomes (MAG) from biomes such as the human gut (UHGG; *N_MAG_* = 204, 938 (1)), ocean (OceanDNA; *N_MAG_* = 52, 325 (2)), and soil (*N_MAG_* = 40, 039 (3)). The aggregate knowledge derived from these large-scale studies as well as many small-to-medium scale studies have provided immeasurable insight into the taxonomic and metabolic complexity of microorganisms. However, aside from prophages, many microbiome studies focus primarily on the prokaryome (i.e., bacteria and archaea) and do not assess the eukaryome and virome from a genome-resolved perspective. The motivating factor for developing the *Viral Eukaryotic Bacterial Archaeal* (*VEBA*) open-source software suite (4) was to extend modular end-to-end (meta-)genomics/transcriptomics methodologies beyond the prokaryome to support the eukaryome and virome. The eukaryome in this context is defined as the fraction of microbes composed of nucleated organisms such as protists and unicellular fungi (adapted from (5)) while the virome consists of all the viruses within an ecosystem, including those integrated into host genomes (6).

Parasitic protists cause many human diseases such as malaria, toxoplasmosis, and giardia (7) which drives the focus of research on the eukaryome towards investigating problems related to pathogens and parasites. This emphasis on biomedical applications is essential for progressing the wellness of humankind but inadvertently cultivates a blind-spot in our knowledge around eukaryotic commensals, mutualists, and extremophiles. Despite constituting most of the eukaryotic phylogenetic diversity (8, 9), protists are frequently overlooked in investigations of extreme environments and, therefore, key opportunities have been missed to advance our understanding of the functional diversity inherent in eukaryotic life and their impact on ecosystems (10).

Viruses are submicroscopic infectious agents that replicates only inside the living cells of organisms (11). From unicellular microorganisms to complex multicellular societies of sapiens, it is hypothesized that every form of cellular life on Earth is susceptible to viral infection (12). Widespread in nearly every ecosystem, viruses are the most numerically abundant biological entity and can drive the evolution of host organisms (13). Despite their ubiquity, viruses are frequently disregarded in studies - possibly as a result from their exclusion on the tree of life, unconventional taxonomic nomenclature, or regulatory complexity - thereby underscoring the need for a more comprehensive understanding of their pervasive presence. From a public health perspective, characterizing the breadth of viral biology can provide key insight into emerging pathogens where novel viruses arise via natural processes such as recombination in diverse hosts (14). From a deeper perspective, the study of viruses from diverse ecosystems can provide insight into not only the evolution of living organisms but the origins of life itself (15).

The objective for *VEBA 1.0* was to unify robust *in silico* prokaryotic, (micro)eukaryotic, and viral computational workflows while providing seamless open-source usage for researchers globally (4). As shown in this work, the objectives for *VEBA 2.0* included optimizing the current workflows, adding complementary workflows, containerization, and expanding community-resources. In this effort, *VEBA* emphasizes the core principles of FAIR scientific stewardship with its commitment to open-source packages/databases, detailed walkthroughs, interoperability with other tools, and structured outputs for reusability (16).

Here we present a major update to *VEBA* with key highlights including a comprehensive clustered microeukaryotic protein database, rapid genome/protein-level clustering, non-coding/organelle gene modeling, genome-resolved taxonomic/pathway profiling, long-read support, and containerization. In addition, *VEBA 2.0* provides resources for the translation of genome mining results into biotechnological solutions with the addition of mobile genetic element identification, AMR gene detection, virulence factor detection, and biosynthetic potential analysis. We showcase the updates by employing *VEBA* to analyze 3 different case studies including marine water, Siberian permafrost, and white-tailed deer lung tissue metagenomes. *VEBA* is freely available as an open-source software suite and is accessible at https://github.com/jolespin/veba providing comprehensive access to its source code, datasets, and instructional walkthroughs.

## Methods

### VEBA Modules

As with the initial release, *VEBA* 2.0 maintains its modularity but now with 20 independent modules each developed for essential workflows that can be used for (meta-)genomics and/or (meta-)transcriptomics (Fig. 1).

**Figure 1.**
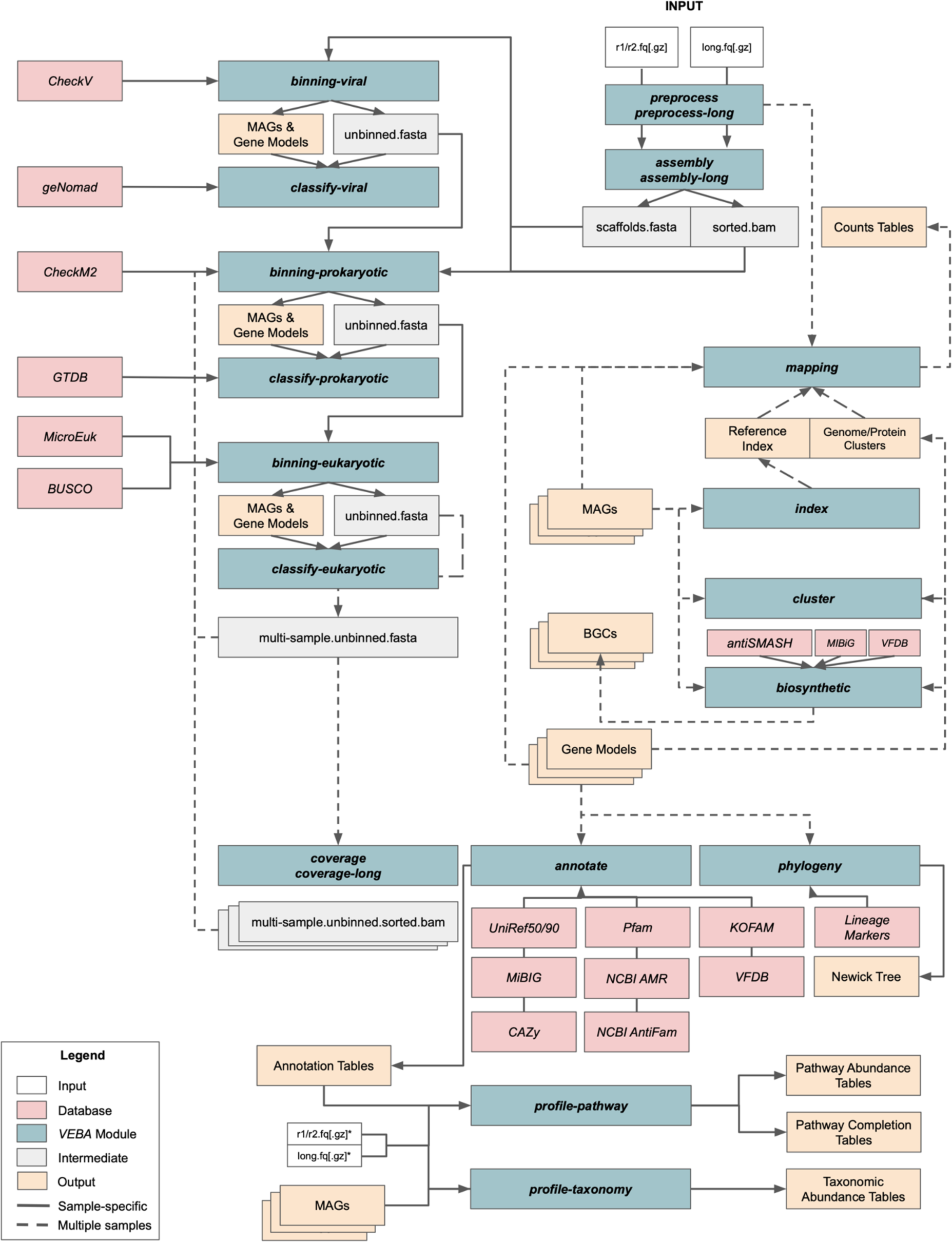
Schematic of VEBA software suite. VEBA modules and workflow I/O connectivity

### ***preprocess*** *– Fastq quality trimming, adapter removal, decontamination, and read statistics calculations (Illumina)*

The *preprocess* module is largely unchanged from the *VEBA 1.0* release. It remains as a wrapper around *fastq_preprocessor* (https://github.com/jolespin/fastq_preprocessor) which is a modernized reimplementation of *KneadData* (17) that relies on the following backend tools: 1) *fastp* for ultra-fast automated adapter removal and quality trimming (18); 2) *Bowtie2* for aligning trimmed reads to a contamination database (19) 3) *BBTools’ BBDuk.sh* (https://sourceforge.net/projects/bbmap) for profiling reads against a k-mer database (e.g., ribosomal k-mers); and 4) *SeqKit* for read accounting purposes (e.g., % contamination or % trimmed) (20). Decontamination and k-mer profiling methods are optional as are the output files for said features. The preprocess environment continues to use *Kingfisher (*https://github.com/wwood/kingfisher-download) as a backend resource for users to download sequencing files and their annotations from public data sources including *ENA, NCBI SRA, Amazon AWS,* and *Google Cloud.* For a detailed description, please refer to the initial *VEBA* publication (4).

### ***preprocess-long*** *– Fastq quality trimming, adapter removal, decontamination, and read statistics calculations (Oxford Nanopore & PacBio)*

The *preprocess-long* module uses the approach from the original *preprocess* module and applies it to long-read technologies such as *Oxford Nanopore* and *PacBio*. More specifically, the following methodology is implemented: 1) reads are quality trimmed using *NanoPack2 Chopper* (21) instead of *fastp;* 2) trimmed reads are aligned to a contamination database using *minimap2* (22) instead of *Bowtie2*; 3) *BBTools’ BBDuk.sh* for k-mer profiling (as in *preprocess*); and 4) *SeqKit* for read accounting purposes. As with the *preprocess* module, decontamination and k-mer quantification is optional.

### ***assembly*** *– Assemble reads, align reads to assembly, and count mapped reads (Illumina)*

The *assembly* module has several changes compared to the *VEBA 1.0* release. One key change is the addition of *MEGAHIT* (23) as an alternative to *SPAdes*-based assemblers (24) with *metaSPAdes* (25) remaining as the default and *rnaSPAdes* (26) as an alternative for transcriptomic workflows. With the update, sample names are prepended to the scaffolds or transcripts to bypass the possibility of duplicate identifiers that may occur in large datasets. Additional *SPAdes*-based algorithms such as *Metaviral SPAdes* are also supported but have not been thoroughly tested with *VEBA* (27). The remaining steps are similar to the initial *VEBA 1.0* assembly module which includes 1) *Bowtie2* for indexing scaffolds or transcripts and aligning reads; 2) *Samtools* for producing sorted BAM files (28); 3) *featureCounts* for counting reads that have aligned to scaffolds or transcripts (29); and 4) *SeqKit* for useful assembly quality control statistics such as N50, number of contigs, and total assembly size. As with the initial release, this module automates many critical yet overlooked workflows dealing with assemblies that are typically performed *post hoc* such as contig-level sequence alignment, contig-level counts tables, assembly indexing, and assembly statistics.

### ***assembly-long*** *– Assemble reads, align reads to assembly, and count mapped reads (Oxford Nanopore & PacBio)*

The *assembly-long* module uses a similar approach of the *assembly* module but using packages designed for long reads instead. For instance, instead of *SPAdes*-based assemblers the module uses *Flye* (30) and *MetaFlye* (31) where the sample name is prepended to the contigs. After (meta-)genomic assembly is finished and assembly statistics are computed with *SeqKit*, long reads are aligned back to the assembly using *minimap2* and BAM files are sorted with *Samtools*. Sorted BAM files are counted using *featureCounts* in long reads mode.

### ***coverage*** *– Align reads to a (multi-sample) reference and counts mapped reads (Illumina)*

The *coverage* module is largely unchanged from initial release and is used to produce mapping index objects and sorted BAM files. Details have been described previously (4).

### ***coverage-long*** *– Align reads to a (multi-sample) reference and counts mapped reads (Oxford Nanopore & PacBio)*

The *coverage-long* module follows the adaptation strategy of *preprocess-long* and *assembly-long* in which *Bowtie2* is replaced with *minimap2*. The approach for *coverage-long* is the same as coverage but with tools designed for long reads.

### ***binning-prokaryotic*** *– Iterative consensus binning for recovering prokaryotic genomes, modeling coding/non-coding genes, and quality assessment*

The fundamentals of the prokaryotic binning module have not changed relative to the initial *VEBA 1.0* release. Namely, *CoverM* (https://github.com/wwood/CoverM) is used for fast coverage calculations, multiple binning algorithms (*MaxBin2* (marker set = 107); *MaxBin2* (marker set = 40) (32); *MetaBat2* (33); and *CONCOCT* (34), a genome size filter (150, 000 bp is the default), consensus binning with *DAS Tool* (35), and a consensus domain wrapper for *Tiara* (36) for removing eukaryotes at the MAG level. The module still maintains its iterative functionality where unbinned contigs are fed back into the start of the algorithm as these unbinned contigs can be viewed as a new lower complexity space to cluster contigs into bins. The main updates include the use of *CheckM2* (37) to massively speed up genome quality assessments with direct support for CPR (CPR support was available in prior versions but required a workaround as detailed later). Coding sequences are now modeled using *Pyrodigal* (38) instead of *Prodigal* (39). For non-coding genes, *Barrnap* (https://github.com/tseemann/barrnap) is used for ribosomal RNA (rRNA) and *tRNAscan-SE* 2.0 is used for transfer RNA (tRNA) (40). MAG naming scheme for prokaryotes follows [SampleID] [Algorithm] P.[Iteration] [Name] (e.g., *SRR13615824 METABAT2 P.1 bin.3*).

### ***binning-eukaryotic*** *– Binning for recovering eukaryotic genomes with nuclear/organelle (exon-aware) coding/non-coding gene modeling and lineage-specific quality assessment*

The fundamentals of the eukaryotic binning module have not changed relative to the initial *VEBA 1.0* release. Namely, the following protocol: 1) coverage calculated with *CoverM*; 2) non-prokaryotic-specific binning algorithms *MetaBat2* or *CONCOCT* for binning out genomes followed by a genome size filter (2, 000, 000 bp is the default); 3) *Tiara* to predict eukaryotic MAGs and remove non-eukaryotic MAGs; 4) coding/non-coding gene modeling for nuclear and organelle genomes; 5) *BUSCO* for lineage-specific quality assessment and the removal of poor quality genomes (41); 6) *featureCounts* for gene-level counts; and 7) genome statistics calculated with *SeqKit*. *Tiara* is now also used for partitioning nuclear, mitochondrion, and plastid genomes where the respective rRNA and tRNA gene modeling parameters are used for *Barrnap* and *tRNAscan-SE 2.0*, respectively. *MetaEuk* is still used for exon-aware gene modeling in nuclear genomes but the database has been updated to include comprehensive clustered options as described in detail below. Coding gene modeling for organelles are called with *Pyrodigal* using organelle-specific genetic code translation tables (e.g., default uses trans_table=4 for mitochondrion and trans_table=11 for plastid genomes). The coding and non-coding gene modeling for nuclear and organelle genomes is wrapped in a standalone script called *eukaryotic_gene_modeling.py* which allows users to model genes that are acquired outside of *VEBA*. MAG naming scheme for eukaryotes follows [SampleID] [Algorithm] E.[Iteration] [Name] (e.g., *ERR2002407 METABAT2 E.1 bin.2*) where iteration currently always is 1 but is used as a placeholder for future methodologies where iterative eukaryotic binning is supported.

### ***binning-viral*** *– Detection of viral genomes and quality assessment*

The viral binning module has been entirely reimplemented. Viral binning is performed using either *geNomad* (default) (42) or *VirFinder* (43) to identify candidate viral genomes on a per contig basis. Genes are modeled using a modified version of *Prodigal* designed specifically for viruses called *Prodigal-GV* (42). The candidate viral genomes are then input into *CheckV* (44) where quality assessment removes poor quality or low confidence viral predictions. The default filtering scheme (recommended by *CheckV* developers (44, 45)), is summarized by the following: 1) number of viral genes ≥ 5 × number of host genes; 2) completeness ≥ 50%; 3) *CheckV* quality is either medium-quality, high-quality, or complete; and 4) *MIUViG* quality is either medium-quality, high-quality, or complete. If *geNomad* is selected, then candidate plasmid sequences (along with any conjugation or AMR genes) are identified. Iterative binning is not applicable for viral detection as algorithms are executed on a per-contig basis and all viral genomes will be identified on first pass. MAG naming scheme for viruses follows [SampleID] [Algorithm] [Name] (e.g., SRR9668957 GENOMAD Virus.1).

### ***classify-prokaryotic*** *– Taxonomic classification of prokaryotic genome<u>s</u>*

The prokaryotic classification module is a useful wrapper around the updated *GTDB-Tk 2.0* (46) and the r214.1 *GTDB* release (47). If genome clusters are provided, then it performs consensus lineage classification. A mash prescreen is used to screen organisms based on ANI using the following database we provide as an unofficial mirror of the mash sketched r214.1 *GTDB* (https://zenodo.org/records/8048187). Interactive *Krona* graphs are generated in HTML and tabular format for prokaryotic taxonomy.

### ***classify-eukaryotic*** *– Taxonomic classification of eukaryotic genomes*

As in the prior release, the eukaryotic classification module utilizes genes from *BUSCO’s eukaryota_odb10* marker set (and the curated score cutoffs), the target field of *MetaEuk* gene identifiers, and the taxonomic lineage associated with each source genome. The eukaryotic classification module can reuse previous intermediate files if genomes were binned using *VEBA* but also provides a workflow for classifying eukaryotic taxonomy from genome assemblies which runs *MetaEuk* in the background with *MicroEuk100.eukaryota_odb10*. The classification protocol has been previously described (4) and the only modification is that it can now handle taxa with incomplete lineages such is the case for many protists and fungi. If genome clusters are provided, then it performs consensus lineage classification. Interactive *Krona* graphs are generated in HTML and tabular format for eukaryotic taxonomy.

### ***classify-viral*** *– Taxonomic classification of viral genomes*

The viral classification module utilizes *geNomad’s* taxonomy if provided or runs the *geNomad taxonomy* module if genome assemblies are provided from a different source. If genome clusters are provided, then it performs consensus lineage classification.

### ***cluster*** *– Species-level clustering of genomes and pangenome-specific protein clustering*

The *cluster* module has been completely reimplemented using state-of-the-art tools designed specifically for genome- and protein-level clustering. The *cluster* module first uses either *Skani* (default) or *FastANI* (48) to compute pairwise ANI and these are used to construct a *NetworkX* graph objects for each organism type where nodes are genomes and weighted edges are ANI values (49). These *NetworkX* graphs are converted into subgraphs of connected components whose edges are filtered by a particular threshold (default ≥ 95% ANI) for species-level clustering. Proteins are partitioned by these species-level clusters (SLC) and protein clustering is performed on each pangenome to yield SLC-specific protein clusters (SSPC). *VEBA 2.0* now supports *MMseqs2* (default) (50) and *Diamond DeepClust* (51) in both sensitive clustering and linclust modes. These are provided with a convenient wrapper called *clustering_wrapper.py* that can be used for clustering in protein-space (*MMseqs2* and *Diamond DeepClust*) and nucleotide-space (*MMseqs2* only). This wrapper clusters sequences, relabels representatives, and provides representative sequences in either fasta or tabular format. Clustering is performed for prokaryotic, eukaryotic, and viral organisms separately. By default, clustering is global (i.e., containing all samples) with an optional support for local clustering on a sample-specific basis. Genomic and functional feature compression ratios are calculated for prokaryotic, eukaryotic, and viral organisms separately as described previously (4). The nomenclature preferred by *VEBA* is the PSLC-, ESLC-, and VSLC-for the prefix of each genome cluster (e.g., *PSCL-0*) with SSPC-appended for protein clusters (e.g., PSLC-0_SSPC-10) which indicates protein cluster 10 from pangenome 0 but this can be customized.

### ***annotate*** *– Functional annotation of protein sequences*

Annotation is performed using best-hit annotations and profile HMMs. *Diamond* (52, 53) is used to align proteins to the following databases: 1) *UniRef50* (default) or *UniRef90* (54); 2) *MIBiG* (55); 3) *VFDB* (56); and *CAZy* (57). *HMMER* (58) is used to profile sequences for protein domains using the following databases: 1) the *Pfam* database (59); 2) *NCBIfam-AMR* (60); 3) *AntiFam* (61); and 4) *KOFAM* (62). If clustering is performed prior, then the identifier mapping table can be provided as input to produce additional output which includes: 1) consensus annotations for each protein-level cluster; 2) *KEGG* module completion ratios for genomes and SLCs using a custom *MicrobeAnnotator* build (63) from KEGG orthologs identified via *KofamScan*.

### ***biosynthetic*** *– Identify biosynthetic gene clusters and cluster in both protein and nucleotide space*

The *biosynthetic* module is a new addition to *VEBA* 2.0 and is described in detail later under “*New features and updates* section below”. In brief, the *biosynthetic* module does the following: 1) identifies candidate biosynthetic gene clusters (BGC) and secondary metabolite pathways using *antiSMASH* (64); 2) converts the GenBank formatted outputs into tabular and fasta formats; 3) compiles *Krona* graphs of BGC protocluster types nested within genomes; 4) aligns BGC proteins to *MIBiG* and *VFDB* using *Diamond*; 5) calculates novelty scores by identifying the ratio of proteins that have no homology to *MIBiG*; 6) clusters BGCs in protein-space via *MMseqs2* and builds prevalence tables relative to genomes; and 7) clusters full-length BGCs in nucleotide-space via *MMseqs2* and builds prevalence tables relative to samples of origin. This module includes a convenient script called *biosynthetic_genbanks_to_table.py* to parse GenBank files from a particular genome-specific *antiSMASH* run to build with biosynthetic information relative to the protocluster-type, BGC, and BGC proteins which can be used separately.

### ***profile-taxonomy*** *-Taxonomic profiling of de novo genomes*

The *profile-taxonomy* module is another new addition to *VEBA* 2.0 and is described in detail later under “*New features and updates* section below”. In brief, the *profile-taxonomy* module does the following: 0) builds a *Sylph* sketch database (65) for non-viral and viral genomes using the *compile_custom_sylph_sketch_database_from_genomes.py* script prior to running the module; 1) converts paired reads to a query sketch database using *Sylph;* 2) profiles the genome sketch databases using the query sketch database generated from the reads; 3) reformats the *Sylph* output tables; and 4) aggregates abundances with respect to SLC if clusters are provided.

### ***profile-pathway*** *- Pathway profiling of de novo genomes*

The *profile-pathway* module is yet another new addition to *VEBA* 2.0 and is described in detail later under “*New features and updates* section below”. In brief, the *profile-pathway* module does the following: 0) builds a custom *HUMAnN* database based on protein annotations, identifier mapping tables, and taxonomy assignments using the *compile_custom_humann_database_from_annotations.py* script prior to running the module; 1) either accepts pre-joined reads, joins paired-end reads using *bbmerge.sh* from *BBSuite*, or a BAM file of paired-end reads and joins them; 2) builds a *Diamond* database of proteins from the custom *HUMAnN* annotation table; 3) uses *HUMAnN* for pathway profiling of the joined reads using the custom *HUMAnN* database (17); and 4) reformats the output files.

### ***phylogeny*** *– Constructs phylogenetic trees based on concatenated alignments of marker genes*

The phylogeny module has only minor changes relative to the initial *VEBA 1.0* release. Briefly, the following methodology is performed: 1) identifying marker proteins using *HMMSearch* from the *HMMER* suite; 2) creating protein alignments for each marker identified via *MUSCLE5* (66); 3) trimming the alignments using *ClipKIT* (67); 4) concatenating the alignments; 5) approximately-maximum-likelihood phylogenetic inference using either *FastTree2* (default) (68) or *VeryFastTree* (69); and 6) optional maximum likelihood phylogenetic inference using *IQ-TREE2* (70). More details such as using scoring parameters or determining the minimum number of genomes or minimum markers to include have been described previously (4).

### ***index*** *– Builds local or global index for alignment to genomes*

The *index* module is largely unchanged from initial release and is used to produce mapping index objects. Currently, *Bowtie2* (19) is the only alignment software packages supported. Details have been described previously (4).

### ***mapping*** *– Aligns reads to local or global index of genomes*

The *mapping* module is largely unchanged from initial release and is used to generate counts tables at the genome, SLC, gene, and SSPC levels. Currently, *Bowtie2* (19) is the only alignment software packages supported but wrappers for *STAR* are provided for exon-aware read mapping (71). Details have been described previously (4).

#### Compositional network analysis and community detection

To build compositionally-valid association networks (i.e., co-occurrence) we implemented the following strategy: 1) profiling the taxonomic abundances at the genome and SLC-level using *VEBA profile-taxonomy* module; 2) computing compositionally-valid ensemble association networks (*N_Draws_* = 100) (72) using partial correlation with basis shrinkage (73–75) with the SLC abundances; 3) subsetting only the positive associations; 4) computing consensus Leiden communities (*N_Seeds_* = 100) (76); and 5) calculating the weighted degree for each SLC and ranking by connectivity.

#### Visualization

Genomic neighborhoods were visualized using *DNA Features Viewer v3.1.3* (77). Phylogenetic trees were constructed from multiple sequence alignments computed by *MUSCLE v5.1* (78) and visualized using *Toytree v2.0.5* (79). Multiple sequence alignments were visualized using *pyMSAviz v0.4.2* (https://moshi4.github.io/pyMSAviz/). CRISPR-Cas systems were identified in genomes recovered from *VEBA* using *CRISPRCasTyper* v1.8.0 (80) which generates plots using *drawsvg v2.3.0* (https://github.com/cduck/drawsvg). Association networks and genome cluster networks were visualized using *NetworkX v3.2.1* (49). Clustered heatmaps and bar charts were visualized using *Seaborn v0.13.2* (81) and *Matplotlib v3.8.3* (82).

## Results and Discussion

### Walkthroughs and workflow tutorials

In *VEBA’s* mission toward FAIR principles and widespread accessibility, we have compiled several walkthroughs on our *GitHub*. We provide multiple end-to-end workflows including a complete metagenomics analysis which covers assembling metagenomic reads, binning, clustering, classification, and annotation. In a similar vein, we provide a walkthrough for recovering viruses from metatranscriptomics datasets which covers assembling metatranscriptomic reads, viral binning, clustering, and classification. We also show how to use the unbinned contigs in a pseudo-coassembly, a concept described in the initial *VEBA* publication (4), with guidelines on when this should be performed. Lastly, we provide a walkthrough for setting up a *bona fide* co-assembly for metagenomics or metatranscriptomics which may be useful in scenarios where all or most samples are of low sequencing depth relative to the microbial diversity that is present. This walkthrough goes through concatenating reads, creating a reads table, co-assembly of concatenated reads, aligning sample-specific reads to the co-assembly for multiple sorted BAM files, and mapping reads for scaffold/transcript-level counts. For abundance estimation walkthroughs, we cover traditional approaches for aligning reads using the *mapping* and *index* modules as well as profiling approaches for both genome-resolved taxonomic abundance and pathway profiling using genomes identified through *VEBA* or elsewhere. Lastly, we provide walkthroughs on converting counts tables to *anndata* format for integration with *scverse* (83) and *BIOM* format (84) for integration with *QIIME2* (85). We also include additional miscellaneous walkthroughs such as downloading/preprocess fastq files from NCBI, phylogenetic inference, bioprospecting for BGCs, screening for CRISPR-Cas systems, and adapting commands for use with *Docker* containers.

### New features and updates

#### Expanded functionality, streamlined user-interface, and Docker containerization

The updated *VEBA 2.0* includes 20 modules and 95 accessory scripts to streamline workflows (Fig. 1, Table 1) with 51 peer-reviewed software dependencies and 21 databases (Table S1). Since the initial release, *VEBA* has included hundreds of *GitHub* commits to add new features suggested by the user-base and to address issues when flagged by the community. As each module requires a unique set of dependencies, groups of similar modules (e.g., *classify-prokaryotic, classify-viral,* and *classify-eukaryotic*) use shared *Conda* environments where dependencies are installed (e.g., *VEBA-classify_env*). While *VEBA 1.0* required users to activate specific *Conda* environments for each workflow this is now automated with a convenient wrapper program. For example, the previous functionality required the following syntax: ‘source activate VEBA-preprocess_env && preprocess.py ${PARAMS}’ while the current functionality is streamlined to ‘veba--module preprocess--params “${PARAMS}”’ where the *Conda* environment is abstracted and determined automatically in the backend.

**Table 1.** VEBA modules.

| Module | Description | Docker Registry |
| --- | --- | --- |
| **preprocess** | Fastq quality trimming, adapter removal, decontamination, and read statistics calculations (short reads) | https://hub.docker.com/r/jolespin/veba_preprocess |
| **preproces-long** | Fastq quality trimming, adapter removal, decontamination, and read statistics calculations (long reads) | https://hub.docker.com/r/jolespin/veba_preprocess |
| **assembly** | Assemble reads, align reads to assembly, and count mapped reads (short reads) | https://hub.docker.com/r/jolespin/veba_assembly |
| **assembly-long** | Assemble reads, align reads to assembly, and count mapped reads (long reads) | https://hub.docker.com/r/jolespin/veba_assembly |
| **coverage** | Align reads to (concatenated) reference and counts mapped reads (short reads) | https://hub.docker.com/r/jolespin/veba_assembly |
| **coverage-long** | Align reads to (concatenated) reference and counts mapped reads (long reads) | https://hub.docker.com/r/jolespin/veba_assembly |
| **binning-prokaryotic** | Iterative consensus binning for recovering prokaryotic genomes with lineage-specific quality assessment | https://hub.docker.com/r/jolespin/veba_binning-prokaryotic |
| **binning-eukaryotic** | Binning for recovering eukaryotic genomes with exon-aware gene modeling and lineage-specific quality assessment | https://hub.docker.com/r/jolespin/veba_binning-eukaryotic |
| **binning-viral** | Detection of viral genomes and quality assessment | https://hub.docker.com/r/jolespin/veba_binning-viral |
| **classify-prokaryotic** | Taxonomic classification of prokaryotic genomes | https://hub.docker.com/r/jolespin/veba_classify |
| **classify-eukaryotic** | Taxonomic classification of eukaryotic genomes | https://hub.docker.com/r/jolespin/veba_classify |
| **classify-viral** | Taxonomic classification of viral genomes | https://hub.docker.com/r/jolespin/veba_classify |
| **cluster** | Species-level clustering of genomes and lineage-specific orthogroup detection | https://hub.docker.com/r/jolespin/veba_cluster |
| **annotate** | Annotates translated gene calls several databases | https://hub.docker.com/r/jolespin/veba_annotate |
| **phylogeny** | Constructs phylogenetic trees given a marker set | https://hub.docker.com/r/jolespin/veba_phylogeny |
| **index** | Builds local or global index for alignment to genomes | https://hub.docker.com/r/jolespin/veba_mapping |
| **mapping** | Aligns reads to local or global index of genomes | https://hub.docker.com/r/jolespin/veba_mapping |
| **biosynthetic** | Identify biosynthetic gene clusters in prokaryotes and fungi | https://hub.docker.com/r/jolespin/veba_biosynthetic |
| **profile-pathway** | Pathway profiling of de novo genomes | https://hub.docker.com/r/jolespin/veba_profile |
| **profile-taxonomy** | Taxonomic profiling of de novo genomes | https://hub.docker.com/r/jolespin/veba_profile |

Many of *VEBA’s* software dependencies are incompatible in the same compute environment. While *VEBA’s* installation process remains streamlined with partitioning compute environments for modules with similar dependencies in specific *Conda* environments, *VEBA 2.0* now provides an option for containerization support via *Docker.* Each *Conda* environment has been prepackaged into *Docker* containers for seamless usage on local machines and high-performance compute servers where containers are supported (e.g., *AWS*). Each *Docker* container comes equipped with input, output, and database mount points that allow for generalized syntax. As the switch from *Conda*-based workflows to containerized solutions can have a steep learning curve, *VEBA’s* documentation also provides walkthroughs that guide the user through pulling and running containers on their local machine or on AWS.

#### MicroEuk100/90/50: Clustered database of ∼80M microeukaryotic protein sequences

*VEBA’s Microeukaryotic Protein Database* has been completely redesigned using the logic of *UniRef* and their clustered database (54). The initial microeukaryotic protein database from the previous publication, hereby referred to as *MicroEuk_v2*, contained 48, 006, 918 proteins from 44, 647 source organisms while the updated database, *MicroEuk_v3*, contains 79, 920, 430 proteins from 52, 495 source organisms (https://zenodo.org/records/10139451). As in the prior major release, *MicroEuk_v3* concentrates on microeukaryotic organisms while excluding higher eukaryotes, as the former are the most common eukaryotes captured by shotgun metagenomics and metatranscriptomics. Source organisms in this context are defined as organisms from which the proteins were derived.

*MicroEuk_v3* is built using the following logic: 1) remove stop codons if they exist; 2) filter proteins that are less than 11 AA; 3) convert protein sequence to a unique md5 hash to use as the identifier; and 5) add protein if it does not already exist in the database. The removal of stop codons serves 2 functions; first, it creates slightly smaller file sizes and second, more importantly, it ensures that 2 identical proteins that differ only in the presence of a stop codon will have the same md5 hash for true dereplication. The removal of proteins less than 11 AA long is to avoid greedy clustering as implemented by *UniRef* (54). The databases are added with the following priority: 1) *JGI MycoCosm* (86), 2) *JGI PhycoCosm* (87), 3) *EnsemblProtists* (88), 4) *MMETSP* (89), 5) *TARA SAGv1* (90), 6) *EukProt* (91), 7) *EukZoo* (92), 8) *TARA SMAGv1* (93), and 9) *NCBI non-redundant (protists and fungi)* (94) as detailed in Table 2. The majority of *MicroEuk* proteins (86.8%) are either genome or transcriptome resolved (source organisms) from databases 1-8 while the remaining protein sequence space is padded with *NCBI’s non-redundant* database (hereby referred to as *nr*) to provide additional context for eukaryotic gene modeling and classification as these have reliable taxonomic identifiers.

**Table 2.**
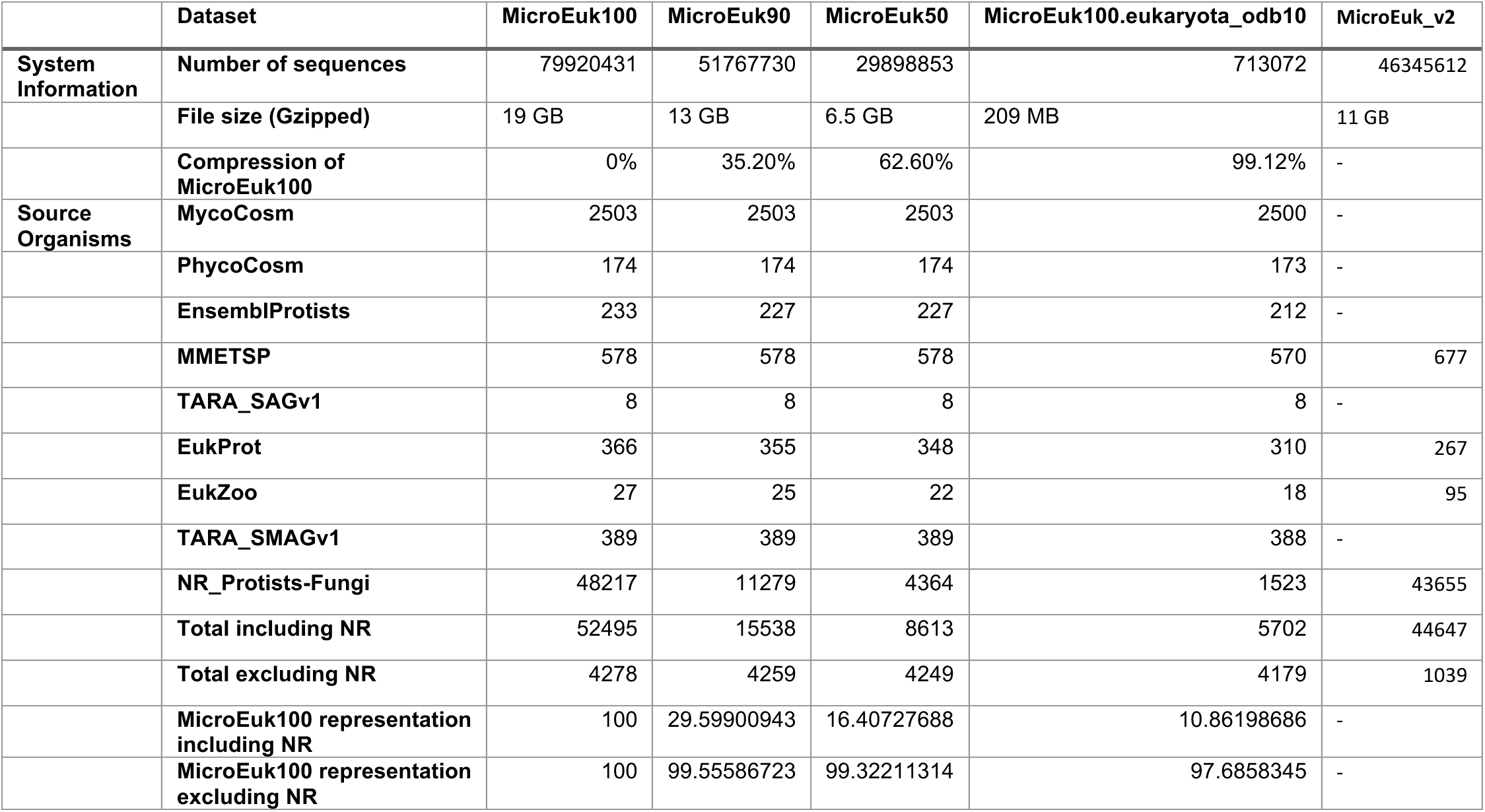
MicroEuk100/90/50.

As *UniRef100* contains non-redundant proteins, *MicroEuk100* contains the comprehensive non-redundant database of microeukaryotic proteins while *MicroEuk90* and *MicroEuk50* are clustered at 90% identity and 50% identity (80% query coverage) in a stepwise manner. Excluding the proteins padded from *nr*, the clustered databases include most of the genome/transcriptome-resolved source organisms from *MicroEuk100* with *MicroEuk90*, *MicroEuk50*, and *MicroEuk100.eukaryota_odb10* at 99.56%, 99.32%, and 97.69%, respectively (Table 2). The *MicroEuk100.eukaryota_odb10* is a subset of *MicroEuk100* that contains only hits from *BUSCO*’s *eukaryota_odb10* marker database which is used for eukaryotic classification.

#### Fast and memory-efficient genome- and protein-level clustering

*VEBA 2.0* has substantially improved the efficiency and scalability of both genome- and protein-level clustering by incorporating the most cutting-edge open-source tools available. Until 2023, *FastANI* had been the main tool used for pairwise genome ANI calculations. However, to calculate pairwise associations the genome must be loaded into an index twice followed by *N^2^* pairwise ANI calculations. While this is not a problem for small to mid-sized datasets, this implementation can cause significant performance issues for larger datasets. Recently, there has been a major innovation for ANI calculations with *skani* (∼25X faster than *FastANI*) which has a module designed specifically for pairwise ANI calculations that loads the genomes into an index only once and then performs *(N^2^ – N)/2* calculations. In addition to increased performance, *skani* also calculates alignment factors and confidence intervals for ANI between genomes. Further *skani* has higher performance on fragmented genomes (typical of metagenome-assembled genomes) and small genomes (e.g., viruses) with tunable parameters. *VEBA 2.0* provides presets for clustering various genomes based on guidelines detailed in the *skani* documentation and sets *skani* as the default ANI calculation algorithm while providing support for *FastANI* to retain previous functionality. As in previous versions, *NetworkX’s connected_components* algorithm is used to identify species-level genome clusters (SLC). In addition, the *NetworkX* graph objects are serialized for seamless downstream usage such as generating constellation plots of genome clusters.

While the performance gains of genome clustering have been improved, the most substantial improvements have been implemented in protein space. *VEBA 1.0* used *OrthoFinder* for clustering in protein space. While *OrthoFinder* is useful for detailed protein orthology, it was not designed for large-scale protein clustering as it creates several output files per protein cluster including alignments and phylogenetic trees allocating a significant amount of time dedicated to system I/O for hundreds of thousands of intermediate files. *VEBA 2.0* now includes options for using either *MMseqs2* or *Diamond’s DeepClust* algorithm. With either *MMseqs2* or *DeepClust* used in the backend, *VEBA* can not only calculate protein clusters in linear time with the *linclust* algorithm but also able to output representative sequences for each protein making downstream tasks such as annotation less resource intensive. While proteins are only clustered within a SLC to yield SLC-specific protein clusters (SSPC), *VEBA* provides wrappers around *MMseqs2* and *DeepClust* for other clustering tasks such as the large-scale protein clustering used to produce *MicroEuk100/90/50* databases.

In addition to the essentials of protein clustering, *VEBA 2.0* also produces protein-cluster prevalence tables which are then used to identify core proteins within SLCs (i.e., pangenomes) and singleton proteins that are only in a single genome within a SLC. These core proteins are output into separate fasta formatted files for each SLC in both protein and nucleotide space which can be used for downstream analysis such as marker gene detection or ratios of non-synonymous to synonymous mutation (dN/dS) calculations used in evolutionary analyses. Further, *VEBA* also includes an independent script called *marker_gene_clustering.py* for determining marker genes from these protein-cluster prevalence tables which identifies marker genes that are core within a pangenome and not detected in any other pangenomes. The marker gene detection capabilities of *VEBA* 2.0 are designed to be used for building custom profiling databases such as *Metaphlan4* (95) or *MIDAS2* (96).

#### Automatic calculation of feature compression ratios

In *VEBA 1.0*, the concept of a feature compression ratio (FCR) was introduced as a measure of a community’s complexity from an unsupervised perspective. While the theory of FCR was established, there was no automated calculation of this ratio within *VEBA* 1.0 and the calculation needed to be performed *post hoc*. To address this issue, *VEBA* 2.0 automates FCR calculations in both genome- and protein-space. To fully leverage the power of FCR when analyzing the complexity of a dataset, *VEBA* 2.0 has adapted clustering to be performed both at the global and local level where global and local refer to inter- and intra-sample clustering, respectively, with local clustering turned off by default. This functionality allows FCR calculations to be calculated for an entire dataset but also for each sample within a dataset. The FCR calculations are expected to provide useful insight on the complexity of a dataset or sample by providing a metric for the redundancy in genome and protein space (i.e., taxonomy and functionality).

#### Large/complex metagenomes and long-read technology support

Since version 1.0, *VEBA* has used *SPAdes*-based programs such as *SPAdes*, *metaSPAdes*, and *rnaSPAdes* for genomic, metagenomic, and transcriptomic assembly. While the resulting assemblies are high quality, *SPAdes*-based assemblers can be resource intensive with large and complex genomes such as soil and sediment metagenomes. *MEGAHIT* is another assembler that has more accommodating memory requirements than *SPAdes*-based assemblers and is particularly useful for large and complex metagenomes (23, 97). While *metaSPAdes* remains the default assembler in *VEBA 2.0*, there is now an option to use *MEGAHIT* with seamless access to the large and complex preset. In addition to adding a new assembler, *VEBA* 2.0 also addresses the critical, albeit rare, issue of more than one contig having the same name which can occur with very large complex datasets with many samples. To bypass this issue, *VEBA* prepends the sample name to the contig identifiers by default but this can be customized.

With the increased accuracy and widescale adoption of long-read technologies such as *Oxford Nanopore* and *PacBio* in genomics, transcriptomics and their meta-counterparts, the need for *VEBA* to accommodate long reads was inevitable to properly address the directive of genome-resolved metagenomics. To accommodate long reads, *VEBA* has restructured *fastq_preprocessor* (a light-weight extension of *VEBA* and the basis for the *preprocess* module) to include short-read and long-read modules wrapped natively by *VEBA* with the latter including *chopper* and *minimap2* instead of *fastp* and *bowtie2*. *VEBA* now also provides the *assembly-long* module which includes *Flye*, *MetaFlye*, and *Minimap2* instead of *SPAdes, metaSPAdes,* and *bowtie2.* Starting with *Flye* v2.9+, which is included with *VEBA 2.0*, users are no longer required to specify an assembly size making general usage much more accessible especially for metagenomics. The coverage calculation has been adapted with *coverage-long* which includes the same coverage method but using *Minimap2* instead of *Bowtie2*. Last of all, the viral, prokaryotic, and eukaryotic binning modules now incorporate long reads as well.

#### Bioprospecting and natural product discovery support

*antiSMASH* is a very popular package that is used for secondary metabolite identification and characterization in natural product discovery (64). However, interpreting the results in high-throughput can be challenging as the output formats include JSON and GenBank both of which are verbose and difficult to query for multiple genomes. To provide increased accessibility to high-throughput bioprospecting, the *VEBA 2.0* update includes a new module called *biosynthetic* that runs *antiSMASH* on each genome, assigns structured identifiers for each biosynthetic gene cluster and each gene within the cluster, constructs multiple outputs that can be queried for multiple genomes, scores the novelty of the BGC, and clusters the BGCs.

The first stage of *VEBA biosynthetic* is an *antiSMASH* wrapper and provides the following output: 1) tabular outputs at both the BGC and gene level; 2) fasta file for each genome containing BGCs in nucleotide space with informative attributes in the description such as the BGC length, GC-content, number of genes, and whether or not the BGC is on the edge of contig; and 3) fasta file for each genome containing the proteins for all the BGCs. The structured identifier scheme for BGCs is *[id_genome]|[id_contig]|[id_region]* while the scheme for BGC genes is *[id_bgc]_[gene_position_on_bgc]|[start_on_contig]:[end_on_contig]([strand])*. For instance, the BGC gene *SRR13615824 METABAT2 P.1 bin.3|SRR13615824 k127_496383|region001_1| 1:184(+)* is the first gene found on the *SRR13615824 METABAT2 P.1 bin.3|SRR13615824 k127_496383|region001* BGC from genome *SRR13615824 METABAT2 P.1 bin.3* between position 1-184 in the positive direction of contig *SRR13615824 k127_496383*.

The second stage of the *biosynthetic* module aligns all the translated BGC genes to *MIBiG* (55) and *VFDB* (56). The ratio of BGC genes that have homology to *MIBiG* is used to compute the novelty score for a BGC. The novelty score was developed for *VEBA 2.0* to quantify the percentage of genes within a BGC that have no homology to *MIBiG* and, thus, can be used for flagging BGCs with potentially novel activity. The annotations are summarized into a table that can be used for investigating individual protein products within a BGC.

The third and final stage of the *biosynthetic* module clusters BGCs in both protein and nucleotide space using *MMseqs2* each with their own set of parameters for minimum coverage and identity. The proteins are clustered to produce a BGC protein cluster where identifier mapping tables, representative sequences, and prevalence tables are output. The prevalence table is structured with the genomes as the rows, BGC protein clusters as the columns, and the number of BGC proteins in a genome that are within each cluster. For the nucleotide clustering, *MMseqs2* is also used for clustering with identifier mapping tables, representative sequences, and prevalence tables being the output as well. However, the prevalence tables for BGC nucleotide clusters contain samples for the rows and BGC nucleotide clusters for the columns. The prevalence tables can easily be assessed post hoc to identify singleton BGCs and core BGCs. Further, the prevalence tables can be converted to a Boolean matrix and pairwise distance for the genomes or samples can be computed using Jaccard distance which can be directly integrated into unsupervised machine learning methodologies such as principal coordinate analysis or agglomerative hierarchical clustering.

#### Ribosomal RNA, transfer RNA, and organelle support

While *VEBA 1.0* focused mainly on protein-coding sequence (CDS) genes, the *VEBA 2.0* implementation automates the detection of ribosomal RNA (rRNA) and transfer RNA (tRNA) using *Barrnap* and *t-RNAscan-SE 2.0*, respectively, which are now added directly to the GFF gene modeling output. In addition to fasta and GFF records for non-coding RNA, secondary structures are also provided for tRNA sequences. For eukaryotic gene modeling, *VEBA* 2.0 automates the identification of plastid and mitochondrion organelles using *Tiara* and performs organelle-specific gene modeling for CDS, rRNA, and tRNA. Prokaryotic and organelle CDS genes previously modeled with *Prodigal* are now performed more efficiently with *Pyrodigal*.

#### Genome-resolved taxonomic and pathway profiling

Reads-based profiling of shotgun metagenomic and metatranscriptomics have been established as computationally efficient techniques for quantifying taxonomic (e.g., *Sylph* (65), *Kraken 2* (98), *Ganon* (99)) and pathway-level (e.g., *HUMAnN*) abundance and expression, respectively. While small-to-medium sized datasets can be assessed with traditional read alignment tools, this becomes unfeasible with larger numbers of genomes and pathways introducing more genomic redundancies and greater sequence complexity. Despite the computational advantages of reads-based profiling, there are several caveats with the most notable being related to the available databases. More specifically, the findings from these approaches are only as informative as the databases used for profiling allow and most are designed for specific environments. Even in scenarios where the appropriate ecological site is analyzed with respect to the database being used, there is the issue of not knowing which genome the feature is directly associated with in the context of the query samples. The ability to directly associate a specific read with a specific microbe is paramount when studying species with similar strains that occupy different ecological niches (e.g., *Prochlorococcus* ecotypes (100)). While powerful tools exist for building robust custom genome-resolved prokaryotic databases (e.g., *Struo2* (101)) they are not specifically designed to handle eukaryotic organisms. *VEBA 2.0* includes methodologies to build custom databases from genomes (either derived from *VEBA’s* binning modules or externally acquired) that can be used for taxonomic abundance or pathway profiling. In addition to profiling methods, *VEBA 2.0* also provides tools to easily convert counts tables (either generated from profiling or traditional alignment methods) to *biom*, *anndata*, and *pandas pickle* formats.

For taxonomic abundance profiling, *VEBA’s profile-taxonomy* module uses *Sylph* an ultrafast taxonomic profiling method for shotgun metagenomic (and metatranscriptomic) samples by abundance-corrected minhash (65). *Sylph* takes 10x less CPU time and uses 30x less memory than *Kraken2*. Another benefit of using *Sylph* is the ability to customize databases for fragmented genome assemblies and small genomes such as those found within most viruses. *VEBA* uses custom *Sylph* presets designed for viral genomes and non-viral genomes such as prokaryotes and eukaryotes for maximum utility. As with the traditional *mapping* module based on *Bowtie2* alignments, *VEBA* presents an option for aggregating genome-level abundances to SLC-level abundances if clustering results are provided.

For pathway abundance profiling, *VEBA 2.0* uses *HUMAnN* which has been the industry-standard methodology for profiling the abundance of metabolic pathways and other molecular functions. However, as the name suggests, *HUMAnN* databases are developed with human microbiomes as the focus and not environmental samples but the method itself is generalizable. *VEBA* compiles custom *HUMAnN* databases using the genome-resolved proteins, taxonomic classifications, and *UniRef50/90* annotations identified with *VEBA’s* binning, classification, and annotation modules (or acquired elsewhere). *VEBA’s profile-pathway* module not only produces stratified genome-level metabolic pathway abundances but also stratified metabolic pathway completion ratios. As *HUMAnN* operates on single-ended reads, *VEBA* automates read joining via *BBSuite’s bbmerge.sh* program if paired-ended reads or BAM files are used as input. The module also produces a full accounting of reads that align to the custom *HUMAnN* database via *Diamond* and fasta files for reads that do not align.

#### Expanded protein annotation database

*VEBA 1.0* heavily relied on NCBI’s *nr* database. While *nr* is extremely comprehensive, the database is massive on disk (∼200 GB), contains many redundant annotations, and has an inconsistent taxon-specific naming scheme for the protein records. Instead of using *nr* as the base annotation, *VEBA 2.0* uses either *UniRef90* or *UniRef50* for well-characterized and under characterized systems, respectively. *VEBA 2.0* has retained the *Pfam* and *KOFAM* annotations via *HMMER3* and *KofamScan*, respectively; although, these may be replaced with *MMseqs2* in future versions. *VEBA 2.0* also includes several additional databases for protein annotation: 1) *MIBiG* for secondary metabolite synthesis; 2) *VFDB* for virulence factors; 3) *CAZy* for catalytic/carbohydrate metabolism; 4) *NCBIfam-AMR* for antimicrobial resistance; and 5) *AntiFam* for candidate spurious gene calls.

*VEBA 2.0* provides support for identifying either an entire protein database or clustered representatives of a protein database which can be useful for large and complex datasets. With a more uniform annotation syntax using *UniRef* instead of *nr*, *VEBA* 2.0 provides more interpretable consensus annotations for SSPCs. These consensus annotations are useful for assessing the full functional space of protein domains that are within a protein cluster of a pangenome. Annotating SSPCs is much faster than annotating each protein without a significant decrease in information content. For example, consider the 41, 971 SSPCs identified in the reanalysis of the marine eukaryotic organisms from the original *VEBA* publication (described below). On average for each SSPC, 99% of the *Pfam* domains within all proteins of the cluster were detected in the representative of the cluster. Less than 0.5% of the SSPC representatives were missing *Pfam* domains that were exclusive to one of the proteins within the cluster.

As *KEGG* orthology is computed by *KofamScan*, *VEBA* now provides a customized version of *MicrobeAnnotator’s ko_mapper.py* module called *module_completion_ratios.py* that automates the calculation of *KEGG* module completion ratios for a genome (and pangenome if clustering results are provided). This customized script leverages the strengths of *MicrobeAnnotator* while tailoring the input and output to synergize with *VEBA*. The module completion ratio scripts can be used externally without the need for running the full annotation module allowing flexibility for broad usage.

#### Identification and classification of mobile genetic elements

*VEBA 2.0* introduces substantial improvements in the identification and classification of mobile genetic elements such viruses and plasmids; namely, *geNomad* which is the new default backend algorithm. *geNomad* is a classification and annotation framework that combines information from gene content and a deep neural network to identify sequences of plasmids and viruses while using more than 200, 000 marker protein profiles to provide functional gene annotation and taxonomic assignment (42). However, to retain similar functionality to previous versions *VEBA* continues to support *VirFinder* but classification is performed with *geNomad* regardless of the identification algorithm. The taxonomy for *geNomad* uses the most recent revision of viral taxonomy nomenclature ratified by *International Committee on Taxonomy of Viruses* (102) which is currently not yet supported with *vContact2* (103). However, the *vContact* developers are currently working on an updated version which is likely to include the recent change in viral taxonomy nomenclature and may be included as an option in future updates. Lastly, there have also been advances in *CheckV* which have improved the ability to quality assess viruses.

#### Native support for candidate phyla radiation quality assessment and memory-efficient genome classification

*VEBA 1.0* relied on *CheckM* for prokaryotic quality assessment as this was industry-standard at the time of release. Although, *CheckM* could not natively handle Candidate Phyla Radiation (CPR) organisms it contained a *post hoc* workflow for using custom marker sets to correct the quality for CPR. *VEBA 1.0* automated this procedure to abstract away the involved workflow required to correctly assess quality on CPR organisms but it required running *GTDB-Tk* in the backend, identifying CPR, running *CheckM* separately, and updating the existing tables which required considerably more compute time and memory allocation per sample processed (∼128GB at the time). With the release of *CheckM2*, CPR quality assessment is handled natively and bypasses the need to run *GTDB-Tk* or rerun quality assessment *post hoc*. This also drops the compute time and memory allocation substantially (∼16GB per sample). Further, with iterative binning, poor quality MAGs are removed and added back to the grouping of unbinned contigs for the next round. As *GTDB-Tk* runs were computationally expensive, the CPR adjustment was only implemented after the last iteration which made it possible to either include poor quality MAGs that could have been rebinned with higher confidence if the proper quality assessments were determined at each iteration; *CheckM2* bypasses this edge case.

There have also been major improvements in *GTDB-Tk* that have lowered the resource requirements substantially while including more comprehensive reference databases. *VEBA 1.0* used *GTDB-Tk v1.7.0* with *GTDB vR202* (∼128GB memory) while the *VEBA 2.0* uses *GTDB-Tk v2.3.x* with *GTDB vR214.1* (∼72GB memory) with faster runtime. The newer version of *GTDB-Tk* also supports ANI screening using a *mash* database but this is not officially precompiled nor available with the installation or database download. Using an ANI prescreen can reduce computation by more than 50% depending on whether the set of input genomes have a high scoring representative in the database. In addition, the ANI prescreen reduces the number of genomes that need to be classified by *pplacer* which reduces computation time substantially (up to 60%). *VEBA* 2.0 provides an unofficial mirror for the mash build of the *GTDB vR214.1* database (https://zenodo.org/records/8048187) and uses this by default in the backend without any user intervention allowing for effortless access to the most cutting edge prokaryotic classifications.

#### Standalone support for generalized multi-split binning

In 2021, a novel deep learning algorithm called *VAMB* was introduced for using variational auto encoder models for binning genomes from metagenomic assemblies (104). While *VAMB* is not currently supported by *VEBA* due to dependency conflicts with existing packages, *VAMB* introduced an intuitive new approach for binning which they refer to as “multi-split binning”. In multi-split binning, sample-specific assemblies are concatenated but information regarding the samples of origin are retained. Since *VAMB* is not reliant on marker genes (e.g., *MaxBin2*), these contigs can be binned together allowing for more data available for modeling by the neural networks. Once the multi-sample binning is completed, the bins are partitioned according to their samples. While *VAMB* is the first to implement this simple yet powerful approach, the methodology isn’t requisite to *VAMB* and can be generalized. To empower researchers with this approach, we have provided an option in our *binning_wrapper.py* that provides this functionality for binning algorithms that do not require marker sets such as *Metabat2* and *CONCOCT.* We constructed a style guide for binning algorithms implemented in the *binning_wrapper.py*, which is used in the backend of the prokaryotic and eukaryotic binning modules. This technology is designed to be adaptable to handle new binning algorithms if they do not require optimizing marker gene sets for each bin.

#### Automated phylogenomic functional category feature engineering support

Amalgamations are a compositionally-valid dimensionality reduction technique that aggregate low-level features into engineered features using either data-driven approaches (105) or user-specified categories (106). The *PhyloGenomic Functional Category* (PGFC) is a special case of amalgamations designed specifically for microbiomes where counts from low-level features (e.g., SSPCs) are aggregated with respect to a taxonomic category (SLC) and a functional category (KEGG module) producing outputs similar to *HUMAnN* (17). These composite features can be used for downstream statistical analysis and can unpacked back to original features (e.g., SSPCs) unlike other dimensionality reduction methods such as PC[o]A, *t*-SNE, or UMAP. PGFCs are built using the *EnsembleNetworkX* Python package (72) via the *CategoricalEngineeredFeature*class. *VEBA* 2.0 provides a script *compile_phylogenomic_functional_categories.py* which builds PGFCs and genome-level (or SLC-level) MCRs with respect to each sample. Since *VEBA* calculates MCRs for each PGFCs on a per sample basis, they can be used for quality assessment. For example, as it is standard practice to filter low-prevalence compositional features (73), low-prevalence PGFCs can be filtered both by their counts and by MCR thresholds (e.g., MCR < 50%). This functionality provides users with additional approaches for deriving meaning for large and complex datasets.

#### Visualizations of hierarchical data and phylogenies

While a minor addition, *VEBA* 2.0 automates the visualization of hierarchical data types and phylogenetic trees. For instance, *VEBA* 2.0 builds interactive HTML pie charts in the form of *Krona* graphs (107) for prokaryotic and eukaryotic classifications as well as biosynthetic gene clusters within genomes. In addition, phylogenetic trees are rendered automatically using *ETE3* (108) and saved as PDF documents for easily assessing vectorized dendrograms with the added ability of text searching interactively in PDF viewers. While not a visualization, *VEBA* also provides the *NetworkX* graph objects for genome clustering which can be used to produce constellation plots showing each node as a genome and each connection as the ANI connecting the genomes.

### Case studies

#### Revisiting marine phytoplankton case studies from biotechnological and public health perspectives

The *VEBA 1.0* publication analyzed several datasets as case studies to showcase the capabilities. One of these case studies was the *Plastisphere* microbiome (BioProject: PRJNA777294, N = 44 metagenomic samples, 237 gigabases) dataset which included environmental microbial communities from early and mature stage biofilms formed on macroplastics in a marine environment (109). Another case study included the *MarineAerosol* microbiome (BioProject: PRJEB20421, N = 64 metagenomic samples, 90 gigabases) dataset which investigated ocean–atmosphere aerosolization mesocosms and included environmental microbial communities in ocean water collected before, during, and after an algal bloom using the *Wave Flume* ocean simulator (110).

Plastics include a wide range of synthetic or semi-synthetic organic compounds that are durable, malleable, and cheap to manufacture. While the combination of these traits led to accelerated human advancement, the widespread production and distribution of plastics has simultaneously caused public health and environmental crises. The durability property that makes plastic an engineering marvel allows plastics to circulate through the ocean over the course of hundreds to thousands of years before degrading (111). Plastic represents ∼80% of ocean debris (112) and between 4.8 – 12.7 million metric tons are predicted to be deposited into the ocean every year (113). The high deposition and slow degradation rates have caused an accumulation of plastic in the world’s oceans that is projected to exceed 150 million tons by 2025 (113, 114).

From an environmental perspective, plastic pollution threatens marine life across the trophic levels and accumulates in large mid-ocean gyres due to ocean currents. Plastics degrade into micro- and nano-plastic particles containing chemicals that can enter the tissues of marine organisms, including species consumed by humans (115). Further, previous research has shown that microplastics can transmit protozoan pathogens (116), induce reproductive toxicity (117), and have been identified across sensitive regions of the human body including human waste (118) and the placenta (119).

With *VEBA 1.0,* we were able to recover 5 eukaryotic genomes from the *Plastisphere* (early and mature-stage biofilms) and 3 eukaryotic genomes from the *MarineAerosol* datasets (epipelagic and sea-surface microlayers) which were not detected in the original studies (Table 3). All the eukaryotic genomes were medium-to-high quality (*BUSCO* completion ≥ 50% & contamination < 10%). While the backend eukaryotic binning algorithms of *VEBA* have not changed since the initial release, *VEBA* 2.0 has introduced more comprehensive gene modeling databases, more memory efficient gene modeling parameters as defaults, non-coding RNA detection, and organelle detection. The objective for reanalyzing these eukaryotic genomes was to demonstrate the increase in information gain per genome and how this new information can be used for bioprospecting and public health assessments.

**Table 3.** Genome stats for case study 1.

| Genome | Source | Taxonomy Classification | CDS<br>MicroEuk_v2 | CDS<br>MicroEuk50 | CDS<br>MicroEuk90 | CDS<br>MicroEuk100 |
| --- | --- | --- | --- | --- | --- | --- |
| ERR2002407__METABAT2__E.1__bin.2 | Epipelagic Layer | c__Coscinodiscophyceae;<br>o__Thalassiosirales;<br>f__Thalassiosiraceae;<br>g__Stephanocyclus;<br>s__Stephanocyclus meneghinianus | 15655 | 16478 | 16755 | 17098 |
| ERR2002416__METABAT2__E.1__bin.1 | Epipelagic Layer | c__Coscinodiscophyceae;<br>o__Thalassiosirales;<br>f__Thalassiosiraceae;<br>g__Stephanocyclus;<br>s__Stephanocyclus meneghinianus | 17176 | 18258 | 18518 | 18932 |
| ERR2002419__METABAT2__E.1__bin.2 | Sea-Surface Microlayer | c__Coscinodiscophyceae;<br>o__Thalassiosirales;<br>f__Thalassiosiraceae;<br>g__Stephanocyclus;<br>s__Stephanocyclus meneghinianus | 17127 | 18192 | 18459 | 18843 |
| SRR17458614__METABAT2__E.1__bin.2 | Mature Plastic Biofilm | c__Bacillariophyceae;<br>o__Naviculales;<br>f__Phaeodactylaceae;<br>g__Phaeodactylum;<br>s__Phaeodactylum tricornutum | 11475 | 12196 | 12249 | 12384 |
| SRR17458615__METABAT2__E.1__bin.2 | Early Plastic Biofilm | c__Bacillariophyceae;<br>o__Naviculales;<br>f__Phaeodactylaceae;<br>g__Phaeodactylum;<br>s__Phaeodactylum tricornutum | 11434 | 12169 | 12229 | 12362 |
| SRR17458630__METABAT2__E.1__bin.3 | Mature Plastic Biofilm | c__Pelagophyceae;<br>o__Sarcinochrysidales;<br>f__Chrysocystaceae;<br>g__Chrysoreinhardia;<br>s__ | 19827 | 21130 | 21344 | 21647 |
| SRR17458638__METABAT2__E.1__bin.3 | Mature Plastic Biofilm | c__Bacillariophyceae;<br>o__Naviculales;<br>f__Naviculaceae;<br>g__Seminavis;<br>s__Seminavis robusta | 21006 | 23208 | 23336 | 23612 |
| SRR17458638__METABAT2__E.1__bin.2 | Mature Plastic Biofilm | c__Bacillariophyceae;<br>o__Thalassiophysales;<br>f__Catenulaceae;<br>g__Amphora;<br>s__Amphora coffeiformis | 15008 | 15548 | 15691 | 16025 |

The updated *eukaryotic_gene_modeling_wrapper.py* script was run using *MicroEuk100, MicroEuk90, MicroEuk50,* and *MicroEuk_v2*. In all cases, more genes were modeled using the updated protein databases in *VEBA 2.0*. The number of genes detected using the *MicroEuk50* was similar to *MicroEuk90* and *MicroEuk100* with only a fraction of the memory requirements, therefore, *MicroEuk50* is set as the new default with options for using *MicroEuk90* or *MicroEuk100* if users prefer to maximize the gene candidates (Table 3). The updated databases were able to recover on average 6.4%, 7.5%, and 9.2% more genes for *MicroEuk50, MicroEuk90,* and *MicroEuk100*, respectively.

In addition to coding sequences, the gene modeling script identified between 12-42 tRNA and between 0-7 rRNA for the eukaryotic genomes (Table 3, Table S2). The only genome with organelles recovered was a *Stephanocyclus meneghinianus* (*ERR2002419 METABAT2 E.1 bin.2*) which included both a partial mitochondrion and a plastid genome. The previous classification for this genome from *VEBA 1.0* was *Cyclotella meneghiniana* but previous research has confirmed that the genus *Stephanocyclus* includes species that have traditionally been classified under *Cyclotella meneghiniana* (120).

Since plastics contain polymers of carbon atoms with common organic components such as hydrogen, oxygen, nitrogen, and sulphur (121), some organisms have evolved mechanisms to digest these complex forms of carbon (e.g., *Ideonella sakaiensis* hydrolyzed polyethylene terephthalates (PET) and uses the biproducts as building blocks for growth (122)). To investigate the plastic degrading potential of these eukaryotic genomes, protein homologs to *PlasticDB* (123) were assessed. Of the 140, 903 protein coding genes modeled, 156 proteins had plastic degrading properties with the majority representing protease, PEG aldehyde dehydrogenase, hydrolase, and PETase enzymes (Fig. 2, Table S3). These candidates are not surprising as other photosynthetic microeukaryotes such as the marine diatom *Navicula pupula* (124) and fresh water algae (*Scenedesmus dimorphus and Uronema africanum* Borge (125)) have been associated with low-density polyethylene plastic degradation. Naturally occurring enzymes could be used as the starting point for crafting powerful, low-cost, and sustainable biotechnological solutions to lessen the impacts of the plastic crisis. Many different solutions are being actively explored such as adding post-translational glycan modifications to increase both activity and thermostability (126) or codon optimization for fast growing model organisms such as *Escherichia coli* (127). There have even been cases of cross-domain bioengineering such as the introduction of a bacterial PHB biosynthesis pathway into the cytosolic compartment of diatom *Phaeodactylum tricornutum* (128). Further, microplastics are known vectors for microbial pathogens (115). Pathogenic microbes have been detected on microplastics and plastic-containing sea surface biofilms in the Baltic Sea (129). In addition, ciliates associated with coral disease (130) and algae associated with harmful algal blooms (131) have been found hitchhiking on oceanic microplastics.

**Figure 2.**
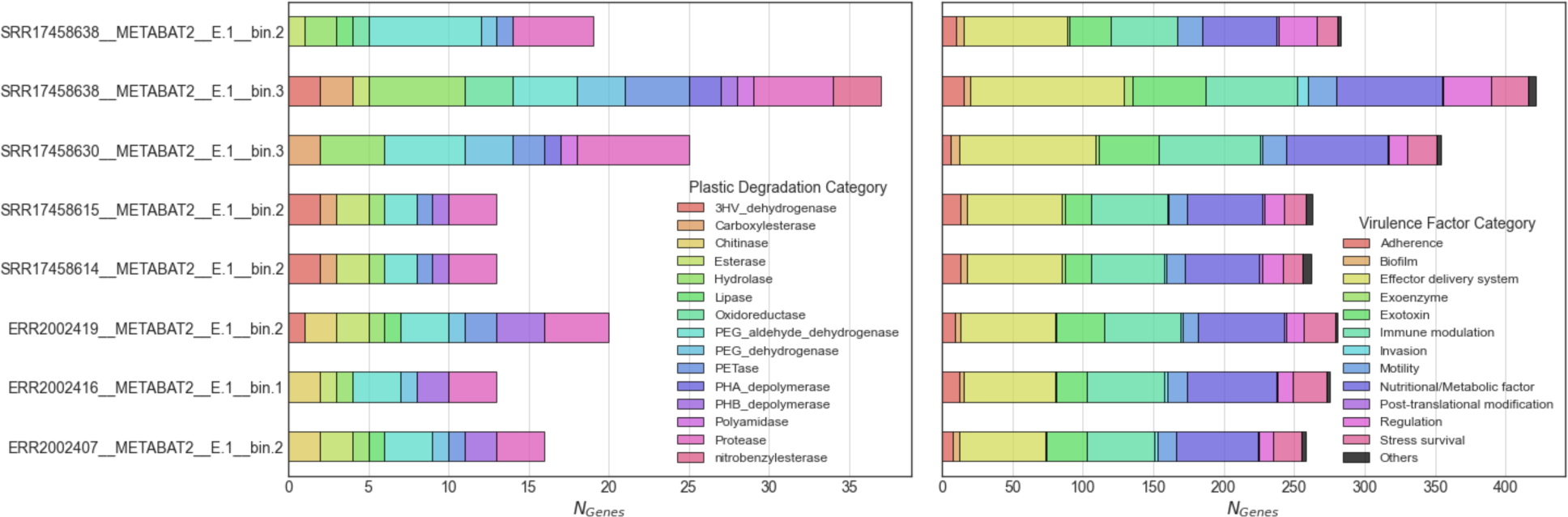
Revisiting microeukaryotic gene calls from marine plastisphere and aerosols (Left) Number of plastic degrading enzymes and (Right) antimicrobial resistance genes.

To assess the pathogenicity of these organisms, protein homologs to *NCBIfam-AMR* (60) and *VFDB* (56) were identified (Fig. 2, Table S3). While *VFDB* is designed for bacterial virulence factors, querying with microeukaryotic phytoplankton proteomes is appropriate as horizontal gene transfer from bacteria to protists has been previously characterized (132). Only two genomes contained AMR genes and both of which were ABC-F type ribosomal protection proteins (*ribo_prot_ABC_F*). Antimicrobial resistant ABC-F family of proteins mediate resistance to many antibiotic classes that bind to the 50S subunit of the ribosome including ketolides (133), lincosamides (134, 135), macrolides (136), oxazolidinones (137), phenicols (137), pleuromutilins (138), and streptogramins of groups A (135, 139) and B (136). Previous research has demonstrated that the ABC-F family can protect the bacterial ribosome from antibiotic-mediated inhibition (140). ABC-F family proteins have been observed in microeukaryotic phytoplankton including the diatom *Fragilariopsis* cylindrus and haptophyte *Emiliania huxleyi* which have 30 and 26 ABC-F proteins, respectively (141).

The most prominent virulence factors included effector delivery systems, nutritional/metabolic factors, and immune modulation. Within *VFDB*, these categories can be partitioned into subcategories: 1) effector delivery systems consisting of Type I-VII secretion systems; 2) nutritional/metabolic factors consisting of metal uptake and metabolic adaptation (e.g., nutrient uptake); and 3) immune modulation consisting of antiphagocytosis, serum resistance, immunoglobulin, antigen variation, apoptosis, and inflammatory signaling pathways (56). While not the most prominent virulence factor, biofilm formation and quorum sensing factors were detected, which play a key role in colonizing marine plastics. As biofilms mature into complex structures, they produce secondary metabolites for quorum sensing and antimicrobial activity for competing microbes (109). While categorized broadly as virulence factors in *VFDB*, anti-inflammatory factors can be used in biomedical research which can drive the translational aspect of ecological research and conservation. Anti-inflammatory properties have been characterized previously in marine phytoplanketon including diatoms (e.g., *Porosira glacialis*, *Attheya longicornis* (142), *Cylindrotheca closterium*, *Odontella mobiliensis*, *Pseudonitzschia pseudodelicatissima* (143) and *Phaeodactylum tricornutum* (144)), dinoflagellates (e.g., *Amphidinium carterae* (144)), and green algae (e.g., *Dunaliella bardawil* and *Dunaliella tertiolecta* (145)).

As an additional confirmation regarding virulence factors against a gold-standard curated metabolic assessment from the same study (109), we analyzed the bacterial *Alteromonadaceae* pangenome recovered from the *Plastisphere* in the initial *VEBA 1.0* release (4). *Bos et al., 2023* conducted a comprehensive analysis, as this was the source study, and highlighted the importance of the mannose-sensitive hemagglutinin (MSHA) operon recognizing it as a key factor for intestinal colonization and for early colonization of hydrophobic plastic surfaces. They also determined positive selection for *mshA* alleles suggesting that *mshA* provides a competitive advantage for surface colonization and nutrient acquisition. In our revisit of the *Alteromonadaceae* pangenome, we identified 92 proteins associated with the MSHA pathway including pili biogenesis (*mshG, mshO, mshQ*), outer membrane secretin (mshL), biotic inner membrane (mshJ, mshI), and pili minor prepilin proteins (*mshD, mshB*) (Table S3). With *VEBA 2.0*, we were able to rapidly screen for MSHA pathway components.

In this updated case study, we showcase the increase in information gain per genome for phytoplankton analyzed in previous case studies. Not only are more coding genes modeled using the updated *MicroEuk_v3* database but we also recover rRNA, tRNA, and partial organelles from the same genomes. Further, we demonstrate how the output from the various modules can be analyzed *post hoc* for identifying candidate plastic degrading enzymes and virulence factors.

#### Ancient 20, 000 - 1, 000, 000 year old Siberian permafrost ecology from ecological and biomedical perspectives

The Siberian permafrost microbiome (BioProject: PRJNA596250, *N* = 7 metagenomic samples) is an *Illumina* MiSeq dataset investigating permafrost from the Kolyma-Indigirka Lowland permafrost from 6 depth profiles ranging from −3.5m to −20m below the surface (146). The geological site where samples were collected originates from the late Pleistocene Era (147) and the soil pertaining to these samples are estimated to have been continuously frozen for ∼20, 000 – 1, 000, 000 years (148, 149). Permafrost environments contain unique microbial ecosystems that are currently under threat from climate change (150). Even though these soils remain frozen year-round, often for thousands or millions of years, they nevertheless maintain living populations of microorganisms operating at low metabolic rates that can be revived and grown in a laboratory (151). The premise of the *Sipes et al., 2021* study was to investigate the distribution of microorganisms that persist in this environment and characterize the metabolism associated with long-term energy starvation.

In the original study, 33 MAGs were identified with varying quality including 10 MAGs of medium-to-high quality (completeness ≥ 50% and contamination < 10%) and 8 MAGs of high-quality (completeness ≥ 80% and < 10% contamination); the 8 high-quality MAGs were used for analysis in *Sipes et al., 2021*. As the purpose of this case study is to showcase updated features, *VEBA 2.0* used the recently added *MEGAHIT* (large and complex metagenome preset) support for metagenomic assembly and was able to increase the number of recovered genomes including 33 MAGs of medium-to-high quality and 11 MAGs of high quality (Table 4, Table S2). Of these 33 MAGs recovered by *VEBA*, 7 MAGs were identified after the first iteration which means they would have been discarded by most binning methods even those using consensus methods such as *DAS Tool*. The 33 MAGs clustered into 17 SLCs with 10 SLCs being singletons with only one representative. Of the bacterial MAGs, there are 6 Actinomycetota, 6 Acidobacteriota, 5 Chloroflexota, 4 Atribacterota, 3 Planctomycetota, 2 Spirochaetota, 1 JANLFM01, 1 Desulfobacterota, and 1 Bacteroidota. Of the archael MAGs, there are 3 Thermoplasmatota and 1 Thermoproteota (Table 4). These prokaryotic MAGs contained 98, 135 protein coding genes that clustered into 65, 523 SSPCs.

**Table 4.**
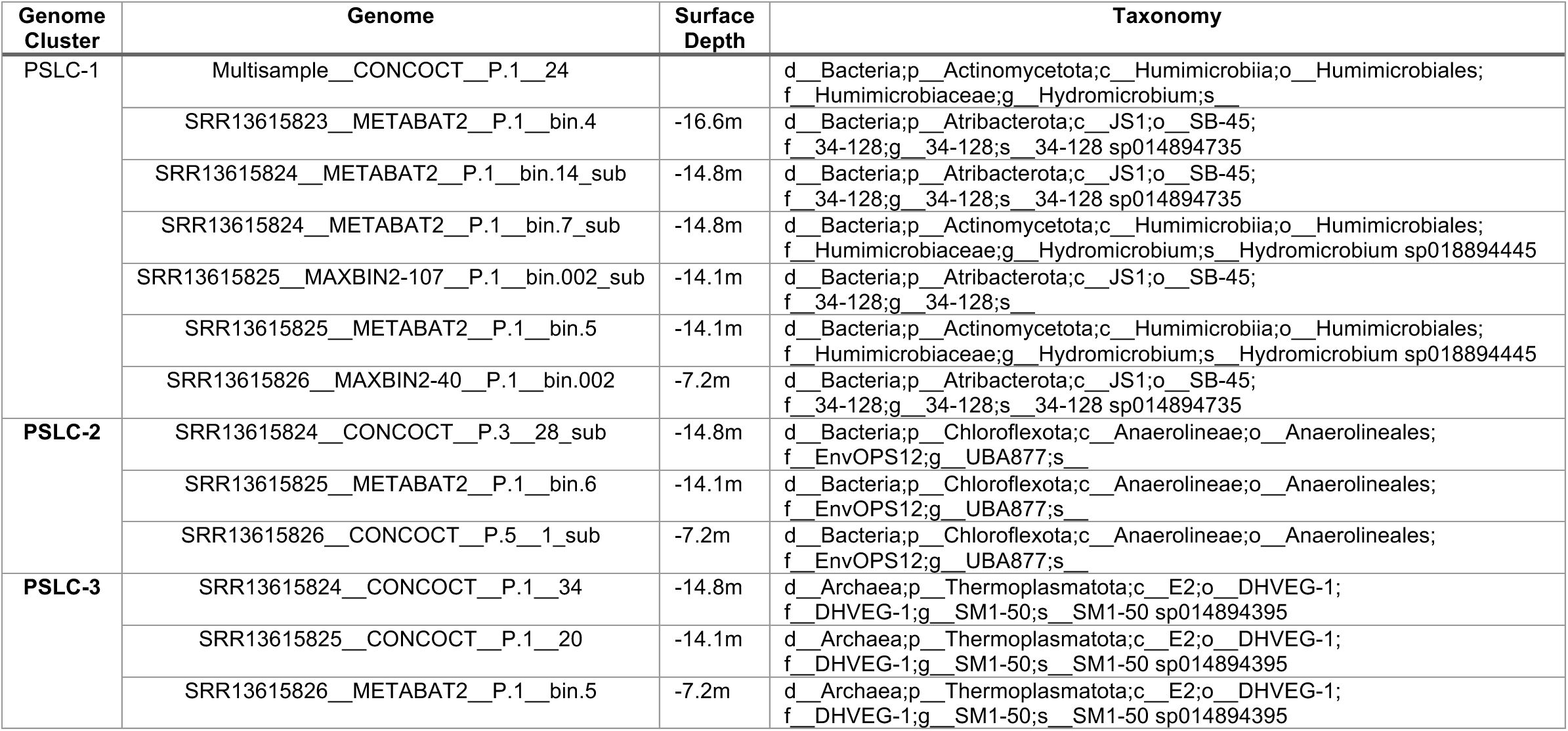
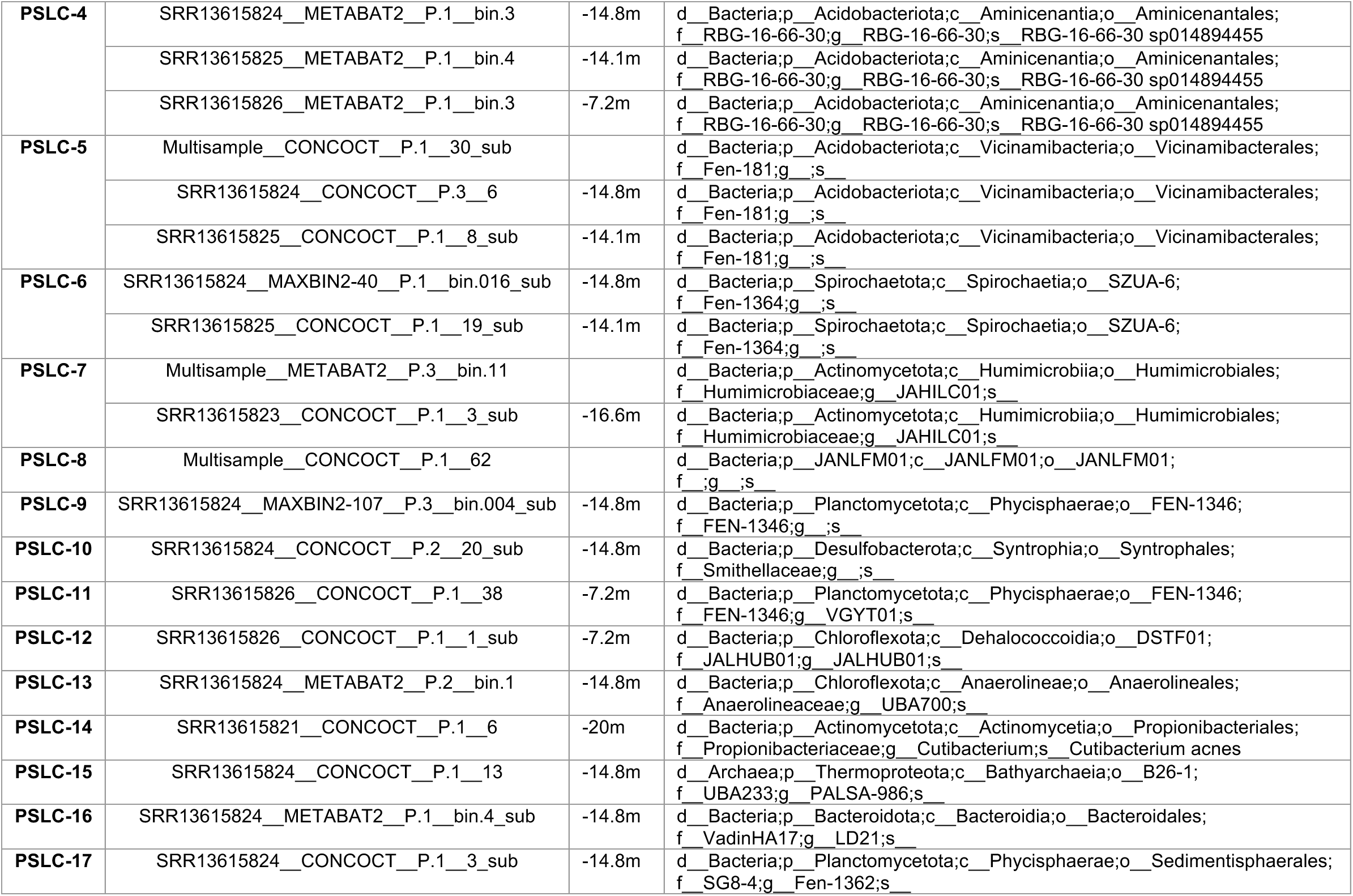
Genome stats for case study 2.

*VEBA* was not able to recover any medium-to-high quality viruses. However, the intermediate step of *geNomad* in the *binning-viral* module identified 925 viral candidates that did not meet the strict criteria *VEBA* uses for defaults (see *Methods* for settings). In addition to viruses, *geNomad* was able to identify 382 plasmids, 141 of which were binned with the prokaryotic MAGs. Large-scale studies have shown that soil viruses are incredibly abundant, highly diverse, and largely uncharacterized (152) with permafrost soils in particular representing a largely understudied genetic resource (153). While *CheckV* is highly robust, poorly-characterized biomes such as permafrost may contain viruses that are not represented in the current reference database. Regarding eukaryotic MAGs, we recovered 1 candidate fungal genome that *BUSCO* determined to be related to Sordariomycetes (completeness 33.3%, contamination 0%) which is not surprising as these fungi have been previously identified in Siberian permafrost (154). Although the candidate Sordariomycetes genome did not meet *VEBA’s* default eukaryotic quality standards (completeness ≥ 50%, contamination < 10%), this level of *BUSCO* completion has been acceptable in *Tara Oceans* protist-centric studies (90, 93); therefore, similar studies may be used to set default thresholds in later *VEBA* releases. These findings suggests that with deeper sequencing one may be able to recover full-length viral and eukaryotic organisms.

To evaluate the biomedical potential of ancient permafrost, we investigated the virulence factors and AMR genes that are now identified automatically with the updated *VEBA annotate* module. PSLC-5 (d Bacteria; p Acidobacteriota; c Vicinamibacteria; o Vicinamibacterales; f Fen-181; g; s) and PSCL-11 (d Bacteria; p Planctomycetota; c Phycisphaerae; o FEN-1346; f FEN-1346; g VGYT01; s) had the highest number of virulence factors per genome. The most prominent virulence factors were related to immune modulation including LPS transporters, daunorubicin resistance, and NAD dependent epimerases (Fig. S1, Table S3). The relevance of immune modulation to biomedical potential was discussed in the previous case study.

Regarding antimicrobial resistance, there were a total of 13 AMR genes from Atribacterota, Chloroflexota, Actinomycetota, and Thermoplasmatota. These genes were mostly involved with mercury metabolism (mercury(II) reductase, mercury resistance co-regulator MerD, mercury resistance system periplasmic binding protein MerP, broad-spectrum mercury transporter MerE), arsenic metabolism (arsenite efflux transporter metallochaperone ArsD, arsinothricin resistance N-acetyltransferase ArsN1 family B), and ABC-F type ribosomal protection (Table S3). Mercury metabolism in permafrost is of potential global importance, mercury is a natural component of soils and high emissions scenarios predict globally significant releases of mercury from thawing permafrost to the atmosphere (155). In addition to these predicted atmospheric emissions, methylmercury - a dissolved form of mercury that can be a toxin in the food web - can be remobilized from thawing permafrost (156). Mercury and arsenic may be remobilized from industrially contaminated permafrost sites (157). Arsenic may also be weathered out of arsenic sulfates in thawing permafrost from which it can enter surface waters (158).

To assess biosynthetic potential of the permafrost microbiome, we ran the *VEBA’s biosynthetic* module and identified 31 BGCs from all SLCs except PSLC-1, PSLC-15, PSLC-6, PSLC-8, and PSLC-9 (Fig. S2, Table S4). The most prominent BGC classes include terpenes, RiPP-like, NRPS/NRPS-like, T3PKS, ranthipeptide, and acyl amino acids but there are many more single occurrence BGC classes as well (Table S4). As the focus of this case study is to showcase the utility of *VEBA* and not to exhaustively explore the genomic landscape, we used a data driven approach to identify which organism to focus on both from an ecological perspective and a biotechnological perspective. In particular, we identified the most highly co-occurring organisms in a community and assessed their potential for producing novel natural products. Using a compositionally-valid network approach, we ascertained that PSLC-3 pangenome (d Archaea;p Thermoplasmatota;c E2;o DHVEG-1;f DHVEG-1;g SM1-50;s SM1-50 sp014894395) was the most highly co-occurring organism in the community despite having average relative abundance (Fig. 3B, C). This archael pangenome consists of 3 genomes from depths −14.8m, −14.1m, and −7.2m and each have their own version of a RiPP-like BGC with the −14.8m and −7.2m strains having novelty scores > 85% based on *MIBiG* homology. RiPP pathways encode a myriad of chemical and functional diversity due to the various modifications added post-translationally to a core peptide via maturase enzymes (159). Each PSLC-3 genome RiPP BGC contained a copy of AsnC family transcriptional regulator and radical SAM domain protein. Radical SAM proteins catalyze diverse reactions including unusual methylations, isomerization, sulfur insertion, ring formation, anaerobic oxidation and protein radical formation while also functioning in DNA precursor, vitamin, cofactor, antibiotic biosynthesis, and in biodegradation pathways (160).

**Figure 3.**
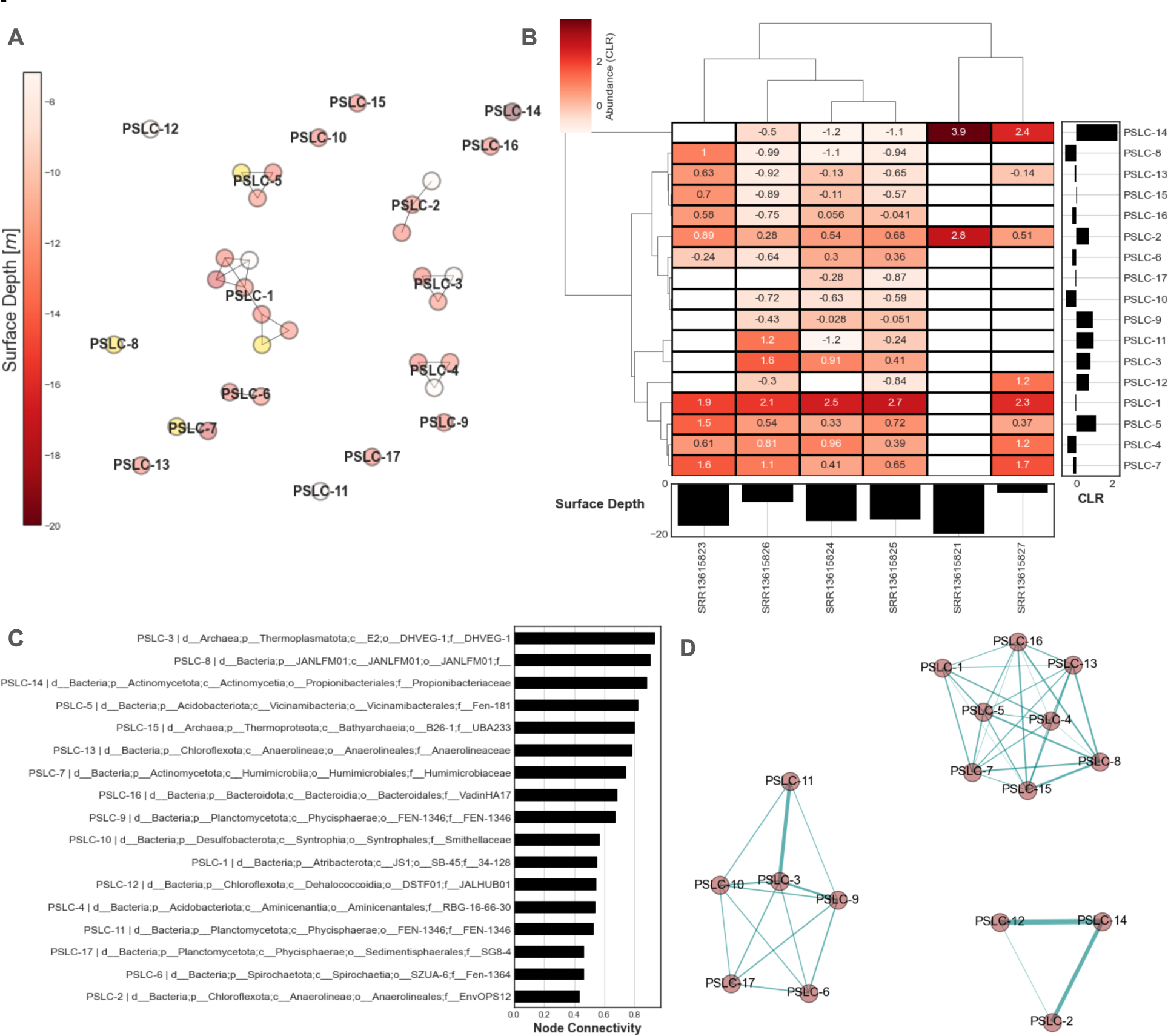
Pangenome analysis of permafrost prokaryotes. (A) Constellation plot of prokaryotic SLCs (yellow indicates multi-sample MAG). (B) Clustermap of transformed SLC abundances with Aitchison distance for samples and pairwise variance log-ratio for SLCs. (C) Graph of Leiden community detection for positive associations calculated from partial correlation with basis shrinkage. (D) Weighted degree shown as node connectivity of Leiden graph.

Recently, RiPP-like natural products have been discovered within the Patescibacteria Eudoremicrobiaceae recovered from the ocean (161). *Paoli et al., 2022* assessed two novel Eudoremicrobiaceae RiPP pathways empirically to provide evidence for their function. In particular, they showed that the deep ocean Eudoremicrobiaceae RiPP-like natural product displayed low-micromolar protease inhibitory activity against neutrophil elastase within a concentration range comparable to other natural products (162) while the second case was predicted to encode a proteusin with unique biochemistry with the first occurrence of a FkbM O-methyltransferase family member in a RiPP pathway. While empirical assessments are out of scope for this case study, the methodology employed by *Paoli et al., 2022* provides the basis and rationale for interrogating natural ecosystems, especially unique systems such as permafrost, for biosynthetic potential and biomedical solutions. It is important to note that all the detected BGCs in the permafrost case study reside on the edge of a contig which means that they are likely incomplete clusters and cannot be fully assessed nor synthesized i*n vitro*. However, these findings show strong evidence that there is biomedically relevant and largely unexplored natural product potential in permafrost which may be unlocked with deeper sequencing using these same *in silico* methods.

For the final *post hoc* analysis for showcasing the utility of genome mining in unique biomes, we screened for CRISPR-Cas systems using *CRISPRCasTyper* (80) which includes classifications for 50 subtypes (163, 164). CRISPR-Cas systems are an adaptive immunity mechanism evolved by prokaryotic organisms in which fragments of foreign DNA (e.g., phage) get stored in the host prokaryotic genome as spacer sequences which are separated by repeat sequences collectively known as a CRISPR array; upon transcription spacer sequences guide *Cas* effector nucleases to destroy the source invader (165). Exploring CRISPR-Cas systems from unique conditions such as permafrost can provide insight into the evolution of CRISPR-Cas systems and those that may perform better in harsh conditions such as those of permafrost.

*CRISPRCasTyper* is not currently implemented in *VEBA* because the current release does not allow for precomputed gene models but this feature is being developed in *CRISPRCasTyper* and will be added to *VEBA* once the update is stable. In the interim, we provide a walkthrough on GitHub to encourage users to screen their datasets for candidate CRISPR-Cas systems.

We recovered 1 high-confidence CRISPR-Cas system (subtype I-E) with *Cas8*, *Cse2*, *Cas7*, *Cas5*, and *Cas6* in SRR13615826 CONCOCT P.1 38 (p Planctomycetota; c Phycisphaerae; o FEN-1346; f FEN-1346; g VGYT01; s VGYT01 sp016872895). In addition, we identified an orphaned high-confidence subtype II-D *Cas* operon with a *Cas9* in SRR13615824 CONCOCT P.3 28_sub (p Chloroflexota; c Anaerolineae; o Anaerolineales; f EnvOPS12; g UBA877; s UBA877 sp017882065) and 2 orphaned high-confidence CRISPR sites including: 1) I-C subtype CRISPR from SRR13615824 CONCOCT P.3 28_sub; and 2) an unknown CRISPR from SRR13615826 CONCOCT P.1 38 (Fig. 4). Previous research has shown that the phylum Chloroflexota comprised 18% (Anaerolineae amounting to 6%) of the total *Cas1* enzymes identified in thermophiles recovered from hot spring metagenomes (166). As of January 2024, the only *Cas9* enzyme from Anaerolineales in NCBI is a partial *Cas9* enzyme from a *Methanoperedens* enrichment culture (CAG0947278.1 from BioSample SAMEA8236570). To date, there are no published *Cas8* in Planctomycetota on *NCBI* or *UniProt* but there are several *Cse2, Cas7, Cas5,* and *Cas6* enzymes. While CRISPR-associated genes from Planctomycetota populate databases (e.g., Cas9 UniProt: A0A954DXG8 (167)) they are often not mentioned or characterized other than *Cas12e* (168). While follow up analysis such as assessing the functional diversity of rare CRISPR-Cas systems (169) or interrogating the spacerome (170) to predict viral source organisms from spacer sequences (171) is out of scope for this study, we showcase the utility of mining extremophile microorganisms for biotechnological potential.

**Figure 4.**
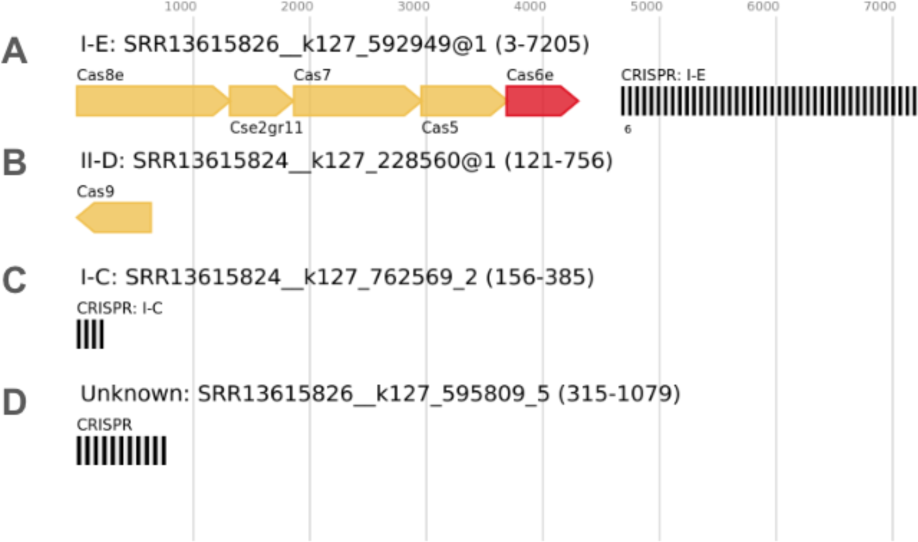
CRISPR-Cas systems in permafrost prokaryotes. (A) CRISPR-Cas system (subtype I-E) with *Cas8*, *Cse2*, *Cas7*, *Cas5*, and *Cas6* in SRR13615826 CONCOCT P.1 38. (B) Orphaned subtype II-D *Cas* operon with a *Cas9* in SRR13615824 CONCOCT P.3 28_sub. (C) Orphaned I-C subtype CRISPR from SRR13615824 CONCOCT P.3 28_sub. (D) Orphaned unknown CRISPR from SRR13615826 CONCOCT P.1 38

Permafrost soils are rapidly thawing, due to climate change (172, 173). This may expose previously sequestered organic matter to microbial degradation to greenhouse gases, further exacerbating climate change (174). Another threat from permafrost thaw is losing habitat for organisms that could potentially have unique biomedical or biotechnological potentials (175). Demonstrating these potentials through bioinformatics can help estimate the scope of potential losses when such habitats are destroyed.

#### Respiratory viruses of white-tailed deer from a veterinary and public health perspective

The white-tailed deer (*Odocoileus virginianus)* case study is an *Oxford Nanopore* dataset derived from lung tissue of pneumonia-related deer fatalities (BioProject: PRJNA1025254, *N* = 34 samples, Prentice et al. in review). While-tailed deer are a species broadly distributed across North America with an estimated population size of ∼30 million animals (176) making it the most abundant large mammal species in North America (177). The distributions of deer and human populations overlap substantially, and is increasing due to changes in human land use and deer habitat range expansion. With increasing human-deer interactions, the circulation of respiratory viruses in white-tailed deer can be a cause of concern as we do not know the zoonotic potential of viruses that are endemic in these species. However, there is documented anthroponosis of SARS-CoV-2 from humans to deer and continued viral transmission within deer populations that increases the potential risk of novel virulent strains emerging back into humans (177). Wildlife reservoirs of broad-host-range viruses have the potential to infect humans or livestock and previous research provides evidence for sustained evolution of SARS-CoV-2 in white-tailed deer and of deer-to-human transmission (178).

Towards better defining the virome of animal species that have dynamic interactions with humans, we showcase the long-read adaptation and viral identification capabilities of *VEBA 2.0*. Here, we analyzed the candidate virome of 17 case samples of deer with pneumonia-related fatalities and 17 control samples. Pneumonia diagnosis and classification was determined by the presence of gross pneumonia lesions and histologic evaluation of lung tissue/lesions as described in *Gilbertson et al., 2022* (179). Only samples from the *Gilbertson et al., 2022* study that were classified as bronchopneumonia or mixed-pneumonia were used in this case study, thus, avoiding cases of interstitial pneumonia.

For *in silico* host depletion we used a concatenated reference including the white-tailed deer reference genome (*Ovir.te_1.0*:GCF_002102435.1) and the human genome (*T2T-CHM13v2.0*:GCF_009914755.1) using the latter to account for potential laboratory contamination. It should be noted that the *Ovir.te_1.*0 genome assembly build was sequenced using short-read *Illumina* sequencing and such assemblies lack structural variants and repeat regions which is relevant to our interpretations.

Using the strict default settings for high-confidence viral genomes (see *Methods* for settings), *VEBA* recovered *N* = 78 viral genomes from case samples and *N* = 77 viral genomes from control samples with genome sizes ranging from 6, 730 – 232, 054 bp (Table 5, Table S2). Candidate viruses recovered in case samples were novel uncharacterized (*N_MAG_* = 38), uncharacterized *Ortervirales* (*N_MAG_* = 35), uncharacterized *Retroviridae* (*N_MAG_* = 3), and *Caudoviricetes* (*N_MAG_* = 2). While in control samples, the types of viruses recovered were uncharacterized *Ortervirales* (*N_MAG_* = 42), novel uncharacterized (*N_MAG_* = 28), and uncharacterized *Retroviridae* (*N_MAG_* = 7). These genomes clustered into 19 SLCs based on 95% ANI threshold with 16 being singletons and the largest cluster VSLC-1 containing ∼84% (130/155) of the viral genomes sourced from ∼85% (29/34) of the samples. These 155 candidate viral genomes contain 2901 proteins that cluster into 1250 SSPCs (Table S3). All viral genome clusters that contained more than 1 genome had representatives in both case and control including VSLC-1 (*N_Case_* = 65 genomes, *N_Control_* = 65 genomes), VSLC-2 (*N_Case_* = 3 genomes, *N_Control_* = 2 genomes), and VSLC-3 (*N_Case_* = 2 genomes, *N_Control_* = 2 genomes).

**Table 5.**
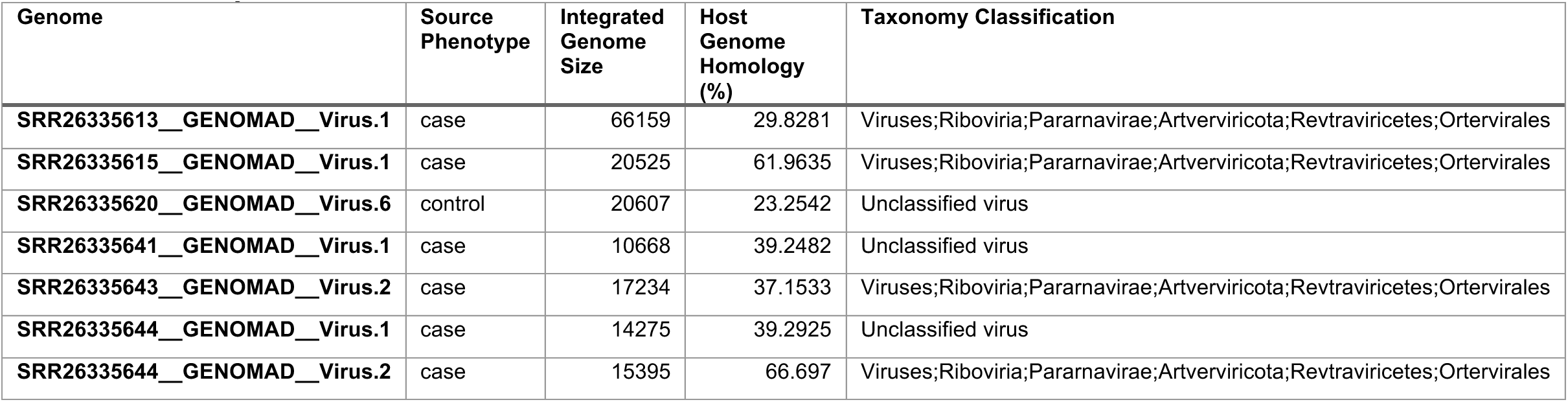
Genome stats for VSLC-1 viral genome cluster with repeat paralogs from case study 3.

While these viruses were classified using robust cutoffs with *geNomad* and were determined to be both high-quality and complete via *CheckV*, with closer examination of the resulting candidate viral genomes (e.g., length, host alignments, and retroviral genes) suggest that these genomes appear to be integrated retroviruses (Table 5). This remains plausible despite the host depletion because individual long reads that align to these genomes span both viral and host genomic content and would not be removed during the decontamination phase which presents the opportunity for identifying these candidate viral integration events. In this case study, we present our rationale for this hypothesis and the evidence to support this largely unexplored area of *in silico* microbiome research.

To place our candidate integrated viral genomes into the context of existing research, we aligned the viral proteins recovered from this case study against viral proteins to white-tailed deer retroviral proteins from NCBI. We identified 79 hits when aligning the viral proteins to known white-tailed deer retroviral proteins, namely: *N* = 36 proteins aligned to endogenous retrovirus group K member 10 Gag polyprotein-like protein (XP_020750633.1); *N* = 36 proteins aligned to endogenous retrovirus group PABLB member 1 Env polyprotein-like protein (XP_020763988.1); and *N* = 7 proteins aligned to endogenous retrovirus group K member 25 Env polyprotein-like protein (XP_020732150.1) (Table S5).

VSLC-1 is classified as an uncharacterized *Ortervirales* and contains no core proteins that are detected in all genomes; the most prevalent is VSLC-1_SSPC-3 which is detected in ∼35% of the genomes. The functional space for this protein cluster has varied activity with most of the annotations suggesting reverse transcription and RNA-directed DNA polymerase but also includes catenin domains for cell adhesion and an ABC-transporter domain. Most of the genomes containing VSLC-1_SSPC-3 have 1 copy (*N_MAG_* = 24) or 2 copies (*N_MAG_* = 13) but some genomes contain up to 8 paralogs. Similar annotations exist for VSLC-1_SSPC-4 which is present in ∼28% of the genomes in VSLC-1.

Another protein cluster worthy of note is VSLC-1_SSPC-1 which is in ∼22% of the VSLC-1 genomes and is the cluster with the largest number of proteins (*N_Proteins_* =144). The number of copies per genome ranges from 1 to as many as 29 paralogs all in the same orientation as in SRR26335613 GENOMAD Virus.1 which has a genome size of 66, 159 bp (Fig. 5, Fig. S3). The only annotation for VSLC-1_SSPC-1 across the entire viral pangenome is a hypothetical protein (UniRef50_UPI001C9E20F9) from *Klebsiella pneumoniae* but the following analysis on VSLC-1_SSPC-1 refers to SRR26335613 GENOMAD Virus.1 unless otherwise noted.

**Figure 5.**
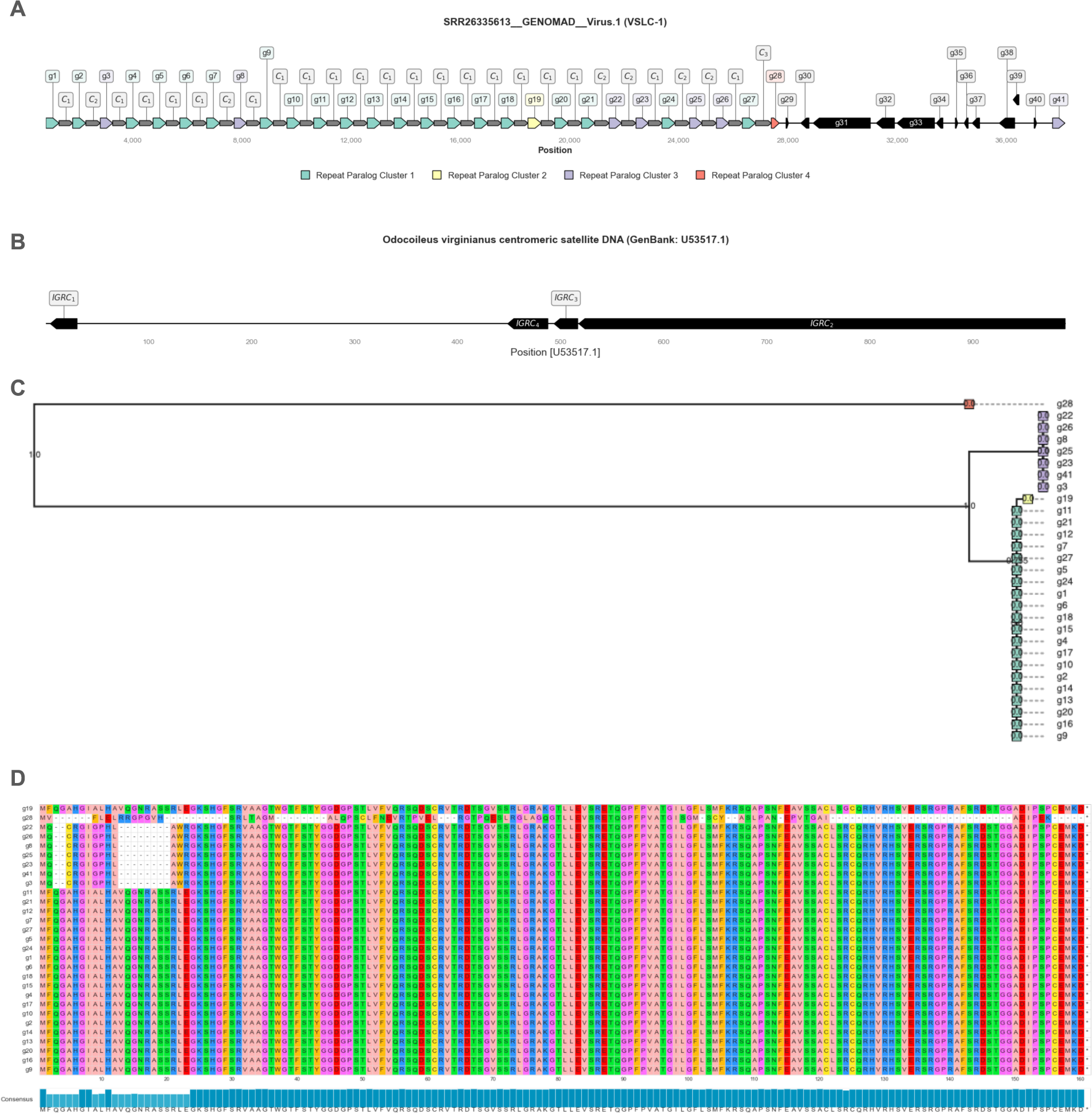
Viral repeat arrays in SRR26335613 GENOMAD Virus.1. (A) Genomic neighborhood plot showing repeat arrays. (B) Intergenic repeated components. (C) Phylogenetic tree showing VSLC-1_SSPC-1 repeat paralogs from SRR26335613 GENOMAD Virus.1 rooted at *g28* to display alignment structure. (D) Multiple sequence alignment of repeat paralogs with leaf ordering from (C).

The first 28/29 paralogs in the repeat array are interspaced by either 497-498bp (*N* = 19), 529bp (*N* = 6), and 577bp for the penultimate intergenic region (Fig. 5A). The final intergenic region between paralogs genes *g28* and *g41* is separated by 10, 010 bp but this section includes viral coding genes. Within the region between the penultimate and last paralog - *g28* and *g41*, respectively - there are 12 genes with robust annotations including several uncharacterized host proteins (UniRef50_A0A6J0X5M2, UniRef50_A0A6J0XDE6), Gag polyproteins (UniRef50_A0A6J0XL79, UniRef50_A0A3G2KVA0), a human endogenous retrovirus K endopeptidase (UniRef50_A0A6J0B691), a zinc ion binding integrase (UniRef50_A0A173DSU7), and several other uncharacterized proteins.

The repeat array includes the following intergenic repeated components (IGRC): IGRC_1_ (27bp intergenic region); IGRC_2_ (470bp intergenic region); IGRC_3_ (30bp intergenic region); and IGRC_4_ (37bp intergenic region) (Fig. 5A, B). All intergenic regions in the main paralog array between genes *g1-g28* begin with IGRC_1_ and this region has 100% identity to *O. virginianus* centromeric satellite DNA reverse complement from position 5-31 denoted as U53517.1-RevComp[5:31]. IGRC_2_ is also in all intergenic regions in the main repeat array region between *g1-g28* and aligns to U53517.1-RevComp[518:990] with 98% identity (1 gap) where a G is deleted in the viral copy relative to the host chromosome. IGRC_3_ is in 7 of the intergenic sequences and aligns with 96% identity to U53517.1-RevComp[494:517]. Lastly, IGRC_4_ is only in the penultimate intergenic region within g27-g28 and aligns with 95% identity (2 gaps) to U53517.1-RevComp[449:488] where AT dinucleotide was deleted in the viral copy relative to the host chromosome. It should be noted that IGRC_2_, IGRC_3_, IGRC_4_ all occur sequentially on the host chromosome but IGRC_1_ is homologous to another region on the other end of the host chromosome. These IGRCs are combined with the viral intergenic regions in three combinations denoted as C_1_ (IGRC_1_ - IGRC_2_), C_2_ (IGRC_1_ - IGRC_2 –_ IGRC_3_); and C_3_ (IGRC_1_ - IGRC_2 –_ IGRC_3 –_ IGRC_4_) (Fig. 5B). Sequences and alignments for these IGRCs are available in the *Supplementary Material*.

Phylogenetic inference reveals that these repeat paralogs cluster into 4 distinct clusters (RPC, Repeat Paralog Clusters) (Fig. 5C, D) with RPC_1_ (*N* = 20), RPC_2_ (*N* = 1), RPC_3_ (*N* = 7), and RPC_4_ (*N* = 1) containing the paralog at end of the main repeat array (Fig. 5A, B). RPC_3_ paralogs always are preceded by C_2_ repeats (Fig. 5A). Further characterization of VSLC-1_SSPC-1 repeat array in this uncharacterized *Ortervirales* viral genome and its effect on white-tailed respiratory disease is outside the scope of this case study. Also, since the data described was generated from genomic DNA and not RNA, the current study may not provide a viral RNA etiology to the respiratory disease found in the cases. However, we encourage researchers to unravel this thread perhaps with defining the viral RNA landscape and deep-learning based structural alignments for exploring functional dark matter.

Although SRR26335613 GENOMAD Virus.1 contains the highest number of paralogs for VSLC-1_SSPC-1, this trend is also observed in other viruses in VSLC-1. To assess whether IGRCs with homology to host centromeres were detected in other viruses we performed exact sequence searches against all the recovered viral genomes and identified the pattern in 6 other viruses (*N* = 7 viruses total, Fig. S4, S5). Of these seven viruses that contain IGRCs, all were members of VSLC-1 and 6/7 were from pneumonia-associated case specimens while only one was from a control specimen (SRR26335620 GENOMAD Virus.6). These six viral genomes recovered from case samples contain different numbers of repeat paralogs that are dispersed at regular intervals. The IGRC-containing virus recovered from a control specimen contains 17 repeat paralogs of VSLC-1_SSPC-1 and does not contain VSLC-1_SSPC-2. A unique feature that was observed in 5/7 viral genomes, exclusive to viruses recovered from case specimens (though, not observed in SRR26335613 GENOMAD Virus.1) was the pattern of VSLC-1_SSPC-2 preceding VSLC-1_SSPC-1 (Fig. S5). There are no annotations for VSLC-1_SSPC-2 from any of the databases used by *VEBA 2.0*. With similar integration patterns at the same centromeric site among several specimens, we speculate that these could be active integrated viruses and not dormant artifacts.

With similar integration patterns at the same centromeric site among several specimens, we speculate that these are viruses that have integrated into the host genome. Integrating into or near centromere regions has been observed with HIV (180) and could be advantageous for an actively infecting virus by ensuring its genome gets copied along with the host chromosome. Investigating the function of these repeated paralogs, intergenic repeats homologous to host centromere satellite DNA, and how the virus integrates into the cell may provide key insight into building stable artificial chromosomes which has been demonstrated in diatoms (181), yeast (182), and mammals (183). The discovery of this repeat array and its implications showcase the utility of *VEBA* and what type of information can be gained only using long-read sequencing technology as these repeat regions are longer than typical *Illumina* reads. *VEBA 2.0* can be used for identifying and annotating integrated viruses in long-read genome assemblies and unlocks numerous potential new studies. For example, within these 34 samples our data suggests the prevalence of integrated viruses across different individual deer and that there is a unique aspect in terms of paralogous repeat structure.

To assess if any functional domains were statistically enriched in viral populations in the case samples relative to control samples or vice versa, we performed Fisher’s exact tests. First, we investigated protein clusters within VSLC-1 as this pangenome contains most of the viral genomes. The only unsupervised VSLC-1 protein-cluster that was statistically enriched in control (*N* = 6 genomes) relative to case (*N* = 0 genomes) was VSLC-1_SSPC-14 (P ≅ 0.01) which has no homology to any databases used by *VEBA 2.0*. From a supervised approach, the only *Pfam* domain that was statistically depleted in control (*N* = 15) relative to case samples (*N* = 28) was PF18697.4 (P ≅ 0.02) which is a murine leukemia virus (MLV) integrase C-terminal domain interacts with the bromo and extra-terminal proteins through the ET domain. This interaction provides a structural basis for global *in vivo* integration-site preferences and disruption of this interaction through truncation mutations affects the global targeting profile of MLV (184).

As protein clustering is performed within SLCs and not globally, we assessed the entire virome from a supervised perspective using *Pfam* domains which yielded 3 enriched protein domains with two being enriched in case relative control samples (Murine leukemia virus integrase C-terminal domain PF18697.4 (N_Case_ = 31, N_Control_ = 16, P ≅ 0.01) and Integrase core domain PF00665.29 (N_Case_ = 37, N_Control_ = 23, P ≅ 0.03)) and one enriched in control relative to case samples (RNase H-like domain found in reverse transcriptase PF17917.4 (N_Case_ = 6, N_Control_ = 15, P ≅ 0.04)). Integrase mediates integration of a DNA copy of the viral genome into the host chromosome which consists of three domains including the central catalytic domain for zinc binding, a non-specific DNA binding domain, and a catalytic domain that acts as an endonuclease (185). Ribonuclease H (RNase H) activities allow reverse transcriptases to convert retroviral ssRNA genome into dsDNA which is integrated into the host genome during infection (186). While it is unclear on how these viral features may contribute to pneumonia-related illness, it is possible that these integrated viruses may be involved in host immune response (187). As previously noted, our current approach of analyzing genomic DNA may not fully reflect the etiology of pneumonia-related events.

There are an estimated 40, 000 viral species circulating in mammals, a quarter of which have zoonotic potential (188); in particular, many of the recent epidemics of emerging respiratory viruses are believed to have originated in wildlife (189, 190). The reverse can also occur where reverse zoonosis or anthroponosis contributes to the diversity of viral species in animals that have high interactions with humans (191). The dynamic relationship between humans and animal species and the associated diseases not only put the persistence of wildlife populations at risk (e.g., bighorn sheep pneumonia (192)) but also pose a risk for transmission to domesticated livestock (193) and humans (178) Thus, these events threaten both public health and food security. Advances in molecular sequencing have been instrumental in facilitating pathogen detection and characterization for microbes including viruses that are difficult to identify using culturing-based diagnostic approaches. The added support for long-read sequencing provided insight into complex genomic features of candidate integrated white-tailed deer viruses including repetitive paralogs and host centromeric satellite incorporation. Further, this case study provides evidence for the discovery and characterization of integrated retroviruses which may play a role in the health of white-tailed deer populations experiencing fatalities attributed to pneumonia (*Prentice et al. in review*) and chronic wasting disease (194).

### Future perspectives

The ability to integrate genomics into research studies has become increasingly routine due to the continuous decline in sequencing costs per megabase – costs that have far exceeded predictions by Moore’s Law (195). The astonishing rate of new of sequencing advancements has led to petabytes of publicly available structured sequence data including 19.6 trillion base pairs from over 2.9 billion nucleotide sequences in *NCBI’s GenBank* database (196) and even more raw sequencing data including over 90.11 quadrillion base pairs in *NCBI’s SRA* database (<u>ncbi.nlm.nih.gov/sra/docs/sragrowth/</u>). Comprehensive software suites such as *VEBA* are a requisite for keeping up with and deriving meaning from this massive amount of sequencing data that is generated and deposited in public repositories each year. Further, as we have demonstrated previously (4) and in this most recent edition, there is potential value to be gained when reanalyzing existing datasets with updated methodologies especially in the context of metagenomics and *in silico* genome mining for bioprospecting.

To ensure that *VEBA* can address the deluge of sequencing data, we plan to add performance improvements, new features as new methods are developed, and increased interoperability with other high-quality workflows under active development such as *DRAM*(197), *Bakta* (198), *MuDoGeR* (199), *EukHeist* (200), and *Anvi’o* (201). In the near future, *VEBA* will be adding a module for CRISPR-Cas screening and historical viral infections prediction from spacer sequences once *CRISPRCasTyper* fully supports preexisting gene calls. In addition, we also plan to add support for addressing functional dark matter (202) to demystify the function of hypothetical proteins using remote homology detection and structural alignment methods (203). In the long term on the path towards *VEBA* 3.0, we plan to reimplement the entire software suite using *Nextflow* to further maximize our performance gains in terms of parallelization capabilities and resource usage. However, this reimplementation is non-trivial as it requires rewriting all the modules and scripts in *Nextflow* and the *Groovy* programming language.

Regarding feature updates, we will add new technologies as they are developed to solve new problems or to update existing solutions. In particular, there is a need for further developments in eukaryotic taxonomy classification as many of the *in silico* methodologies (e.g., *VEBA’s classify-eukaryotic* module and *EUKulele* (204)) rely on protein alignments which are not as flexible as the methods developed for prokaryotic classifications. To address this need, developers with expertise in eukaryotic taxonomy would have to build an *in silico* eukaryotic taxonomic classification tool that is as robust as *GTDB-Tk.* Such improvements in eukaryotic classification methods would illuminate the blind-spots in microbial ecology casted by microeukaryotic organisms and would provide a major milestone for large-scale efforts such as *Earth BioGenome Project* (205, 206).

The need for cataloguing Earth’s biodiversity is time sensitive due to anthropogenic climate change. The authoritative *Intergovernmental Panel on Climate Change* (IPCC) recently concluded that human activities have unequivocally warmed the planet while causing substantial biodiversity loss associated with downstream effects such as desertification, decreased precipitation, land/forest degradation, glacial retreat, ocean acidification, and sea level rise (207). The loss of biodiversity is expected to alter ecosystem functionality and the ability to provide society with the requisite resources to prosper (208). Further, with a decrease in biodiversity we also forfeit the potential for discovering natural products with biomedical relevance such as antimicrobial (209–211), chemotherapy (212), and antiviral agents (213). With biodiversity loss growing increasingly dire (214), curbing anthropogenic biodiversity loss is paramount and requires policies to address the multifaceted crisis (215). However, basic research has not been enough to drive substantial change in policies needed to prevent such catastrophes. Translating basic research findings into cogent policy and providing candidate assets for biotechnology industry will not only instill growing interest in basic research but will also provide an economic incentive to preserve natural systems in the prospect of identifying novel natural products from high biodiversity regions.

With the advent of comprehensive software suites such as *VEBA*, further developments for *in silico* bioprospecting/screening methods and continued advancements in sequencing technologies, the future of environmental/biodiversity preservation may be driven not just from ethical concerns but also from biotechnological potential for discovering solutions developed by nature. That is, it is essential to preserve ecosystems in an effort to not diminish the catalog of candidate natural products before they can be discovered. Bridging the gap between environmental sustainability and translational potential may be the requisite change needed for policy changes that can dampen the twin crisis of climate change and biodiversity collapse (216).

## Data Availability

The source code for *VEBA* (https://github.com/jolespin/veba), the clustered *MicroEuk* database (https://zenodo.org/records/10139451), and the analysis for case studies (https://zenodo.org/records/10780433) are publicly available. The original studies for the Plastisphere microbiome (BioProject: PRJNA777294), MarineAerosol microbiome (BioProject: PRJEB20421), Siberian permafrost microbiome (BioProject: PRJNA596250), and white-tailed deer lung microbiome (BioProject: PRJNA1025254) are available on NCBI (https://www.ncbi.nlm.nih.gov/bioproject/).

## Funding

This work was funded by the following sources: 1) NIH R21AI160098, NIH 1R01AI170111-01, NIH 1U54GH009824, NSF OCE-1558453, and PolyBio Foundation awarded to CLD; 2) NSF Dimensions of Biodiversity (DEB-1442262), and US Department of Energy, Office of Science, Office of Biological and Environmental Research, Genomic Science Program (DE-SC0020369) to KL, Additional funding was provided by the Federal Aid in Wildlife Restoration Act administered by the U.S. Fish and Wildlife Service through the Wisconsin Department of Natural Resources.

## Conflict of Interest Disclosure

Authors declare not competing interest.

## Acknowledgements

We would like to thank our funding sources and our colleagues for support during this research. In addition, we would like to thank Harinder Singh and Yang Chen for testing beta versions of this software.

---

Table S1 – Software dependencies, prevalence across modules, and licenses Table S2 – Summary statistics of genomes and taxonomic classifications Table S3 – Annotations for all protein-coding genes

Table S4 – BGCs for case study 2

Table S5 – Viral proteins from case study 3 homologous to known white-tailed deer retroviral proteins

Supplementary Material – Document containing sequences for SRR26335613 GENOMAD Virus.1 (VSLC-1): i) repeat paralog multiple sequence alignments (MSA); ii) intergenic sequences between repeat paralogs; iii) intergenic repeated components; and iv) intergenic repeated components (MSA).

**Figure S1.**
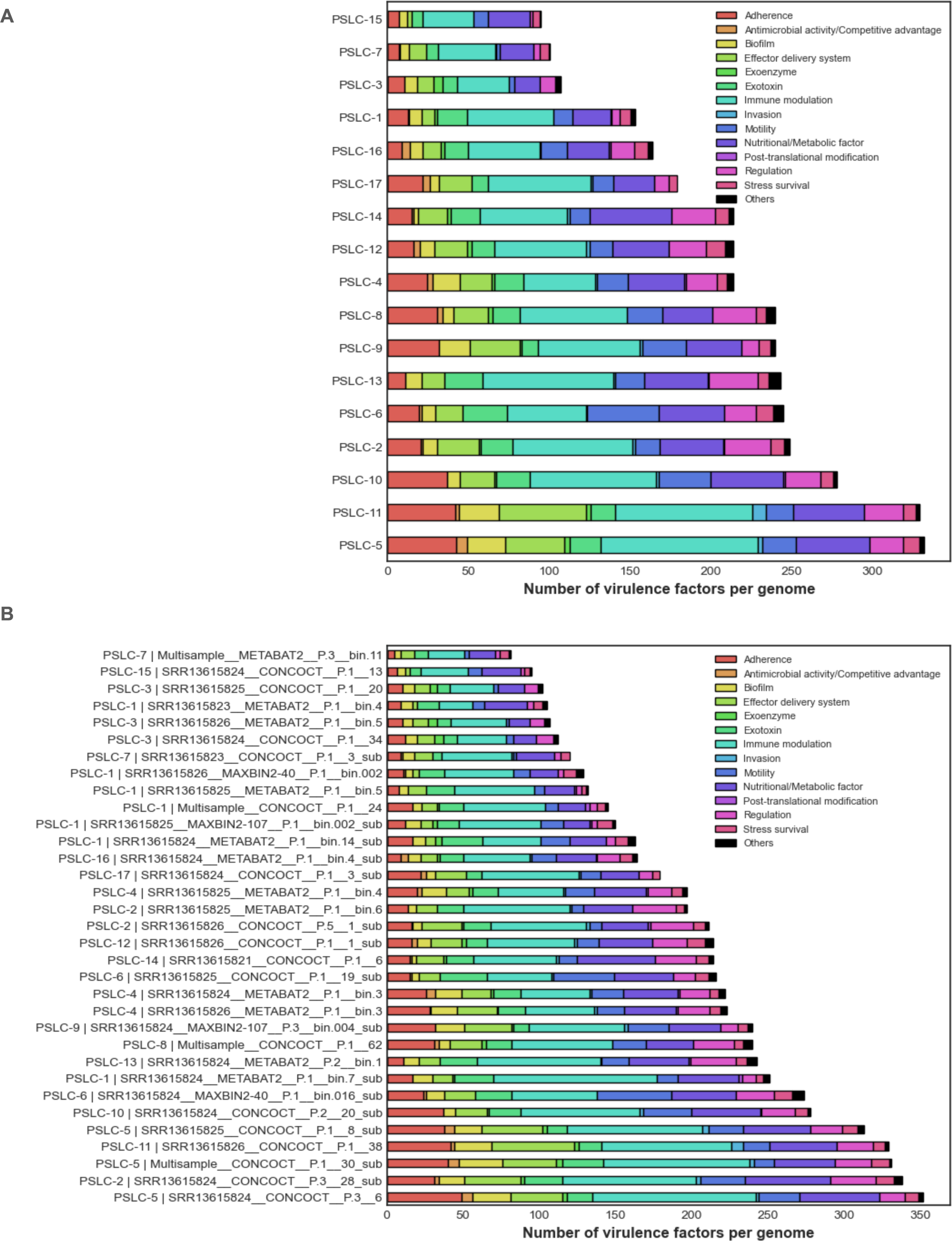
Distribution of virulence factors in permafrost prokaryotes. Number of virulence factor genes grouped by virulence category for (A) each SLC per genome and (B) each genome.

**Figure S2.**
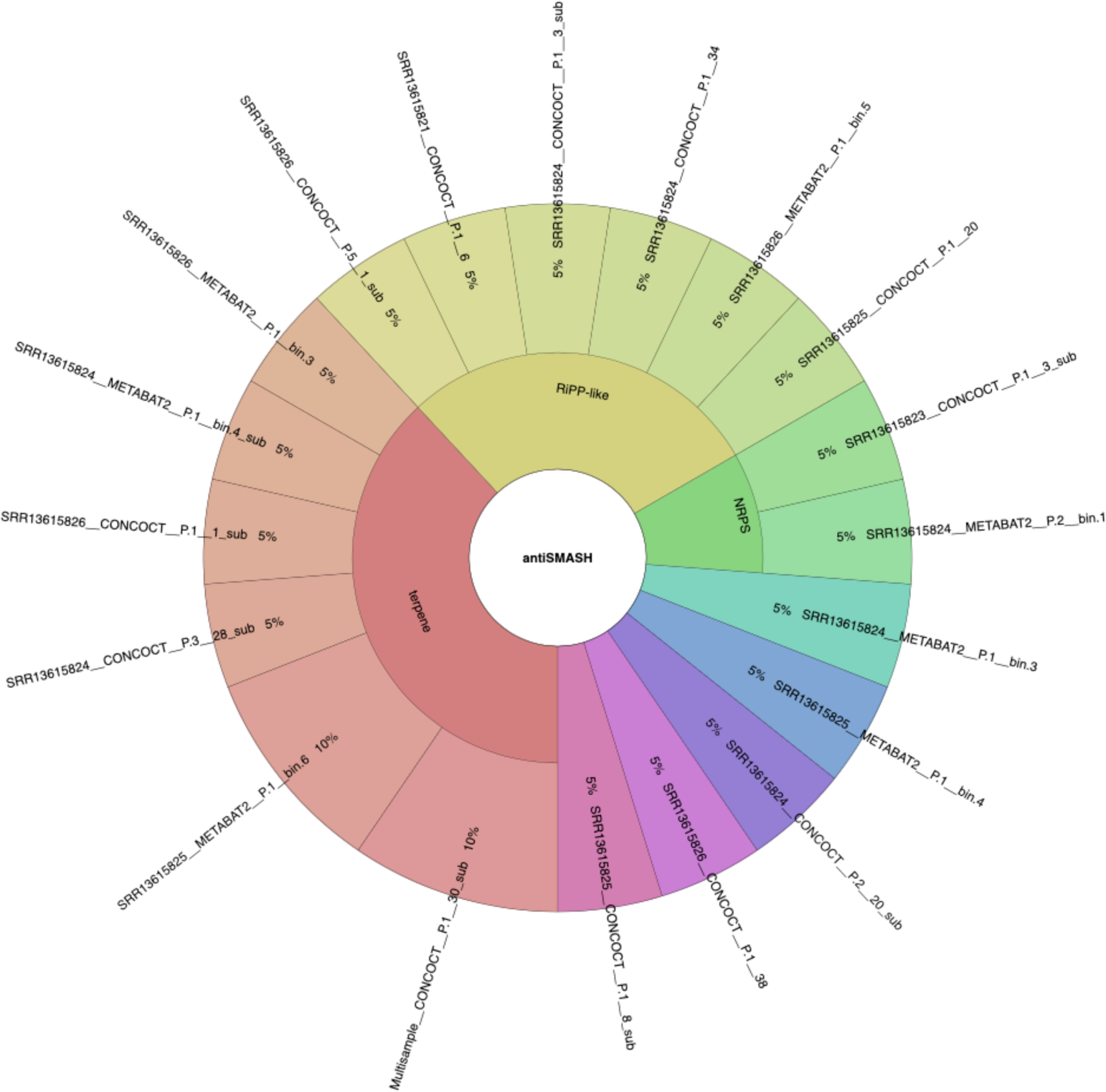
Distribution of biosynthetic gene clusters in permafrost prokaryotes. Distribution of biosynthetic gene clusters identified by *antiSMASH* for each genome.

**Figure S3.**
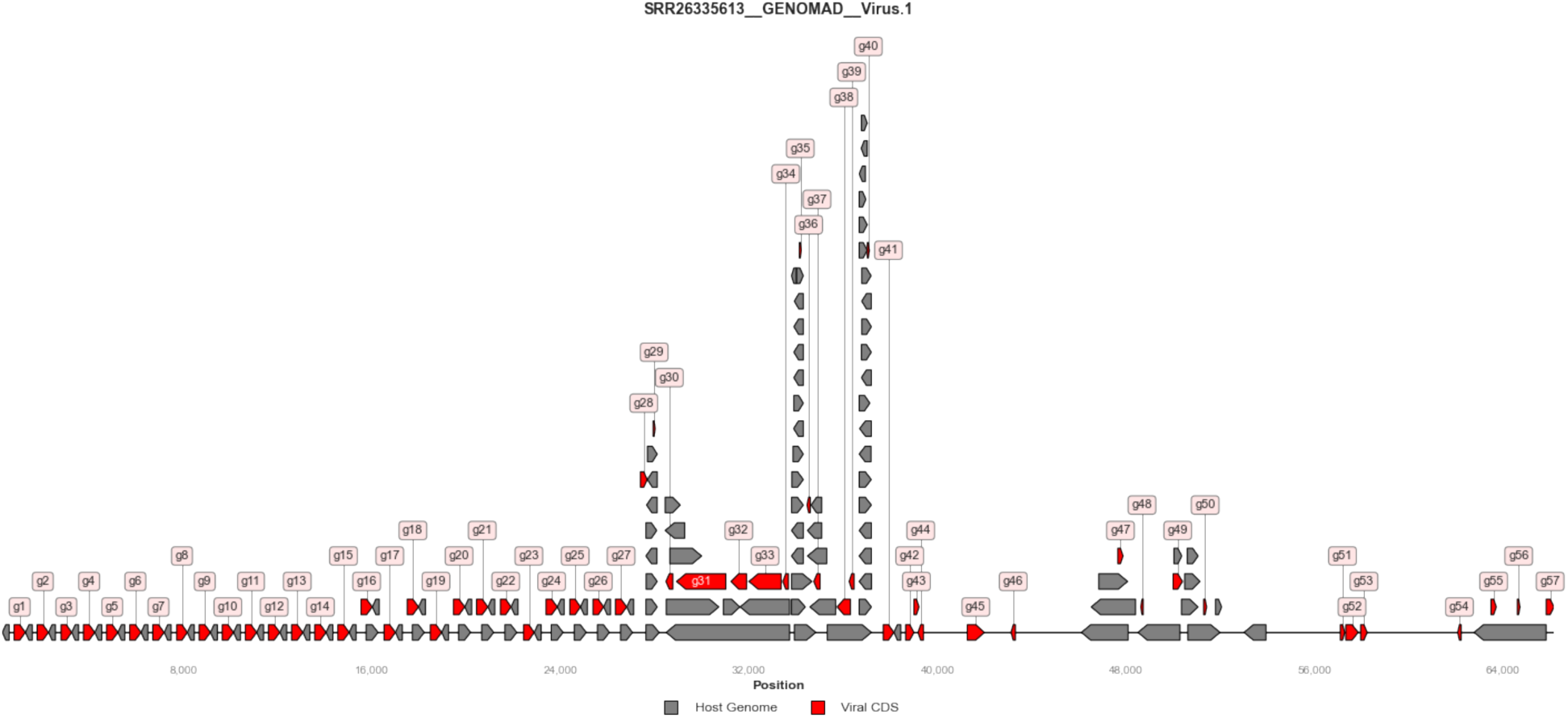
Viral coding sequences and host homology of candidate integrated viral genome. Genomic neighborhood plot showing SRR26335613 GENOMAD Virus.1. Gray indicates homology to host genome and red indicates viral CDS.

**Figure S4.**
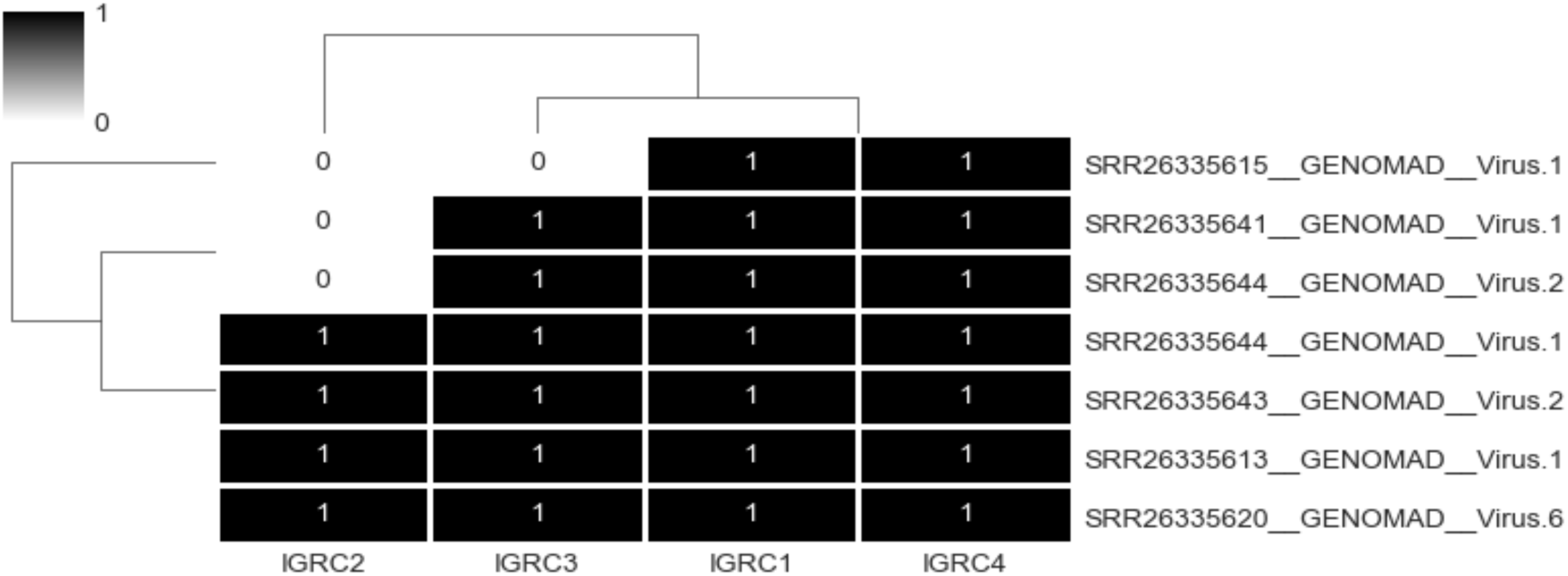
Intergenic repeated components across multiple viruses in VSLC-1. IGRCs detected with 100% sequence identity in viral genomes.

**Figure S5.**
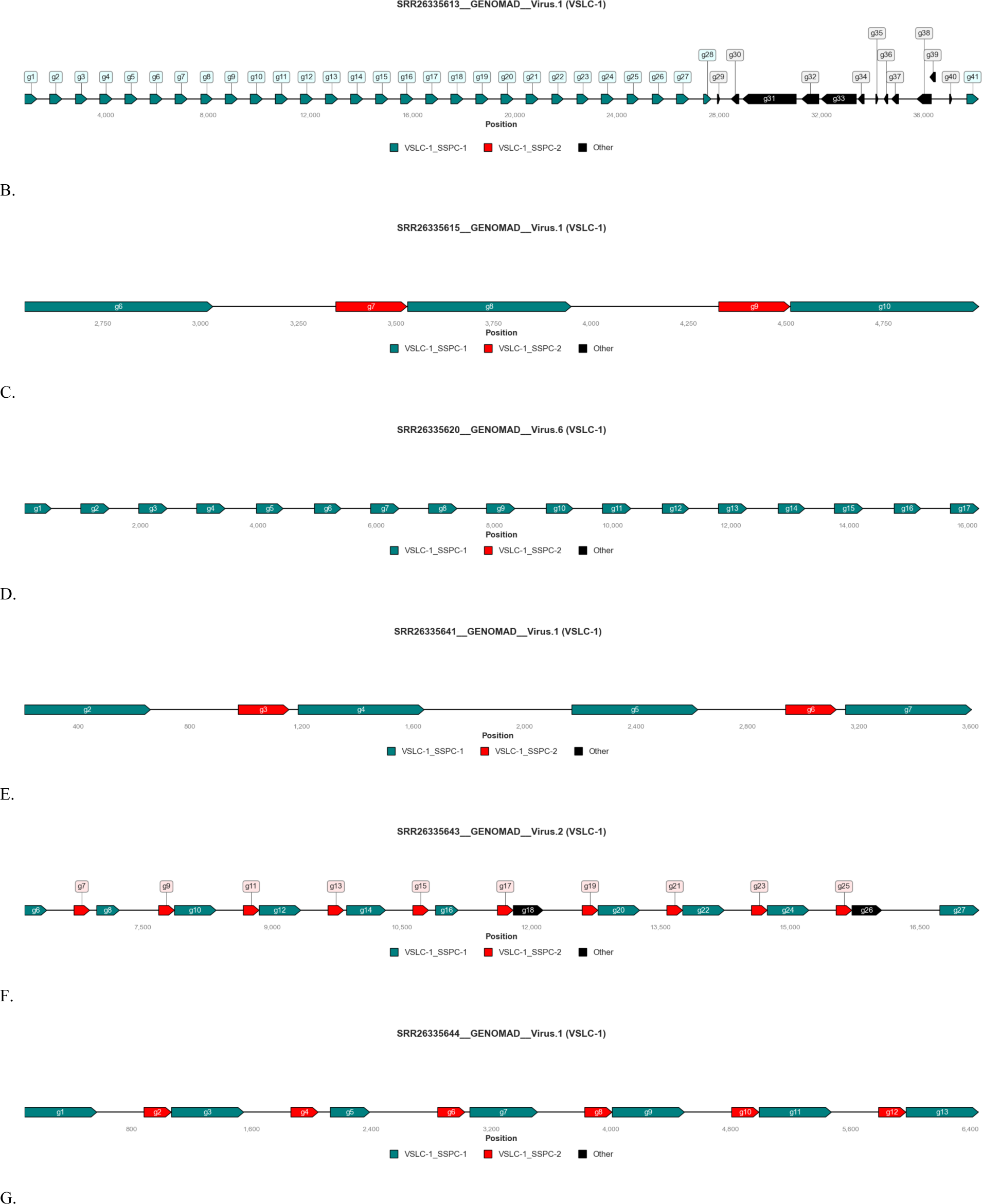
Repeat paralogs across multiple viruses in VSLC-1. Genomic neighborhood plot for viral genomes that contain IGRCs with emphasis on repeat paralogs.

## Notes

https://github.com/jolespin/veba

### Competing Interest Statement

The authors have declared no competing interest.

### Summary of Updates

Error in middle initial on author list.

