## Supplementary Material for "Unveiling the Microbial Realm with VEBA 2.0: A modular bioinformatics suite for end-to-end genome-resolved prokaryotic, (micro)eukaryotic, and viral multi-omics from either short- or long-read sequencing"

#### i. SRR26335613__GENOMAD__Virus.1 (VSLC-1) repeat paralog (MSA)

>g19
MFQGAHGIALHAVQGNRASSRLEGKSHGFSRVAAGTWGTFSTYGGDGPSTLVFVQRSQDSCRVTRDTSGVSSRLGRAKGTLLEVS
RETQGPFPVATGILGFLSMFKRSQAPSNFEAVSSACLSGCQRHVRHSVERSRGPRAFSRDSTGGADIPSPCEMKD*
>g28
MV------FLELRRGPGVH----------SRLTAGM--------ALQPSCLFNEVRTPVEL---
RGTPQESLRGLAGQGTLLEVSRETQGPFPVATGISGM-SCY--ASLPAN-EPVTGAI----------------------------AEIPEK-----*
>g22
MQ--CRGIGPHL--------
AWRGKSHGFSRVAAGTWGTFSTYGGDGPSTLVFVQRSQDSCRVTRDTSGVSSRLGRAKGTLLEVSRETQGPFPVATGILGFLSMFK
RSQAPSNFEAVSSACLSRCQRHVRHSVERSRGPRAFSRDSTGGADIPSPCEMKD*
>g26
MQ--CRGIGPHL--------
AWRGKSHGFSRVAAGTWGTFSTYGGDGPSTLVFVQRSQDSCRVTRDTSGVSSRLGRAKGTLLEVSRETQGPFPVATGILGFLSMFK
RSQAPSNFEAVSSACLSRCQRHVRHSVERSRGPRAFSRDSTGGADIPSPCEMKD*
>g8
MQ--CRGIGPHL--------
AWRGKSHGFSRVAAGTWGTFSTYGGDGPSTLVFVQRSQDSCRVTRDTSGVSSRLGRAKGTLLEVSRETQGPFPVATGILGFLSMFK
RSQAPSNFEAVSSACLSRCQRHVRHSVERSRGPRAFSRDSTGGADIPSPCEMKD*
>g25
MQ--CRGIGPHL--------
AWRGKSHGFSRVAAGTWGTFSTYGGDGPSTLVFVQRSQDSCRVTRDTSGVSSRLGRAKGTLLEVSRETQGPFPVATGILGFLSMFK
RSQAPSNFEAVSSACLSRCQRHVRHSVERSRGPRAFSRDSTGGADIPSPCEMKD*
>g23
MQ--CRGIGPHL--------
AWRGKSHGFSRVAAGTWGTFSTYGGDGPSTLVFVQRSQDSCRVTRDTSGVSSRLGRAKGTLLEVSRETQGPFPVATGILGFLSMFK
RSQAPSNFEAVSSACLSRCQRHVRHSVERSRGPRAFSRDSTGGADIPSPCEMKD*
>g41
MQ--CRGIGPHL--------
AWRGKSHGFSRVAAGTWGTFSTYGGDGPSTLVFVQRSQDSCRVTRDTSGVSSRLGRAKGTLLEVSRETQGPFPVATGILGFLSMFK
RSQAPSNFEAVSSACLSRCQRHVRHSVERSRGPRAFSRDSTGGADIPSPCEMKD*
>g3
MQ--CRGIGPHL--------
AWRGKSHGFSRVAAGTWGTFSTYGGDGPSTLVFVQRSQDSCRVTRDTSGVSSRLGRAKGTLLEVSRETQGPFPVATGILGFLSMFK
RSQAPSNFEAVSSACLSRCQRHVRHSVERSRGPRAFSRDSTGGADIPSPCEMKD*
>g11
MFQGAHGIALHAVQGNRASSRLEGKSHGFSRVAAGTWGTFSTYGGDGPSTLVFVQRSQDSCRVTRDTSGVSSRLGRAKGTLLEVS
RETQGPFPVATGILGFLSMFKRSQAPSNFEAVSSACLSRCQRHVRHSVERSRGPRAFSRDSTGGADIPSPCEMKD*
>g21
MFQGAHGIALHAVQGNRASSRLEGKSHGFSRVAAGTWGTFSTYGGDGPSTLVFVQRSQDSCRVTRDTSGVSSRLGRAKGTLLEVS
RETQGPFPVATGILGFLSMFKRSQAPSNFEAVSSACLSRCQRHVRHSVERSRGPRAFSRDSTGGADIPSPCEMKD*
>g12
MFQGAHGIALHAVQGNRASSRLEGKSHGFSRVAAGTWGTFSTYGGDGPSTLVFVQRSQDSCRVTRDTSGVSSRLGRAKGTLLEVS
RETQGPFPVATGILGFLSMFKRSQAPSNFEAVSSACLSRCQRHVRHSVERSRGPRAFSRDSTGGADIPSPCEMKD*
>g7
MFQGAHGIALHAVQGNRASSRLEGKSHGFSRVAAGTWGTFSTYGGDGPSTLVFVQRSQDSCRVTRDTSGVSSRLGRAKGTLLEVS
RETQGPFPVATGILGFLSMFKRSQAPSNFEAVSSACLSRCQRHVRHSVERSRGPRAFSRDSTGGADIPSPCEMKD*
>g27
MFQGAHGIALHAVQGNRASSRLEGKSHGFSRVAAGTWGTFSTYGGDGPSTLVFVQRSQDSCRVTRDTSGVSSRLGRAKGTLLEVS
RETQGPFPVATGILGFLSMFKRSQAPSNFEAVSSACLSRCQRHVRHSVERSRGPRAFSRDSTGGADIPSPCEMKD*
>g5
MFQGAHGIALHAVQGNRASSRLEGKSHGFSRVAAGTWGTFSTYGGDGPSTLVFVQRSQDSCRVTRDTSGVSSRLGRAKGTLLEVS
RETQGPFPVATGILGFLSMFKRSQAPSNFEAVSSACLSRCQRHVRHSVERSRGPRAFSRDSTGGADIPSPCEMKD*
>g24
MFQGAHGIALHAVQGNRASSRLEGKSHGFSRVAAGTWGTFSTYGGDGPSTLVFVQRSQDSCRVTRDTSGVSSRLGRAKGTLLEVS
RETQGPFPVATGILGFLSMFKRSQAPSNFEAVSSACLSRCQRHVRHSVERSRGPRAFSRDSTGGADIPSPCEMKD*
>g1
MFQGAHGIALHAVQGNRASSRLEGKSHGFSRVAAGTWGTFSTYGGDGPSTLVFVQRSQDSCRVTRDTSGVSSRLGRAKGTLLEVS
RETQGPFPVATGILGFLSMFKRSQAPSNFEAVSSACLSRCQRHVRHSVERSRGPRAFSRDSTGGADIPSPCEMKD*
>g6
MFQGAHGIALHAVQGNRASSRLEGKSHGFSRVAAGTWGTFSTYGGDGPSTLVFVQRSQDSCRVTRDTSGVSSRLGRAKGTLLEVS
RETQGPFPVATGILGFLSMFKRSQAPSNFEAVSSACLSRCQRHVRHSVERSRGPRAFSRDSTGGADIPSPCEMKD*
>g18
MFQGAHGIALHAVQGNRASSRLEGKSHGFSRVAAGTWGTFSTYGGDGPSTLVFVQRSQDSCRVTRDTSGVSSRLGRAKGTLLEVS
RETQGPFPVATGILGFLSMFKRSQAPSNFEAVSSACLSRCQRHVRHSVERSRGPRAFSRDSTGGADIPSPCEMKD*
>g15

MFQGAHGIALHAVQGNRASSRLEGKSHGFSRVAAGTWGTFSTYGGDGPSTLVFVQRSQDSCRVTRDTSGVSSRLGRAKGTLLEVS
RETQGPFPVATGILGFLSMFKRSQAPSNFEAVSSACLSRCQRHVRHSVERSRGPRAFSRDSTGGADIPSPCEMKD*
>g4
MFQGAHGIALHAVQGNRASSRLEGKSHGFSRVAAGTWGTFSTYGGDGPSTLVFVQRSQDSCRVTRDTSGVSSRLGRAKGTLLEVS
RETQGPFPVATGILGFLSMFKRSQAPSNFEAVSSACLSRCQRHVRHSVERSRGPRAFSRDSTGGADIPSPCEMKD*
>g17
MFQGAHGIALHAVQGNRASSRLEGKSHGFSRVAAGTWGTFSTYGGDGPSTLVFVQRSQDSCRVTRDTSGVSSRLGRAKGTLLEVS
RETQGPFPVATGILGFLSMFKRSQAPSNFEAVSSACLSRCQRHVRHSVERSRGPRAFSRDSTGGADIPSPCEMKD*
>g10
MFQGAHGIALHAVQGNRASSRLEGKSHGFSRVAAGTWGTFSTYGGDGPSTLVFVQRSQDSCRVTRDTSGVSSRLGRAKGTLLEVS
RETQGPFPVATGILGFLSMFKRSQAPSNFEAVSSACLSRCQRHVRHSVERSRGPRAFSRDSTGGADIPSPCEMKD*
>g2
MFQGAHGIALHAVQGNRASSRLEGKSHGFSRVAAGTWGTFSTYGGDGPSTLVFVQRSQDSCRVTRDTSGVSSRLGRAKGTLLEVS
RETQGPFPVATGILGFLSMFKRSQAPSNFEAVSSACLSRCQRHVRHSVERSRGPRAFSRDSTGGADIPSPCEMKD*
>g14
MFQGAHGIALHAVQGNRASSRLEGKSHGFSRVAAGTWGTFSTYGGDGPSTLVFVQRSQDSCRVTRDTSGVSSRLGRAKGTLLEVS
RETQGPFPVATGILGFLSMFKRSQAPSNFEAVSSACLSRCQRHVRHSVERSRGPRAFSRDSTGGADIPSPCEMKD*
>g13
MFQGAHGIALHAVQGNRASSRLEGKSHGFSRVAAGTWGTFSTYGGDGPSTLVFVQRSQDSCRVTRDTSGVSSRLGRAKGTLLEVS
RETQGPFPVATGILGFLSMFKRSQAPSNFEAVSSACLSRCQRHVRHSVERSRGPRAFSRDSTGGADIPSPCEMKD*
>g20
MFQGAHGIALHAVQGNRASSRLEGKSHGFSRVAAGTWGTFSTYGGDGPSTLVFVQRSQDSCRVTRDTSGVSSRLGRAKGTLLEVS
RETQGPFPVATGILGFLSMFKRSQAPSNFEAVSSACLSRCQRHVRHSVERSRGPRAFSRDSTGGADIPSPCEMKD*
>g16
MFQGAHGIALHAVQGNRASSRLEGKSHGFSRVAAGTWGTFSTYGGDGPSTLVFVQRSQDSCRVTRDTSGVSSRLGRAKGTLLEVS
RETQGPFPVATGILGFLSMFKRSQAPSNFEAVSSACLSRCQRHVRHSVERSRGPRAFSRDSTGGADIPSPCEMKD*
>g9
MFQGAHGIALHAVQGNRASSRLEGKSHGFSRVAAGTWGTFSTYGGDGPSTLVFVQRSQDSCRVTRDTSGVSSRLGRAKGTLLEVS
RETQGPFPVATGILGFLSMFKRSQAPSNFEAVSSACLSRCQRHVRHSVERSRGPRAFSRDSTGGADIPSPCEMKD*

### ii. SRR26335613__GENOMAD__Virus.1 (VSLC-1) intergenic sequences between repeat paralogs

>g1-g2 SRR26335613__contig_1497 SRR26335613__GENOMAD__Virus.1_1305:1803
ACTGTATTCAAGCCACTGCAAGGAAACCCGGCCTCCTTTCGAGTCAGGGCTTCTCGGTGTATATGCCCATTGAGGCAGCAATCTCAGGGTCCCTCCCTCATACCTATT
GCTGAGAGAAGCCTCCTCTTGAGGGGCTTGTGGAAAGGTGGCCTACGTCTTGAGTCTAAGCCAGGGAATCAGCTCTCATCTCGAGGCAATTTGGGGTACACGGAG
CAGTCCTCGAGTTGCTCTGCTGAACTTGGTACTCCTCTAGACTTGGCCCGTTGTTCTCGGGGAACCTCTGGAGTTGCCTAAAGGGAGTCAAGCCTCAGGTCGTGTT
TGATGGGGAACGGGGGATGGCTCTGGAGTCAATGCAGGGGAAGCGGGCCTCATCTCGAGTTGATTTGGGGCCCATGGAGCTCCTTCGCGATGCTGGTGTGACCT
CAGGGTCCCTCTCTACTTGTGACAGTGTTTTTGCGGACTGTCTCGACATCCATCAAGCAAGTCGAGGCTCCTGAC
>g2-g3 SRR26335613__contig_1497 SRR26335613__GENOMAD__Virus.1_2286:2815
ACTGTATTCAAGCCACTGCAAGGAAACCCGGCCTCCTTTCGAGTCAGGGCTTCTCGGTGTATATGCCCATTGAGGCAGCAATCTCAGGGTCCCTCCCTCATACCTATT
GCTGAGAGAAGCCTCCTCTTGAGGGGCTTGTGGAAAGGTGGCCTACGTCTTGAGTCTAAGCCAGGGAATCAGCTCTCATCTCGAGGCAATTTGGGGTACACGGAG
CAGTCCTCGAGTTGCTCTGCTGAACTTGGTACTCCTCTAGACTTGGCCCGTTGTTCTCGGGGAACCTCTGGAGTTGCCTAAAGGGAGTCAAGCCTCAGGTCGTGTT
TGATGGGGAACGGGGGATGGCTCTGGAGTCAATGCAGGGGAAGCGGGCCTCATCTCGAGTTGATTTGGGGCCCATGGAGCTCCTTCGCGATGCTGGTGTGACCT
CAGGGTCCCTCTCTACTTGTGACAGTGTTTTTGCGGACTGTCTCGACATCCATCAAGCAAGTCGAGGCTCCTGACATGTTTCAGGGGGCTCACGGAATTGCTCTGC
>g3-g4 SRR26335613__contig_1497 SRR26335613__GENOMAD__Virus.1_3268:3766
ACTGTATTCAAGCCACTGCAAGGAAACCCGGCCTCCTTTCGAGTCAGGGCTTCTCGGTGTATATGCCCATTGAGGCAGCAATCTCAGGGTCCCTCCCTCATACCTATT
GCTGAGAGAAGCCTCCTCTTGAGGGGCTTGTGGAAAGGTGGCCTACGTCTTGAGTCTAAGCCAGGGAATCAGCTCTCATCTCGAGGCAATTTGGGGTACACGGAG
CAGTCCTCGAGTTGCTCTGCTGAACTTGGTACTCCTCTAGACTTGGCCCGTTGTTCTCGGGGAACCTCTGGAGTTGCCTAAAGGGAGTCAAGCCTCAGGTCGTGTT
TGATGGGGAACGGGGGATGGCTCTGGAGTCAATGCAGGGGAAGCGGGCCTCATCTCGAGTTGATTTGGGGCCCATGGAGCTCCTTCGCGATGCTGGTGTGACCT
CAGGGTCCCTCTCTACTTGTGACAGTGTTTTTGCGGACTGTCTCGACATCCATCAAGCAAGTCGAGGCTCCTGAC
>g4-g5 SRR26335613__contig_1497 SRR26335613__GENOMAD__Virus.1_4249:4747
ACTGTATTCAAGCCACTGCAAGGAAACCCGGCCTCCTTTCGAGTCAGGGCTTCTCGGTGTATATGCCCATTGAGGCAGCAATCTCAGGGTCCCTCCCTCATACCTATT
GCTGAGAGAAGCCTCCTCTTGAGGGGCTTGTGGAAAGGTGGCCTACGTCTTGAGTCTAAGCCAGGGAATCAGCTCTCATCTCGAGGCAATTTGGGGTACACGGAG
CAGTCCTCGAGTTGCTCTGCTGAACTTGGTACTCCTCTAGACTTGGCCCGTTGTTCTCCGGGAACCTCTGGAGTTGCCTAAAGGGAGTCAAGCCTCAGGTCGTGTTT
GATGGGGAACGGGGGATGGCTCTGGAGTCAATGCAGGGGAAGCGGGCCTCATCTCGAGTTGATTTGGGGCCCATGGAGCTCCTTCGCGATGCTGGTGTGACCTC
AGGGTCCCTCTCTACTTGTGACAGTGTTTTTGCGGACTGTCTCGACATCCATCAAGCAAGTCGAGGCTCCTGAC
>g5-g6 SRR26335613__contig_1497 SRR26335613__GENOMAD__Virus.1_5230:5728
ACTGTATTCAAGCCACTGCAAGGAAACCCGGCCTCCTTTCGAGTCAGGGCTTCTCGGTGTATATGCCCATTGAGGCAGCAATCTCAGGGTCCCTCCCTCATACCTATT
GCTGAGAGAAGCCTCCTCTTGAGGGGCTTGTGGAAAGGTGGCCTACGTCTTGAGTCTAAGCCAGGGAATCAGCTCTCATCTCGAGGCAATTTGGGGTACACGGAG
CAGTCCTCGAGTTGCTCTGCTGAACTTGGTACTCCTCTAGACTTGGCCCGTTGTTCTCCGGGAACCTCTGGAGTTGCCTAAAGGGAGTCAAGCCTCAGGTCGTGTTT
GATGGGGAACGGGGGATGGCTCTGGAGTCAATGCAGGGGAAGCGGGCCTCATCTCGAGTTGATTTGGGGCCCATGGAGCTCCTTCGCGATGCTGGTGTGACCTC
AGGGTCCCTCTCTACTTGTGACAGTGTTTTTGCGGACTGTCTCGACATCCATCAAGCAAGTCGAGGCTCCTGAC
>g6-g7 SRR26335613__contig_1497 SRR26335613__GENOMAD__Virus.1_6211:6709
ACTGTATTCAAGCCACTGCAAGGAAACCCGGCCTCCTTTCGAGTCAGGGCTTCTCGGTGTATATGCCCATTGAGGCAGCAATCTCAGGGTCCCTCCCTCATACCTATT
GCTGAGAGAAGCCTCCTCTTGAGGGGCTTGTGGAAAGGTGGCCTACGTCTTGAGTCTAAGCCAGGGAATCAGCTCTCATCTCGAGGCAATTTGGGGTACACGGAG
CAGTCCTCGAGTTGCTCTGCTGAACTTGGTACTCCTCTAGACTTGGCCCGTTGTTCTCGGGGAACCTCTGGAGTTGCCTAAAGGGAGTCAAGCCTCAGGTCGTGTT
TGATGGGGAACGGGGGATGGCTCTGGAGTCAATGCAGGGGAAGCGGGCCTCATCTCGAGTTGATTTGGGGCCCATGGAGCTCCTTCGCGATGCTGGTGTGACCT
CAGGGTCCCTCTCTACTTGTGACAGTGTTTTTGCGGACTGTCTCGACATCCATCAAGCAAGTCGAGGCTCCTGAC

>g7-g8 SRR26335613__contig_1497 SRR26335613__GENOMAD__Virus.1_7192:7721
ACTGTATTCAAGCCACTGCAAGGAAACCCGGCCTCCTTTCGAGTCAGGGCTTCTCGGTGTATATGCCCATTGAGGCAGCAATCTCAGGGTCCCTCCCTCATACCTATT
GCTGAGAGAAGCCTCCTCTTGAGGGGCTTGTGGAAAGGTGGCCTACGTCTTGAGTCTAAGCCAGGGAATCAGCTCTCATCTCGAGGCAATTTGGGGTACACGGAG
CAGTCCTCGAGTTGCTCTGCTGAACTTGGTACTCCTCTAGACTTGGCCCGTTGTTCTCGGGGAACCTCTGGAGTTGCCTAAAGGGAGTCAAGCCTCAGGTCGTGTT
TGATGGGGAACGGGGGATGGCTCTGGAGTCAATGCAGGGGAAGCGGGCCTCATCTCGAGTTGATTTGGGGCCCATGGAGCTCCTTCGCGATGCTGGTGTGACCT
CAGGGTCCCTCTCTACTTGTGACAGTGTTTTTGCGGACTGTCTCGACATCCATCAAGCAAGTCGAGGCTCCTGACATGTTTCAGGGGGCTCACGGAATTGCTCTGC
>g8-g9 SRR26335613__contig_1497 SRR26335613__GENOMAD__Virus.1_8174:8672
ACTGTATTCAAGCCACTGCAAGGAAACCCGGCCTCCTTTCGAGTCAGGGCTTCTCGGTGTATATGCCCATTGAGGCAGCAATCTCAGGGTCCCTCCCTCATACCTATT
GCTGAGAGAAGCCTCCTCTTGAGGGGCTTGTGGAAAGGTGGCCTACGTCTTGAGTCTAAGCCAGGGAATCAGCTCTCATCTCGAGGCAATTTGGGGTACACGGAG
CAGTCCTCGAGTTGCTCTGCTGAACTTGGTACTCGTCTAGACTTGGCCCGTTGTTCTCCGGGAACCTCTGGAGTTGCCTAAAGGGAGTCAAGCCTCAGGTCGTGTT
TGATGGGGAACTGGGGATGGCTCTGGAGTCAATGCAGGGGAAGCGGGCCTCATCTCGAGTTGATTTGGGGCCCATGGAGCTCCTTCGCGATGCTGGTGTGACCTC
AGGGTCCCTCTCTACTTGTGACAGTGTTTTTGCGGACTGTCTCGACATCCATCAAGCAAGTCGAGGCTCCTGAC
>g9-g10 SRR26335613__contig_1497 SRR26335613__GENOMAD__Virus.1_9155:9653
ACTGTATTCAAGCCACTGCAAGGAAACCCGGCCTCCTTTCGAGTCAGGGCTTCTCGGTGTATATGCCCATTGAGGCAGCAATCTCAGGGTCCCTCCCTCATACCTATT
GCTGAGAGAAGCCTCCTCTTGAGGGGCTTGTGGAAAGGTGGCCTACGTCTTGAGTCTAAGCCAGGGAATCAGCTCTCATCTCGAGGCAATTTGGGGTACACGGAG
CAGTCCTCGAGTTGCTCTGCTGAACTTGGTACTCCTCTAGACTTGGCCCGTTGTTCTCGGGGAACCTCTGGAGTTGCCTAAAGGGAGTCAAGCCTCAGGTCGTGTT
TGATGGGGAACGGGGGATGGCTCTGGAGTCAATGCAGGGGAAGCGGGCCTCATCTCGAGTTGATTTGGGGCCCATGGAGCTCCTTCGCGATGCTGGTGTGACCT
CAGGGTCCCTCTCTACTTGTGACAGTGTTTTTGCGGACTGTCTCGACATCCATCAAGCAAGTCGAGGCTCCTGAC
>g10-g11 SRR26335613__contig_1497 SRR26335613__GENOMAD__Virus.1_10136:10634
ACTGTATTCAAGCCACTGCAAGGAAACCCGGCCTCCTTTCGAGTCAGGGCTTCTCGGTGTATATGCCCATTGAGGCAGCAATCTCAGGGTCCCTCCCTCATACCTATT
GCTGAGAGAAGCCTCCTCTTGAGGGGCTTGTGGAAAGGTGGCCTACGTCTTGAGTCTAAGCCAGGGAATCAGCTCTCATCTCGAGGCAATTTGGGGTACACGGAG
CAGTCCTCGAGTTGCTCTGCTGAACTTGGTACTCCTCTAGACTTGGCCCGTTGTTCTCCGGGAACCTCTGGAGTTGCCTAAAGGGAGTCAAGCCTCAGGTCGTGTTT
GATGGGGAACAGGGGATGGCTCTGGAGTCAATGCAGGGGAAGCGGGCCTCATCTCGAGTTGATTTGGGGCCCATGGAGCTCCTTCGCGATGCTGGTGTGACCTC
AGGGTCCCTCTCTACTTGTGACAGTGTTTTTGCGGACTGTCTCGACATCCATCAAGCAAGTCGAGGCTCCTGAC
>g11-g12 SRR26335613__contig_1497 SRR26335613__GENOMAD__Virus.1_11117:11615
ACTGTATTCAAGCCACTGCAAGGAAACCCGGCCTCCTTTCGAGTCAGGGCTTCTCGGTGTATATGCCCATTGAGGCAGCAATCTCAGGGTCCCTCCCTCATACCTATT
GCTGAGAGAAGCCTCCTCTTGAGGGGCTTGTGGAAAGGTGGCCTACGTCTTGAGTCTAAGCCAGGGAATCAGCTCTCATCTCGAGGCAATTTGGGGTACACGGAG
CAGTCCTCGAGTTGCTCTGCTGAACTTGGTACTCCTCTAGACTTGGCCCGTTGTTCTCGGGGAACCTCTGGAGTTGCCTAAAGGGAGTCAAGCCTCAGGTCGTGTT
TGATGGGGAACGGGGGATGGCTCTGGAGTCAATGCAGGGGAAGCGGGCCTCATCTCGAGTTGATTTGGGGCCCATGGAGCTCCTTCGCGATGCTGGTGTGACCT
CAGGGTCCCTCTCTACTTGTGACAGTGTTTTTGCGGACTGTCTCGACATCCATCAAGCAAGTCGAGGCTCCTGAC
>g12-g13 SRR26335613__contig_1497 SRR26335613__GENOMAD__Virus.1_12098:12596
ACTGTATTCAAGCCACTGCAAGGAAACCCGGCCTCCTTTCGAGTCAGGGCTTCTCGGTGTATATGCCCATTGAGGCAGCAATCTCAGGGTCCCTCCCTCATACCTATT
GCTGAGAGAAGCCTCCTCTTGAGGGGCTTGTGGAAAGGTGGCCTACGTCTTGAGTCTAAGCCAGGGAATCAGCTCTCATCTCGAGGCAATTTGGGGTACACGGAG
CAGTCCTCGAGTTGCTCTGCTGAACTTGGTACTCCTCTAGACTTGGCCCGTTGTTCTCGGGGAACCTCTGGAGTTGCCTAAAGGGAGTCAAGCCTCAGGTCGTGTT
TGATGGGGAACGGGGGATGGCTCTGGAGTCAATGCAGGGGAAGCGGGCCTCATCTCGAGTTGATTTGGGGCCCATGGAGCTCCTTCGCGATGCTGGTGTGACCT
CAGGGTCCCTCTCTACTTGTGACAGTGTTTTTGCGGACTGTCTCGACATCCATCAAGCAAGTCGAGGCTCCTGAC
>g13-g14 SRR26335613__contig_1497 SRR26335613__GENOMAD__Virus.1_13079:13577
ACTGTATTCAAGCCACTGCAAGGAAACCCGGCCTCCTTTCGAGTCAGGGCTTCTCGGTGTATATGCCCATTGAGGCAGCAATCTCAGGGTCCCTCCCTCATACCTATT
GCTGAGAGAAGCCTCCTCTTGAGGGGCTTGTGGAAAGGTGGCCTACGTCTTGAGTCTAAGCCAGGGAATCAGCTCTCATCTCGAGGCAATTTGGGGTACACGGAG
CAGTCCTCGAGTTGCTCTGCTGAACTTGGTACTCCTCTAGACTTGGCCCGTTGTTCTCGGGGAACCTCTGGAGTTGCCTAAAGGGAGTCAAGCCTCAGGTCGTGTT
TGATGGGGAACGGGGGATGGCTCTGGAGTCAATGCAGGGGAAGCGGGCCTCATCTCGAGTTGATTTGGGGCCCATGGAGCTCCTTCGCGATGCTGGTGTGACCT
CAGGGTCCCTCTCTACTTGTGACAGTGTTTTTGCGGACTGTCTCGACATCCATCAAGCAAGTCGAGGCTCCTGAC
>g14-g15 SRR26335613__contig_1497 SRR26335613__GENOMAD__Virus.1_14060:14558
ACTGTATTCAAGCCACTGCAAGGAAACCCGGCCTCCTTTCGAGTCAGGGCTTCTCGGTGTATATGCCCATTGAGGCAGCAATCTCAGGGTCCCTCCCTCATACCTATT
GCTGAGAGAAGCCTCCTCTTGAGGGGCTTGTGGAAAGGTGGCCTACGTCTTGAGTCTAAGCCAGGGAATCAGCTCTCATCTCGAGGCAATTTGGGGTACACGGAG
CAGTCCTCGAGTTGCTCTGCTGAACTTGGTACTCCTCTAGACTTGGCCCGTTGTTCTCGGGGAACCTCTGGAGTTGCCTAAAGGGAGTCAAGCCTCAGGTCGTGTT
TGATGGGGAACGGGGGATGGCTCTGGAGTCAATGCAGGGGAAGCGGGCCTCATCTCGAGTTGATTTGGGGCCCATGGAGCTCCTTCGCGATGCTGGTGTGACCT
CAGGGTCCCTCTCTACTTGTGACAGTGTTTTTGCGGACTGTCTCGACATCCATCAAGCAAGTCGAGGCTCCTGAC
>g15-g16 SRR26335613__contig_1497 SRR26335613__GENOMAD__Virus.1_15041:15539
ACTGTATTCAAGCCACTGCAAGGAAACCCGGCCTCCTTTCGAGTCAGGGCTTCTCGGTGTATATGCCCATTGAGGCAGCAATCTCAGGGTCCCTCCCTCATACCTATT
GCTGAGAGAAGCCTCCTCTTGAGGGGCTTGTGGAAAGGTGGCCTACGTCTTGAGTCTAAGCCAGGGAATCAGCTCTCATCTCGAGGCAATTTGGGGTACACGGAG
CAGTCCTCGAGTTGCTCTGCTGAACTTGGTACTCCTCTAGACTTGGCCCGTTGTTCTCGGGGAACCTCTGGAGTTGCCTAAAGGGAGTCAAGCCTCAGGTCGTGTT
TGATGGGGAACGGGGGATGGCTCTGGAGTCAATGCAGGGGAAGCGGGCCTCATCTCGAGTTGATTTGGGGCCCATGGAGCTCCTTCGCGATGCTGGTGTGACCT
CAGGGTCCCTCTCTACTTGTGACAGTGTTTTTGCGGACTGTCTCGACATCCATCAAGCAAGTCGAGGCTCCTGAC
>g16-g17 SRR26335613__contig_1497 SRR26335613__GENOMAD__Virus.1_16022:16520
ACTGTATTCAAGCCACTGCAAGGAAACCCGGCCTCCTTTCGAGTCAGGGCTTCTCGGTGTATATGCCCATTGAGGCAGCAATCTCAGGGTCCCTCCCTCATACCTATT
GCTGAGAGAAGCCTCCTCTTGAGGGGCTTGTGGAAAGGTGGCCTACGTCTTGAGTCTAAGCCAGGGAATCAGCTCTCATCTCGAGGCAATTTGGGGTACACGGAG
CAGTCCTCGAGTTGCTCTGCTGAACTTGGTACTCCTCTAGACTTGGCCCGTTGTTCTCGGGGAACCTCTGGAGTTGCCTAAAGGGAGTCAAGCCTCAGGTCGTGTT
TGATGGGGAACGGGGGATGGCTCTGGAGTCAATGCAGGGGAAGCGGGCCTCATCTCGAGTTGATTTGGGGCCCATGGAGCTCCTTCGCGATGCTGGTGTGACCT
CAGGGTCCCTCTCTACTTGTGACAGTGTTTTTGCGGACTGTCTCGACATCCATCAAGCAAGTCGAGGCTCCTGAC
>g17-g18 SRR26335613__contig_1497 SRR26335613__GENOMAD__Virus.1_17003:17500
ACTGTATTCAAGCCACTGCAAGGAAACCCGGCCTCCTTTCGAGTCAGGGCTTCTCGGTGTATATGCCCATTGAGGCAGCAATCTCAGGGTCCCTCCCTCATACCTATT
GCTGAGAGAAGCCTCCTCTTGAGGGGCTTGTGGAAAGGTGGCCTACGTCTTGAGTCTAAGCCAGGGAATCAGCTCTCATCTCGAGGCAATTTGGGGTACACGGAG
CAGTCCTCGAGTTGCTCTGCTGAACTTGGTACTCCTCTAGACTTGGCCCGTTGTTCTCGGGAACCTCTGGAGTTGCCTAAAGGGAGTCAAGCCTCAGGTCGTGTTT
GATGGGGAACGGGGGATGGCTCTGGAGTCAATGCAGGGGAAGCGGGCCTCATCTCGAGTTGATTTGGGGCCCATGGAGCTCCTTCGCGATGCTGGTGTGACCTC
AGGGTCCCTCTCTACTTGTGACAGTGTTTTTGCGGACTGTCTCGACATCCATCAAGCAAGTCGAGGCTCCTGAC

>g18-g19 SRR26335613__contig_1497 SRR26335613__GENOMAD__Virus.1_17983:18481
ACTGTATTCAAGCCACTGCAAGGAAACCCGGCCTCCTTTCGAGTCAGGGCTTCTCGGTGTATATGCCCATTGAGGCAGCAATCTCAGGGTCCCTCCCTCATACCTATT
GCTGAGAGAAGCCTCCTCTTGAGGGGCTTGTGGAAAGGTGGCCTACGTCTTGAGTCTAAGCCAGGGAATCAGCTCTCATCTCGAGGCAATTTGGGGTACACGGAG
CAGTCCTCGAGTTGCTCTGCTGAACTTGGTACTCCTCTAGACTTGGCCCGTTGTTCTCGGGGAACCTCTGGAGTTGCCTAAAGGGAGTCAAGCCTCAGGTCGTGTT
TGATGGGGAACGGGGGATGGCTCTGGAGTCAATGCAGGGGAAGCGGGCCTCATCTCGAGTTGATTTGGGGCCCATGGAGCTCCTTCGCGATGCTGGTGTGACCT
CAGGGTCCCTCTCTACTTGTGACAGTGTTTTTGCGGACTGTCTCGACATCCATCAAGCAAGTCGAGGCTCCTGAC
>g19-g20 SRR26335613__contig_1497 SRR26335613__GENOMAD__Virus.1_18964:19462
ACTGTATTCAAGCCACTGCAAGGAAACCCGGCCTCCTTTCGAGTCAGGGCTTCTCGGTGTATATGCCCATTGAGGCAGCAATCTCAGGGTCCCTCCCTCATACCTATT
GCTGAGAGAAGCCTCCTCTTGAGGGGCTTGTGGAAAGGTGGCCTACGTCTTGAGTCTAAGCCAGGGAATCAGCTCTCATCTCGAGGCAATTTGGGGTACACGGAG
CAGTCCTCGAGTTGCTCTGCTGAACTTGGTACTCCTCTAGACTTGGCCCGTTGTTCTCGGGGAACCTCTGGAGTTGCCTAAAGGGAGTCAAGCCTCAGGTCGTGTT
TGATGGGGAACGGGGGATGGCTCTGGAGTCAATGCAGGGGAAGCGGGCCTCATCTCGAGTTGATTTGGGGCCCATGGAGCTCCTTCGCGATGCTGGTGTGACCT
CAGGGTCCCTCTCTACTTGTGACAGTGTTTTTGCGGACTGTCTCGACATCCATCAAGCAAGTCGAGGCTCCTGAC
>g20-g21 SRR26335613__contig_1497 SRR26335613__GENOMAD__Virus.1_19945:20443
ACTGTATTCAAGCCACTGCAAGGAAACCCGGCCTCCTTTCGAGTCAGGGCTTCTCGGTGTATATGCCCATTGAGGCAGCAATCTCAGGGTCCCTCCCTCATACCTATT
GCTGAGAGAAGCCTCCTCTTGAGGGGCTTGTGGAAAGGTGGCCTACGTCTTGAGTCTAAGCCAGGGAATCAGCTCTCATCTCGAGGCAATTTGGGGTACACGGAG
CAGTCCTCGAGTTGCTCTGCTGAACTTGGTACTCCTCTAGACTTGGCCCGTTGTTCTCGGGGAACCTCTGGAGTTGCCTAAAGGGAGTCAAGCCTCAGGTCGTGTT
TGATGGGGAACGGGGGATGGCTCTGGAGTCAATGCAGGGGAAGCGGGCCTCATCTCGAGTTGATTTGGGGCCCATGGAGCTCCTTCGCGATGCTGGTGTGACCT
CAGGGTCCCTCTCTACTTGTGACAGTGTTTTTGCGGACTGTCTCGACATCCATCAAGCAAGTCGAGGCTCCTGAC
>g21-g22 SRR26335613__contig_1497 SRR26335613__GENOMAD__Virus.1_20926:21455
ACTGTATTCAAGCCACTGCAAGGAAACCCGGCCTCCTTTCGAGTCAGGGCTTCTCGGTGTATATGCCCATTGAGGCAGCAATCTCAGGGTCCCTCCCTCATACCTATT
GCTGAGAGAAGCCTCCTCTTGAGGGGCTTGTGGAAAGGTGGCCTACGTCTTGAGTCTAAGCCAGGGAATCAGCTCTCATCTCGAGGCAATTTGGGGTACACGGAG
CAGTCCTCGAGTTGCTCTGCTGAACTTGGTACTCCTCTAGACTTGGCCCGTTGTTCTCGGGGAACCTCTGGAGTTGCCTAAAGGGAGTCAAGCCTCAGGTCGTGTT
TGATGGGGAACGGGGGATGGCTCTGGAGTCAATGCAGGGGAAGCGGGCCTCATCTCGAGTTGATTTGGGGCCCATGGAGCTCCTTCGCGATGCTGGTGTGACCT
CAGGGTCCCTCTCTACTTGTGACAGTGTTTTTGCGGACTGTCTCGACATCCATCAAGCAAGTCGAGGCTCCTGACATGTTTCAGGGGGCTCACGGAATTGCTCTGC
>g22-g23 SRR26335613__contig_1497 SRR26335613__GENOMAD__Virus.1_21908:22437
ACTGTATTCAAGCCACTGCAAGGAAACCCGGCCTCCTTTCGAGTCAGGGCTTCTCGGTGTATATGCCCATTGAGGCAGCAATCTCAGGGTCCCTCCCTCATACCTATT
GCTGAGAGAAGCCTCCTCTTGAGGGGCTTGTGGAAAGGTGGCCTACGTCTTGAGTCTAAGCCAGGGAATCAGCTCTCATCTCGAGGCAATTTGGGGTACACGGAG
CAGTCCTCGAGTTGCTCTGCTGAACTTGGTACTCCTCTAGACTTGGCCCGTTGTTCTCGGGGAACCTCTGGAGTTGCCTAAAGGGAGTCAAGCCTCAGGTCGTGTT
TGATGGGGAACGGGGGATGGCTCTGGAGTCAATGCAGGGGAAGCGGGCCTCATCTCGAGTTGATTTGGGGCCCATGGAGCTCCTTCGCGATGCTGGTGTGACCT
CAGGGTCCCTCTCTACTTGTGACAGTGTTTTTGCGGACTGTCTCGACATCCATCAAGCAAGTCGAGGCTCCTGACATGTTTCAGGGGGCTCACGGAATTGCTCTGC
>g23-g24 SRR26335613__contig_1497 SRR26335613__GENOMAD__Virus.1_22890:23388
ACTGTATTCAAGCCACTGCAAGGAAACCCGGCCTCCTTTCGAGTCAGGGCTTCTCGGTGTATATGCCCATTGAGGCAGCAATCTCAGGGTCCCTCCCTCATACCTATT
GCTGAGAGAAGCCTCCTCTTGAGGGGCTTGTGGAAAGGTGGCCTACGTCTTGAGTCTAAGCCAGGGAATCAGCTCTCATCTCGAGGCAATTTGGGGTACACGGAG
CAGTCCTCGAGTTGCTCTGCTGAACTTGGTACTCCTCTAGACTTGGCCCGTTGTTCTCGGGGAACCTCTGGAGTTGCCTAAAGGGAGTCAAGCCTCAGGTCGTGTT
TGATGGGGAACGGGGGATGGCTCTGGAGTCAATGCAGGGGAAGCGGGCCTCATCTCGAGTTGATTTGGGGCCCATGGAGCTCCTTCGCGATGCTGGTGTGACCT
CAGGGTCCCTCTCTACTTGTGACAGTGTTTTTGCGGACTGTCTCGACATCCATCAAGCAAGTCGAGGCTCCTGAC
>g24-g25 SRR26335613__contig_1497 SRR26335613__GENOMAD__Virus.1_23871:24400
ACTGTATTCAAGCCACTGCAAGGAAACCCGGCCTCCTTTCGAGTCAGGGCTTCTCGGTGTATATGCCCATTGAGGCAGCAATCTCAGGGTCCCTCCCTCATACCTATT
GCTGAGAGAAGCCTCCTCTTGAGGGGCTTGTGGAAAGGTGGCCTACGTCTTGAGTCTAAGCCAGGGAATCAGCTCTCATCTCGAGGCAATTTGGGGTACACGGAG
CAGTCCTCGAGTTGCTCTGCTGAACTTGGTACTCCTCTAGACTTGGCCCGTTGTTCTCGGGGAACCTCTGGAGTTGCCTAAAGGGAGTCAAGCCTCAGGTCGTGTT
TGATGGGGAACGGGGGATGGCTCTGGAGTCAATGCAGGGGAAGCGGGCCTCATCTCGAGTTGATTTGGGGCCCATGGAGCTCCTTCGCGATGCTGGTGTGACCT
CAGGGTCCCTCTCTACTTGTGACAGTGTTTTTGCGGACTGTCTCGACATCCATCAAGCAAGTCGAGGCTCCTGACATGTTTCAGGGGGCTCACGGAATTGCTCTGC
>g25-g26 SRR26335613__contig_1497 SRR26335613__GENOMAD__Virus.1_24853:25382
ACTGTATTCAAGCCACTGCAAGGAAACCCGGCCTCCTTTCGAGTCAGGGCTTCTCGGTGTATATGCCCATTGAGGCAGCAATCTCAGGGTCCCTCCCTCATACCTATT
GCTGAGAGAAGCCTCCTCTTGAGGGGCTTGTGGAAAGGTGGCCTACGTCTTGAGTCTAAGCCAGGGAATCAGCTCTCATCTCGAGGCAATTTGGGGTACACGGAG
CAGTCCTCGAGTTGCTCTGCTGAACTTGGTACTCCTCTAGACTTGGCCCGTTGTTCTCCGGGAACCTCTGGAGTTGCCTAAAGGGAGTCAAGCCTCAGGTCGTGTTT
GATGGGGAACGGGGGATGGCTCTGGAGTCAATGCAGGGGAAGCGGGCCTCATCTCGAGTTGATTTGGGGCCCATGGAGCTCCTTCGCGATGCTGGTGTGACCTC
AGGGTCCCTCTCTACTTGTGACAGTGTTTTTGCGGACTGTCTCGACATCCATCAAGCAAGTCGAGGCTCCTGACATGTTTCAGGGGGCTCACGGAATTGCTCTGC
>g26-g27 SRR26335613__contig_1497 SRR26335613__GENOMAD__Virus.1_25835:26333
ACTGTATTCAAGCCACTGCAAGGAAACCCGGCCTCCTTTCGAGTCAGGGCTTCTCGGTGTATATGCCCATTGAGGCAGCAATCTCAGGGTCCCTCCCTCATACCTATT
GCTGAGAGAAGCCTCCTCTTGAGGGGCTTGTGGAAAGGTGGCCTACGTCTTGAGTCTAAGCCAGGGAATCAGCTCTCATCTCGAGGCAATTTGGGGTACACGGAG
CAGTCCTCGAGTTGCTCTGCTGAACTTGGTACTCCTCTAGACTTGGCCCGTTGTTCTCGGGGAACCTCTGGAGTTGCCTAAAGGGAGTCAAGCCTCAGGTCGTGTT
TGATGGGGAACGGGGGATGGCTCTGGAGTCAATGCAGGGGAAGCGGGCCTCATCTCGAGTTGATTTGGGGCCCATGGAGCTCCTTCGCGATGCTGGTGTGACCT
CAGGGTCCCTCTCTACTTGTGACAGTGTTTTTGCGGACTGTCTCGACATCCATCAAGCAAGTCGAGGCTCCTGAC
>g27-g28 SRR26335613__contig_1497 SRR26335613__GENOMAD__Virus.1_26816:27393
ACTGTATTCAAGCCACTGCAAGGAAACCCGGCCTCCTTTCGAGTCAGGGCTTCTCGGTGTATATGCCCATTGAGGCAGCAATCTCAGGGTCCCTCCCTCATACCTATT
GCTGAGAGAAGCCTCCTCTTGAGGGGCTTGTGGAAAGGTGGCCTACGTCTTGAGTCTAAGCCAGGGAATCAGCTCTCATCTCGAGGCAATTTGGGGTACACGGAG
CAGTCCTCGAGTTGCTCTGCTGAACTTGGTACTCCTCTAGACTTGGCCCGTTGTTCTCGGGGAACCTCTGGAGTTGCCTAAAGGGAGTCAAGCCTCAGGTCGTGTT
TGATGGGGAACGGGGGATGGCTCTGGAGTCAATGCAGGGGAAGCGGGCCTCATCTCGAGTTGATTTGGGGCCCATGGAGCTCCTTCGCGATGCTGGTGTGACCT
CAGGGTCCCTCTCTACTTGTGACAGTGTTTTTGCGGACTGTCTCGACATCCATCAAGCAAGTCGAGGCTCCTGACATGTTTCAGGGGGCTCACGGAATTGCTCTGCA
TGCAGTGCAGGGGAATCGGGCCTCATCTCGCTTGGAGGGGAAGTCTC

#### iii. SRR26335613__GENOMAD__Virus.1 (VSLC-1) intergenic repeated components

>$IGRC_1$
ACTGTATTCAAGCCACTGCAAGGAAAC
>$IGRC_2$

CCGGCCTCCTTTCGAGTCAGGGCTTCTCGGTGTATATGCCCATTGAGGCAGCAATCTCAGGGTCCCTCCCTCATACCTATTGCT
GAGAGAAGCCTCCTCTTGAGGGGCTTGTGGAAAGGTGGCCTACGTCTTGAGTCTAAGCCAGGGAATCAGCTCTCATCTCGAGG
CAATTTGGGGTACACGGAGCAGTCCTCGAGTTGCTCTGCTGAACTTGGTACTCCTCTAGACTTGGCCCGTTGTTCTCGGGGAA
CCTCTGGAGTTGCCTAAAGGGAGTCAAGCCTCAGGTCGTGTTTGATGGGGAACGGGGGATGGCTCTGGAGTCAATGCAGGGG
AAGCGGGCCTCATCTCGAGTTGATTTGGGGCCCATGGAGCTCCTTCGCGATGCTGGTGTGACCTCAGGGTCCCTCTCTACTTG
TGACAGTGTTTTT-GCGGACTGTCTCGACATCCATCAAGCAAGTCGAGGCTCCTGAC
>IGRC$_3$
ATGTTTCAGGGGGCTCACGGAATTGCTCTG
>IGRC$_4$
CATGCAGTGCAGGGGAATCGGCCTC--ATCTCGCTTGG

### iv. SRR26335613__GENOMAD__Virus.1 (VSLC-1) intergenic repeated components (MSA)

>g12-g13
ACTGTATTCAAGCCACTGCAAGGAAACCCGGCCTCCTTTCGAGTCAGGGCTTCTCGGTGTATATGCCCATTGAGGCAGCAATCT
CAGGGTCCCTCCCTCATACCTATTGCTGAGAGAAGCCTCCTCTTGAGGGGCTTGTGGAAAGGTGGCCTACGTCTTGAGTCTAA
GCCAGGGAATCAGCTCTCATCTCGAGGCAATTTGGGGTACACGGAGCAGTCCTCGAGTTGCTCTGCTGAACTTGGTACTCCTC
TAGACTTGGCCCGTTGTTCTCGGGGAACCTCTGGAGTTGCCTAAAGGGAGTCAAGCCTCAGGTCGTGTTTGATGGGGAACGGG
GGATGGCTCTGGAGTCAATGCAGGGGAAGCGGGCCTCATCTCGAGTTGATTTGGGGCCCATGGAGCTCCTTCGCGATGCTGG
TGTGACCTCAGGGTCCCTCTCTACTTGTGACAGTGTTTTT-GCGGACTGTCTCGACATCCATCAAGCAAGTCGAGGCTCCTGAC-
-----------------------------------------------------------------------------------------------------------------------------------------------------------------------------------------------------------------
-----------------------------------------------------------------------------------------------------------------------------------------------------------------------------------------------------------------
-----------------------------------------------------------------------------------------------------------------------------------------------------------------------------------------------------
>g15-g16
ACTGTATTCAAGCCACTGCAAGGAAACCCGGCCTCCTTTCGAGTCAGGGCTTCTCGGTGTATATGCCCATTGAGGCAGCAATCT
CAGGGTCCCTCCCTCATACCTATTGCTGAGAGAAGCCTCCTCTTGAGGGGCTTGTGGAAAGGTGGCCTACGTCTTGAGTCTAA
GCCAGGGAATCAGCTCTCATCTCGAGGCAATTTGGGGTACACGGAGCAGTCCTCGAGTTGCTCTGCTGAACTTGGTACTCCTC
TAGACTTGGCCCGTTGTTCTCGGGGAACCTCTGGAGTTGCCTAAAGGGAGTCAAGCCTCAGGTCGTGTTTGATGGGGAACGGG
GGATGGCTCTGGAGTCAATGCAGGGGAAGCGGGCCTCATCTCGAGTTGATTTGGGGCCCATGGAGCTCCTTCGCGATGCTGG
TGTGACCTCAGGGTCCCTCTCTACTTGTGACAGTGTTTTT-GCGGACTGTCTCGACATCCATCAAGCAAGTCGAGGCTCCTGAC-
-----------------------------------------------------------------------------------------------------------------------------------------------------------------------------------------------------------------
-----------------------------------------------------------------------------------------------------------------------------------------------------------------------------------------------------------------
-----------------------------------------------------------------------------------------------------------------------------------------------------------------------------------------------------
>g26-g27
ACTGTATTCAAGCCACTGCAAGGAAACCCGGCCTCCTTTCGAGTCAGGGCTTCTCGGTGTATATGCCCATTGAGGCAGCAATCT
CAGGGTCCCTCCCTCATACCTATTGCTGAGAGAAGCCTCCTCTTGAGGGGCTTGTGGAAAGGTGGCCTACGTCTTGAGTCTAA
GCCAGGGAATCAGCTCTCATCTCGAGGCAATTTGGGGTACACGGAGCAGTCCTCGAGTTGCTCTGCTGAACTTGGTACTCCTC
TAGACTTGGCCCGTTGTTCTCGGGGAACCTCTGGAGTTGCCTAAAGGGAGTCAAGCCTCAGGTCGTGTTTGATGGGGAACGGG
GGATGGCTCTGGAGTCAATGCAGGGGAAGCGGGCCTCATCTCGAGTTGATTTGGGGCCCATGGAGCTCCTTCGCGATGCTGG
TGTGACCTCAGGGTCCCTCTCTACTTGTGACAGTGTTTTT-GCGGACTGTCTCGACATCCATCAAGCAAGTCGAGGCTCCTGAC-
-----------------------------------------------------------------------------------------------------------------------------------------------------------------------------------------------------------------
-----------------------------------------------------------------------------------------------------------------------------------------------------------------------------------------------------------------
-----------------------------------------------------------------------------------------------------------------------------------------------------------------------------------------------------
>g16-g17
ACTGTATTCAAGCCACTGCAAGGAAACCCGGCCTCCTTTCGAGTCAGGGCTTCTCGGTGTATATGCCCATTGAGGCAGCAATCT
CAGGGTCCCTCCCTCATACCTATTGCTGAGAGAAGCCTCCTCTTGAGGGGCTTGTGGAAAGGTGGCCTACGTCTTGAGTCTAA
GCCAGGGAATCAGCTCTCATCTCGAGGCAATTTGGGGTACACGGAGCAGTCCTCGAGTTGCTCTGCTGAACTTGGTACTCCTC
TAGACTTGGCCCGTTGTTCTCGGGGAACCTCTGGAGTTGCCTAAAGGGAGTCAAGCCTCAGGTCGTGTTTGATGGGGAACGGG
GGATGGCTCTGGAGTCAATGCAGGGGAAGCGGGCCTCATCTCGAGTTGATTTGGGGCCCATGGAGCTCCTTCGCGATGCTGG
TGTGACCTCAGGGTCCCTCTCTACTTGTGACAGTGTTTTT-GCGGACTGTCTCGACATCCATCAAGCAAGTCGAGGCTCCTGAC-
-----------------------------------------------------------------------------------------------------------------------------------------------------------------------------------------------------------------
-----------------------------------------------------------------------------------------------------------------------------------------------------------------------------------------------------------------
-----------------------------------------------------------------------------------------------------------------------------------------------------------------------------------------------------
>g23-g24
ACTGTATTCAAGCCACTGCAAGGAAACCCGGCCTCCTTTCGAGTCAGGGCTTCTCGGTGTATATGCCCATTGAGGCAGCAATCT
CAGGGTCCCTCCCTCATACCTATTGCTGAGAGAAGCCTCCTCTTGAGGGGCTTGTGGAAAGGTGGCCTACGTCTTGAGTCTAA
GCCAGGGAATCAGCTCTCATCTCGAGGCAATTTGGGGTACACGGAGCAGTCCTCGAGTTGCTCTGCTGAACTTGGTACTCCTC
TAGACTTGGCCCGTTGTTCTCGGGGAACCTCTGGAGTTGCCTAAAGGGAGTCAAGCCTCAGGTCGTGTTTGATGGGGAACGGG
GGATGGCTCTGGAGTCAATGCAGGGGAAGCGGGCCTCATCTCGAGTTGATTTGGGGCCCATGGAGCTCCTTCGCGATGCTGG
TGTGACCTCAGGGTCCCTCTCTACTTGTGACAGTGTTTTT-GCGGACTGTCTCGACATCCATCAAGCAAGTCGAGGCTCCTGAC-
-----------------------------------------------------------------------------------------------------------------------------------------------------------------------------------------------------------------
-----------------------------------------------------------------------------------------------------------------------------------------------------------------------------------------------------------------
-----------------------------------------------------------------------------------------------------------------------------------------------------------------------------------------------------
>g6-g7
ACTGTATTCAAGCCACTGCAAGGAAACCCGGCCTCCTTTCGAGTCAGGGCTTCTCGGTGTATATGCCCATTGAGGCAGCAATCT
CAGGGTCCCTCCCTCATACCTATTGCTGAGAGAAGCCTCCTCTTGAGGGGCTTGTGGAAAGGTGGCCTACGTCTTGAGTCTAA
GCCAGGGAATCAGCTCTCATCTCGAGGCAATTTGGGGTACACGGAGCAGTCCTCGAGTTGCTCTGCTGAACTTGGTACTCCTC
TAGACTTGGCCCGTTGTTCTCGGGGAACCTCTGGAGTTGCCTAAAGGGAGTCAAGCCTCAGGTCGTGTTTGATGGGGAACGGG
GGATGGCTCTGGAGTCAATGCAGGGGAAGCGGGCCTCATCTCGAGTTGATTTGGGGCCCATGGAGCTCCTTCGCGATGCTGG
TGTGACCTCAGGGTCCCTCTCTACTTGTGACAGTGTTTTT-GCGGACTGTCTCGACATCCATCAAGCAAGTCGAGGCTCCTGAC-

------------------------------------------------------------------------------------------------------------------------------------------------------------------------------------
------------------------------------------------------------------------------------------------------------------------------------------------------------------------------------
----------------------------------------------------------------------------------------------------------------------------------------------------------------
>g1-g2
ACTGTATTCAAGCCACTGCAAGGAAACCCGGCCTCCTTTCGAGTCAGGGCTTCTCGGTGTATATGCCCATTGAGGCAGCAATCT
CAGGGTCCCTCCCTCATACCTATTGCTGAGAGAAGCCTCCTCTTGAGGGGCTTGTGGAAAGGTGGCCTACGTCTTGAGTCTAA
GCCAGGGAATCAGCTCTCATCTCGAGGCAATTTGGGGTACACGGAGCAGTCCTCGAGTTGCTCTGCTGAACTTGGTACTCCTC
TAGACTTGGCCCGTTGTTCTCGGGGAACCTCTGGAGTTGCCTAAAGGGAGTCAAGCCTCAGGTCGTGTTTGATGGGGAACGGG
GGATGGCTCTGGAGTCAATGCAGGGGAAGCGGGCCTCATCTCGAGTTGATTTGGGGCCCATGGAGCTCCTTCGCGATGCTGG
TGTGACCTCAGGGTCCCTCTCTACTTGTGACAGTGTTTTT-GCGGACTGTCTCGACATCCATCAAGCAAGTCGAGGCTCCTGAC-
------------------------------------------------------------------------------------------------------------------------------------------------------------------------------------
------------------------------------------------------------------------------------------------------------------------------------------------------------------------------------
----------------------------------------------------------------------------------------------------------------------------------------------------------------
>g14-g15
ACTGTATTCAAGCCACTGCAAGGAAACCCGGCCTCCTTTCGAGTCAGGGCTTCTCGGTGTATATGCCCATTGAGGCAGCAATCT
CAGGGTCCCTCCCTCATACCTATTGCTGAGAGAAGCCTCCTCTTGAGGGGCTTGTGGAAAGGTGGCCTACGTCTTGAGTCTAA
GCCAGGGAATCAGCTCTCATCTCGAGGCAATTTGGGGTACACGGAGCAGTCCTCGAGTTGCTCTGCTGAACTTGGTACTCCTC
TAGACTTGGCCCGTTGTTCTCGGGGAACCTCTGGAGTTGCCTAAAGGGAGTCAAGCCTCAGGTCGTGTTTGATGGGGAACGGG
GGATGGCTCTGGAGTCAATGCAGGGGAAGCGGGCCTCATCTCGAGTTGATTTGGGGCCCATGGAGCTCCTTCGCGATGCTGG
TGTGACCTCAGGGTCCCTCTCTACTTGTGACAGTGTTTTT-GCGGACTGTCTCGACATCCATCAAGCAAGTCGAGGCTCCTGAC-
------------------------------------------------------------------------------------------------------------------------------------------------------------------------------------
------------------------------------------------------------------------------------------------------------------------------------------------------------------------------------
----------------------------------------------------------------------------------------------------------------------------------------------------------------
>g9-g10
ACTGTATTCAAGCCACTGCAAGGAAACCCGGCCTCCTTTCGAGTCAGGGCTTCTCGGTGTATATGCCCATTGAGGCAGCAATCT
CAGGGTCCCTCCCTCATACCTATTGCTGAGAGAAGCCTCCTCTTGAGGGGCTTGTGGAAAGGTGGCCTACGTCTTGAGTCTAA
GCCAGGGAATCAGCTCTCATCTCGAGGCAATTTGGGGTACACGGAGCAGTCCTCGAGTTGCTCTGCTGAACTTGGTACTCCTC
TAGACTTGGCCCGTTGTTCTCGGGGAACCTCTGGAGTTGCCTAAAGGGAGTCAAGCCTCAGGTCGTGTTTGATGGGGAACGGG
GGATGGCTCTGGAGTCAATGCAGGGGAAGCGGGCCTCATCTCGAGTTGATTTGGGGCCCATGGAGCTCCTTCGCGATGCTGG
TGTGACCTCAGGGTCCCTCTCTACTTGTGACAGTGTTTTT-GCGGACTGTCTCGACATCCATCAAGCAAGTCGAGGCTCCTGAC-
------------------------------------------------------------------------------------------------------------------------------------------------------------------------------------
------------------------------------------------------------------------------------------------------------------------------------------------------------------------------------
----------------------------------------------------------------------------------------------------------------------------------------------------------------
>g11-g12
ACTGTATTCAAGCCACTGCAAGGAAACCCGGCCTCCTTTCGAGTCAGGGCTTCTCGGTGTATATGCCCATTGAGGCAGCAATCT
CAGGGTCCCTCCCTCATACCTATTGCTGAGAGAAGCCTCCTCTTGAGGGGCTTGTGGAAAGGTGGCCTACGTCTTGAGTCTAA
GCCAGGGAATCAGCTCTCATCTCGAGGCAATTTGGGGTACACGGAGCAGTCCTCGAGTTGCTCTGCTGAACTTGGTACTCCTC
TAGACTTGGCCCGTTGTTCTCGGGGAACCTCTGGAGTTGCCTAAAGGGAGTCAAGCCTCAGGTCGTGTTTGATGGGGAACGGG
GGATGGCTCTGGAGTCAATGCAGGGGAAGCGGGCCTCATCTCGAGTTGATTTGGGGCCCATGGAGCTCCTTCGCGATGCTGG
TGTGACCTCAGGGTCCCTCTCTACTTGTGACAGTGTTTTT-GCGGACTGTCTCGACATCCATCAAGCAAGTCGAGGCTCCTGAC-
------------------------------------------------------------------------------------------------------------------------------------------------------------------------------------
------------------------------------------------------------------------------------------------------------------------------------------------------------------------------------
----------------------------------------------------------------------------------------------------------------------------------------------------------------
>g18-g19
ACTGTATTCAAGCCACTGCAAGGAAACCCGGCCTCCTTTCGAGTCAGGGCTTCTCGGTGTATATGCCCATTGAGGCAGCAATCT
CAGGGTCCCTCCCTCATACCTATTGCTGAGAGAAGCCTCCTCTTGAGGGGCTTGTGGAAAGGTGGCCTACGTCTTGAGTCTAA
GCCAGGGAATCAGCTCTCATCTCGAGGCAATTTGGGGTACACGGAGCAGTCCTCGAGTTGCTCTGCTGAACTTGGTACTCCTC
TAGACTTGGCCCGTTGTTCTCGGGGAACCTCTGGAGTTGCCTAAAGGGAGTCAAGCCTCAGGTCGTGTTTGATGGGGAACGGG
GGATGGCTCTGGAGTCAATGCAGGGGAAGCGGGCCTCATCTCGAGTTGATTTGGGGCCCATGGAGCTCCTTCGCGATGCTGG
TGTGACCTCAGGGTCCCTCTCTACTTGTGACAGTGTTTTT-GCGGACTGTCTCGACATCCATCAAGCAAGTCGAGGCTCCTGAC-
------------------------------------------------------------------------------------------------------------------------------------------------------------------------------------
------------------------------------------------------------------------------------------------------------------------------------------------------------------------------------
----------------------------------------------------------------------------------------------------------------------------------------------------------------
>g3-g4
ACTGTATTCAAGCCACTGCAAGGAAACCCGGCCTCCTTTCGAGTCAGGGCTTCTCGGTGTATATGCCCATTGAGGCAGCAATCT
CAGGGTCCCTCCCTCATACCTATTGCTGAGAGAAGCCTCCTCTTGAGGGGCTTGTGGAAAGGTGGCCTACGTCTTGAGTCTAA
GCCAGGGAATCAGCTCTCATCTCGAGGCAATTTGGGGTACACGGAGCAGTCCTCGAGTTGCTCTGCTGAACTTGGTACTCCTC
TAGACTTGGCCCGTTGTTCTCGGGGAACCTCTGGAGTTGCCTAAAGGGAGTCAAGCCTCAGGTCGTGTTTGATGGGGAACGGG
GGATGGCTCTGGAGTCAATGCAGGGGAAGCGGGCCTCATCTCGAGTTGATTTGGGGCCCATGGAGCTCCTTCGCGATGCTGG
TGTGACCTCAGGGTCCCTCTCTACTTGTGACAGTGTTTTT-GCGGACTGTCTCGACATCCATCAAGCAAGTCGAGGCTCCTGAC-
------------------------------------------------------------------------------------------------------------------------------------------------------------------------------------
------------------------------------------------------------------------------------------------------------------------------------------------------------------------------------
----------------------------------------------------------------------------------------------------------------------------------------------------------------
>g13-g14
ACTGTATTCAAGCCACTGCAAGGAAACCCGGCCTCCTTTCGAGTCAGGGCTTCTCGGTGTATATGCCCATTGAGGCAGCAATCT
CAGGGTCCCTCCCTCATACCTATTGCTGAGAGAAGCCTCCTCTTGAGGGGCTTGTGGAAAGGTGGCCTACGTCTTGAGTCTAA
GCCAGGGAATCAGCTCTCATCTCGAGGCAATTTGGGGTACACGGAGCAGTCCTCGAGTTGCTCTGCTGAACTTGGTACTCCTC
TAGACTTGGCCCGTTGTTCTCGGGGAACCTCTGGAGTTGCCTAAAGGGAGTCAAGCCTCAGGTCGTGTTTGATGGGGAACGGG
GGATGGCTCTGGAGTCAATGCAGGGGAAGCGGGCCTCATCTCGAGTTGATTTGGGGCCCATGGAGCTCCTTCGCGATGCTGG
TGTGACCTCAGGGTCCCTCTCTACTTGTGACAGTGTTTTT-GCGGACTGTCTCGACATCCATCAAGCAAGTCGAGGCTCCTGAC-

------------------------------------------------------------------------------------------------------------------------------------
------------------------------------------------------------------------------------------------------------------------------------
------------------------------------------------------------------------------------------------------------------------------
>g19-g20
ACTGTATTCAAGCCACTGCAAGGAAACCCGGCCTCCTTTCGAGTCAGGGCTTCTCGGTGTATATGCCCATTGAGGCAGCAATCT
CAGGGTCCCTCCCTCATACCTATTGCTGAGAGAAGCCTCCTCTTGAGGGGCTTGTGGAAAGGTGGCCTACGTCTTGAGTCTAA
GCCAGGGAATCAGCTCTCATCTCGAGGCAATTTGGGGTACACGGAGCAGTCCTCGAGTTGCTCTGCTGAACTTGGTACTCCTC
TAGACTTGGCCCGTTGTTCTCGGGGAACCTCTGGAGTTGCCTAAAGGGAGTCAAGCCTCAGGTCGTGTTTGATGGGGAACGGG
GGATGGCTCTGGAGTCAATGCAGGGGAAGCGGGCCTCATCTCGAGTTGATTTGGGGCCCATGGAGCTCCTTCGCGATGCTGG
TGTGACCTCAGGGTCCCTCTCTACTTGTGACAGTGTTTTT-GCGGACTGTCTCGACATCCATCAAGCAAGTCGAGGCTCCTGAC-
------------------------------------------------------------------------------------------------------------------------------------
------------------------------------------------------------------------------------------------------------------------------------
------------------------------------------------------------------------------------------------------------------------------
>g20-g21
ACTGTATTCAAGCCACTGCAAGGAAACCCGGCCTCCTTTCGAGTCAGGGCTTCTCGGTGTATATGCCCATTGAGGCAGCAATCT
CAGGGTCCCTCCCTCATACCTATTGCTGAGAGAAGCCTCCTCTTGAGGGGCTTGTGGAAAGGTGGCCTACGTCTTGAGTCTAA
GCCAGGGAATCAGCTCTCATCTCGAGGCAATTTGGGGTACACGGAGCAGTCCTCGAGTTGCTCTGCTGAACTTGGTACTCCTC
TAGACTTGGCCCGTTGTTCTCGGGGAACCTCTGGAGTTGCCTAAAGGGAGTCAAGCCTCAGGTCGTGTTTGATGGGGAACGGG
GGATGGCTCTGGAGTCAATGCAGGGGAAGCGGGCCTCATCTCGAGTTGATTTGGGGCCCATGGAGCTCCTTCGCGATGCTGG
TGTGACCTCAGGGTCCCTCTCTACTTGTGACAGTGTTTTT-GCGGACTGTCTCGACATCCATCAAGCAAGTCGAGGCTCCTGAC-
------------------------------------------------------------------------------------------------------------------------------------
------------------------------------------------------------------------------------------------------------------------------------
------------------------------------------------------------------------------------------------------------------------------
>U53517.1
---------------------------
CCGGCCTCCTTTCGAGTCAGGGCTTCTCGGTGTATATGCCCATTGAGGCAGCAATCTCAGGGTCCCTCCTTCATACCTATTGCT
GAGAGAAGCCTCCTCTTGAGGGGCTTGTGGAAAGGTGGCCTACGTATTGAGTCTAAGCCAGGGAATCAGCTCTCATCTCGAGG
CAATTTGGGGTACACGGAGCAGTCCTCGAGTTGCTCTGCTGAACTTGGTACTCGTCTAGACTTGGCCCGTTGTTCTCCAGGAAC
CTCTGGAGTTGCCTAAAGGGAGTCAAGCCTCAGGTCGTGTTTGATGGGGAACTGGGGATGGCTCTGGAGTCAATGCAGGGGA
AGCGGGCCTCATCTCAAGTTGATTTGGGGCCCATGGAGCTCCTTCGCGATGCTGGTGTGACCTCAGGGTCCCTCTCTACTTGT
GACAGTGTTTTTGGCGGACTGTCTCGACATCCATCAAGCAAGTCGAGGCTCCTGACATGTTTCAGGGGTCTCACGGAATTGNTA
TGCATGCAGTGCAGGGGAATCGGGCCTCATATCTCGCTTGGAGGGGGGAAGTCTCATGGTTTTTCTCGAGTTCCGGCCCGGACC
TGGGGTACATTCTCGACTTTACGGCGGGGATGCCCCTTCAACCCTCGTGTTTGNTCAACGAAGTCAGGAGTCCTGTTGAGTTAC
GAGGGACACCTCAGGAGTCTCTTCGAGGCTTGGCAGGGCAAAAGGGACGCTTCTCGAGGTGAGTCGGGAGACCCAGGGTCC
CTTTCCAGTAGCCACAGGGATACTGGGATTCCTGTCAATGTTCAAGAGGAGTCAGGCTCCGTCAAATTTTGAAANAGTGAGCTC
TGCGTGCTTCTCGAGGTGTCAGAGGCATGTGAGGCATTCCGTCGAGAGGAGTCGGGGACCTAGGGCTTTCTCTAGGGACTCC
ACAGGTGGTGCAGACATCCCTTCACCTTGTGAGATGAAAGACTAGACTGTATTCAAGCCACTGCAAGGAAACCCGG
>g8-g9
ACTGTATTCAAGCCACTGCAAGGAAACCCGGCCTCCTTTCGAGTCAGGGCTTCTCGGTGTATATGCCCATTGAGGCAGCAATCT
CAGGGTCCCTCCCTCATACCTATTGCTGAGAGAAGCCTCCTCTTGAGGGGCTTGTGGAAAGGTGGCCTACGTCTTGAGTCTAA
GCCAGGGAATCAGCTCTCATCTCGAGGCAATTTGGGGTACACGGAGCAGTCCTCGAGTTGCTCTGCTGAACTTGGTACTCGTC
TAGACTTGGCCCGTTGTTCTCCGGGAACCTCTGGAGTTGCCTAAAGGGAGTCAAGCCTCAGGTCGTGTTTGATGGGGAACTGG
GGATGGCTCTGGAGTCAATGCAGGGGAAGCGGGCCTCATCTCGAGTTGATTTGGGGCCCATGGAGCTCCTTCGCGATGCTGG
TGTGACCTCAGGGTCCCTCTCTACTTGTGACAGTGTTTTT-GCGGACTGTCTCGACATCCATCAAGCAAGTCGAGGCTCCTGAC-
------------------------------------------------------------------------------------------------------------------------------------
------------------------------------------------------------------------------------------------------------------------------------
------------------------------------------------------------------------------------------------------------------------------
>g10-g11
ACTGTATTCAAGCCACTGCAAGGAAACCCGGCCTCCTTTCGAGTCAGGGCTTCTCGGTGTATATGCCCATTGAGGCAGCAATCT
CAGGGTCCCTCCCTCATACCTATTGCTGAGAGAAGCCTCCTCTTGAGGGGCTTGTGGAAAGGTGGCCTACGTCTTGAGTCTAA
GCCAGGGAATCAGCTCTCATCTCGAGGCAATTTGGGGTACACGGAGCAGTCCTCGAGTTGCTCTGCTGAACTTGGTACTCCTC
TAGACTTGGCCCGTTGTTCTCCGGGAACCTCTGGAGTTGCCTAAAGGGAGTCAAGCCTCAGGTCGTGTTTGATGGGGAACAGG
GGATGGCTCTGGAGTCAATGCAGGGGAAGCGGGCCTCATCTCGAGTTGATTTGGGGCCCATGGAGCTCCTTCGCGATGCTGG
TGTGACCTCAGGGTCCCTCTCTACTTGTGACAGTGTTTTT-GCGGACTGTCTCGACATCCATCAAGCAAGTCGAGGCTCCTGAC-
------------------------------------------------------------------------------------------------------------------------------------
------------------------------------------------------------------------------------------------------------------------------------
------------------------------------------------------------------------------------------------------------------------------
>g5-g6
ACTGTATTCAAGCCACTGCAAGGAAACCCGGCCTCCTTTCGAGTCAGGGCTTCTCGGTGTATATGCCCATTGAGGCAGCAATCT
CAGGGTCCCTCCCTCATACCTATTGCTGAGAGAAGCCTCCTCTTGAGGGGCTTGTGGAAAGGTGGCCTACGTCTTGAGTCTAA
GCCAGGGAATCAGCTCTCATCTCGAGGCAATTTGGGGTACACGGAGCAGTCCTCGAGTTGCTCTGCTGAACTTGGTACTCCTC
TAGACTTGGCCCGTTGTTCTCCGGGAACCTCTGGAGTTGCCTAAAGGGAGTCAAGCCTCAGGTCGTGTTTGATGGGGAACGGG
GGATGGCTCTGGAGTCAATGCAGGGGAAGCGGGCCTCATCTCGAGTTGATTTGGGGCCCATGGAGCTCCTTCGCGATGCTGG
TGTGACCTCAGGGTCCCTCTCTACTTGTGACAGTGTTTTT-GCGGACTGTCTCGACATCCATCAAGCAAGTCGAGGCTCCTGAC-
------------------------------------------------------------------------------------------------------------------------------------
------------------------------------------------------------------------------------------------------------------------------------
------------------------------------------------------------------------------------------------------------------------------
>g4-g5
ACTGTATTCAAGCCACTGCAAGGAAACCCGGCCTCCTTTCGAGTCAGGGCTTCTCGGTGTATATGCCCATTGAGGCAGCAATCT
CAGGGTCCCTCCCTCATACCTATTGCTGAGAGAAGCCTCCTCTTGAGGGGCTTGTGGAAAGGTGGCCTACGTCTTGAGTCTAA

GCCAGGGAATCAGCTCTCATCTCGAGGCAATTTGGGGTACACGGAGCAGTCCTCGAGTTGCTCTGCTGAACTTGGTACTCCTC
TAGACTTGGCCCGTTGTTCTCCGGGAACCTCTGGAGTTGCCTAAAGGGAGTCAAGCCTCAGGTCGTGTTTGATGGGGAACGGG
GGATGGCTCTGGAGTCAATGCAGGGGAAGCGGGCCTCATCTCGAGTTGATTTGGGGCCCATGGAGCTCCTTCGCGATGCTGG
TGTGACCTCAGGGTCCCTCTCTACTTGTGACAGTGTTTTT-GCGGACTGTCTCGACATCCATCAAGCAAGTCGAGGCTCCTGAC-
----------------------------------------------------------------------------------------------------------------------------------------------------------------
----------------------------------------------------------------------------------------------------------------------------------------------------------------
-----------------------------------------------------------------------------------------------------------------------------------------------------

>g25-g26
ACTGTATTCAAGCCACTGCAAGGAAACCCGGCCTCCTTTCGAGTCAGGGCTTCTCGGTGTATATGCCCATTGAGGCAGCAATCT
CAGGGTCCCTCCCTCATACCTATTGCTGAGAGAAGCCTCCTCTTGAGGGGCTTGTGGAAAGGTGGCCTACGTCTTGAGTCTAA
GCCAGGGAATCAGCTCTCATCTCGAGGCAATTTGGGGTACACGGAGCAGTCCTCGAGTTGCTCTGCTGAACTTGGTACTCCTC
TAGACTTGGCCCGTTGTTCTCCGGGAACCTCTGGAGTTGCCTAAAGGGAGTCAAGCCTCAGGTCGTGTTTGATGGGGAACGGG
GGATGGCTCTGGAGTCAATGCAGGGGAAGCGGGCCTCATCTCGAGTTGATTTGGGGCCCATGGAGCTCCTTCGCGATGCTGG
TGTGACCTCAGGGTCCCTCTCTACTTGTGACAGTGTTTTT-
GCGGACTGTCTCGACATCCATCAAGCAAGTCGAGGCTCCTGACATGTTTCAGGGGGCTCACGGAATTGCTCTG--------------------
----------------------------------------------------------------------------------------------------------------------------------------------------------------
----------------------------------------------------------------------------------------------------------------------------------------------------------------
---------------------------------------------------------------------------------------------------------C

>g17-g18
ACTGTATTCAAGCCACTGCAAGGAAACCCGGCCTCCTTTCGAGTCAGGGCTTCTCGGTGTATATGCCCATTGAGGCAGCAATCT
CAGGGTCCCTCCCTCATACCTATTGCTGAGAGAAGCCTCCTCTTGAGGGGCTTGTGGAAAGGTGGCCTACGTCTTGAGTCTAA
GCCAGGGAATCAGCTCTCATCTCGAGGCAATTTGGGGTACACGGAGCAGTCCTCGAGTTGCTCTGCTGAACTTGGTACTCCTC
TAGACTTGGCCCGTTGTTCTCG-
GGAACCTCTGGAGTTGCCTAAAGGGAGTCAAGCCTCAGGTCGTGTTTGATGGGGAACGGGGGATGGCTCTGGAGTCAATGCA
GGGGAAGCGGGCCTCATCTCGAGTTGATTTGGGGCCCATGGAGCTCCTTCGCGATGCTGGTGTGACCTCAGGGTCCCTCTCT
ACTTGTGACAGTGTTTTT-GCGGACTGTCTCGACATCCATCAAGCAAGTCGAGGCTCCTGAC------------------------------------------
----------------------------------------------------------------------------------------------------------------------------------------------------------------
----------------------------------------------------------------------------------------------------------------------------------------------------------------
----------------------------------------------------------------------------------------------------------------

>g7-g8
ACTGTATTCAAGCCACTGCAAGGAAACCCGGCCTCCTTTCGAGTCAGGGCTTCTCGGTGTATATGCCCATTGAGGCAGCAATCT
CAGGGTCCCTCCCTCATACCTATTGCTGAGAGAAGCCTCCTCTTGAGGGGCTTGTGGAAAGGTGGCCTACGTCTTGAGTCTAA
GCCAGGGAATCAGCTCTCATCTCGAGGCAATTTGGGGTACACGGAGCAGTCCTCGAGTTGCTCTGCTGAACTTGGTACTCCTC
TAGACTTGGCCCGTTGTTCTCGGGGAACCTCTGGAGTTGCCTAAAGGGAGTCAAGCCTCAGGTCGTGTTTGATGGGGAACGGG
GGATGGCTCTGGAGTCAATGCAGGGGAAGCGGGCCTCATCTCGAGTTGATTTGGGGCCCATGGAGCTCCTTCGCGATGCTGG
TGTGACCTCAGGGTCCCTCTCTACTTGTGACAGTGTTTTT-
GCGGACTGTCTCGACATCCATCAAGCAAGTCGAGGCTCCTGACATGTTTCAGGGGGCTCACGGAATTGCTCTG--------------------
----------------------------------------------------------------------------------------------------------------------------------------------------------------
----------------------------------------------------------------------------------------------------------------------------------------------------------------
---------------------------------------------------------------------------------------------------------C

>g24-g25
ACTGTATTCAAGCCACTGCAAGGAAACCCGGCCTCCTTTCGAGTCAGGGCTTCTCGGTGTATATGCCCATTGAGGCAGCAATCT
CAGGGTCCCTCCCTCATACCTATTGCTGAGAGAAGCCTCCTCTTGAGGGGCTTGTGGAAAGGTGGCCTACGTCTTGAGTCTAA
GCCAGGGAATCAGCTCTCATCTCGAGGCAATTTGGGGTACACGGAGCAGTCCTCGAGTTGCTCTGCTGAACTTGGTACTCCTC
TAGACTTGGCCCGTTGTTCTCGGGGAACCTCTGGAGTTGCCTAAAGGGAGTCAAGCCTCAGGTCGTGTTTGATGGGGAACGGG
GGATGGCTCTGGAGTCAATGCAGGGGAAGCGGGCCTCATCTCGAGTTGATTTGGGGCCCATGGAGCTCCTTCGCGATGCTGG
TGTGACCTCAGGGTCCCTCTCTACTTGTGACAGTGTTTTT-
GCGGACTGTCTCGACATCCATCAAGCAAGTCGAGGCTCCTGACATGTTTCAGGGGGCTCACGGAATTGCTCTG--------------------
----------------------------------------------------------------------------------------------------------------------------------------------------------------
----------------------------------------------------------------------------------------------------------------------------------------------------------------
---------------------------------------------------------------------------------------------------------C

>g2-g3
ACTGTATTCAAGCCACTGCAAGGAAACCCGGCCTCCTTTCGAGTCAGGGCTTCTCGGTGTATATGCCCATTGAGGCAGCAATCT
CAGGGTCCCTCCCTCATACCTATTGCTGAGAGAAGCCTCCTCTTGAGGGGCTTGTGGAAAGGTGGCCTACGTCTTGAGTCTAA
GCCAGGGAATCAGCTCTCATCTCGAGGCAATTTGGGGTACACGGAGCAGTCCTCGAGTTGCTCTGCTGAACTTGGTACTCCTC
TAGACTTGGCCCGTTGTTCTCGGGGAACCTCTGGAGTTGCCTAAAGGGAGTCAAGCCTCAGGTCGTGTTTGATGGGGAACGGG
GGATGGCTCTGGAGTCAATGCAGGGGAAGCGGGCCTCATCTCGAGTTGATTTGGGGCCCATGGAGCTCCTTCGCGATGCTGG
TGTGACCTCAGGGTCCCTCTCTACTTGTGACAGTGTTTTT-
GCGGACTGTCTCGACATCCATCAAGCAAGTCGAGGCTCCTGACATGTTTCAGGGGGCTCACGGAATTGCTCTG--------------------
----------------------------------------------------------------------------------------------------------------------------------------------------------------
----------------------------------------------------------------------------------------------------------------------------------------------------------------
---------------------------------------------------------------------------------------------------------C

>g21-g22
ACTGTATTCAAGCCACTGCAAGGAAACCCGGCCTCCTTTCGAGTCAGGGCTTCTCGGTGTATATGCCCATTGAGGCAGCAATCT
CAGGGTCCCTCCCTCATACCTATTGCTGAGAGAAGCCTCCTCTTGAGGGGCTTGTGGAAAGGTGGCCTACGTCTTGAGTCTAA
GCCAGGGAATCAGCTCTCATCTCGAGGCAATTTGGGGTACACGGAGCAGTCCTCGAGTTGCTCTGCTGAACTTGGTACTCCTC
TAGACTTGGCCCGTTGTTCTCGGGGAACCTCTGGAGTTGCCTAAAGGGAGTCAAGCCTCAGGTCGTGTTTGATGGGGAACGGG
GGATGGCTCTGGAGTCAATGCAGGGGAAGCGGGCCTCATCTCGAGTTGATTTGGGGCCCATGGAGCTCCTTCGCGATGCTGG
TGTGACCTCAGGGTCCCTCTCTACTTGTGACAGTGTTTTT-
GCGGACTGTCTCGACATCCATCAAGCAAGTCGAGGCTCCTGACATGTTTCAGGGGGCTCACGGAATTGCTCTG--------------------

-----------------------------------------------------------------------------------------------------------------------------------------
-----------------------------------------------------------------------------------------------------------------------------------------
--------------------------------------------------------------------------------------------------------------C
>g22-g23
ACTGTATTCAAGCCACTGCAAGGAAACCCGGCCTCCTTTCGAGTCAGGGCTTCTCGGTGTATATGCCCATTGAGGCAGCAATCT
CAGGGTCCCTCCCTCATACCTATTGCTGAGAGAAGCCTCCTCTTGAGGGGCTTGTGGAAAGGTGGCCTACGTCTTGAGTCTAA
GCCAGGGAATCAGCTCTCATCTCGAGGCAATTTGGGGTACACGGAGCAGTCCTCGAGTTGCTCTGCTGAACTTGGTACTCCTC
TAGACTTGGCCCGTTGTTCTCGGGGAACCTCTGGAGTTGCCTAAAGGGAGTCAAGCCTCAGGTCGTGTTTGATGGGGAACGGG
GGATGGCTCTGGAGTCAATGCAGGGGAAGCGGGCCTCATCTCGAGTTGATTTGGGGCCCATGGAGCTCCTTCGCGATGCTGG
TGTGACCTCAGGGTCCCTCTCTACTTGTGACAGTGTTTTT-
GCGGACTGTCTCGACATCCATCAAGCAAGTCGAGGCTCCTGACATGTTTCAGGGGGCTCACGGAATTGCTCTG---------------------
-----------------------------------------------------------------------------------------------------------------------------------------
-----------------------------------------------------------------------------------------------------------------------------------------
--------------------------------------------------------------------------------------------------------------C
>g27-g28
ACTGTATTCAAGCCACTGCAAGGAAACCCGGCCTCCTTTCGAGTCAGGGCTTCTCGGTGTATATGCCCATTGAGGCAGCAATCT
CAGGGTCCCTCCCTCATACCTATTGCTGAGAGAAGCCTCCTCTTGAGGGGCTTGTGGAAAGGTGGCCTACGTCTTGAGTCTAA
GCCAGGGAATCAGCTCTCATCTCGAGGCAATTTGGGGTACACGGAGCAGTCCTCGAGTTGCTCTGCTGAACTTGGTACTCCTC
TAGACTTGGCCCGTTGTTCTCGGGGAACCTCTGGAGTTGCCTAAAGGGAGTCAAGCCTCAGGTCGTGTTTGATGGGGAACGGG
GGATGGCTCTGGAGTCAATGCAGGGGAAGCGGGCCTCATCTCGAGTTGATTTGGGGCCCATGGAGCTCCTTCGCGATGCTGG
TGTGACCTCAGGGTCCCTCTCTACTTGTGACAGTGTTTTT-
GCGGACTGTCTCGACATCCATCAAGCAAGTCGAGGCTCCTGACATGTTTCAGGGGGCTCACGGAATTGCTCTGCATGCAGTGC
AGGGGAATCGGGCCTC--ATCTCGCTTGG----------------------------------------------------------------------------------------------------
-----------------------------------------------------------------------------------------------------------------------------------------
-----------------------------------------------------------------------------------------------------------------------------------AGGGGAAGTCTC

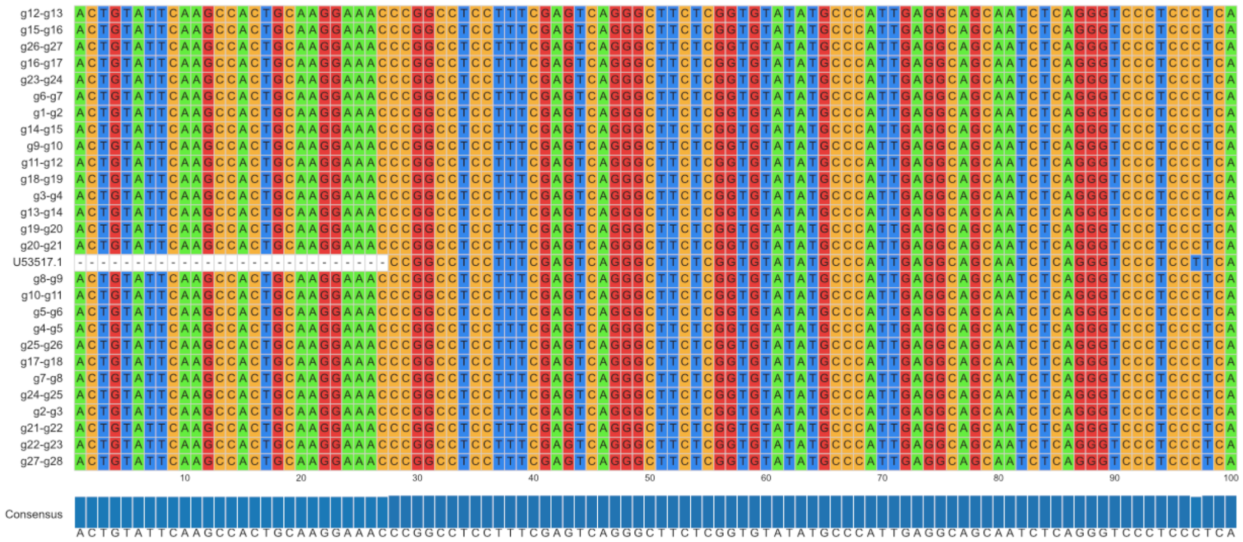

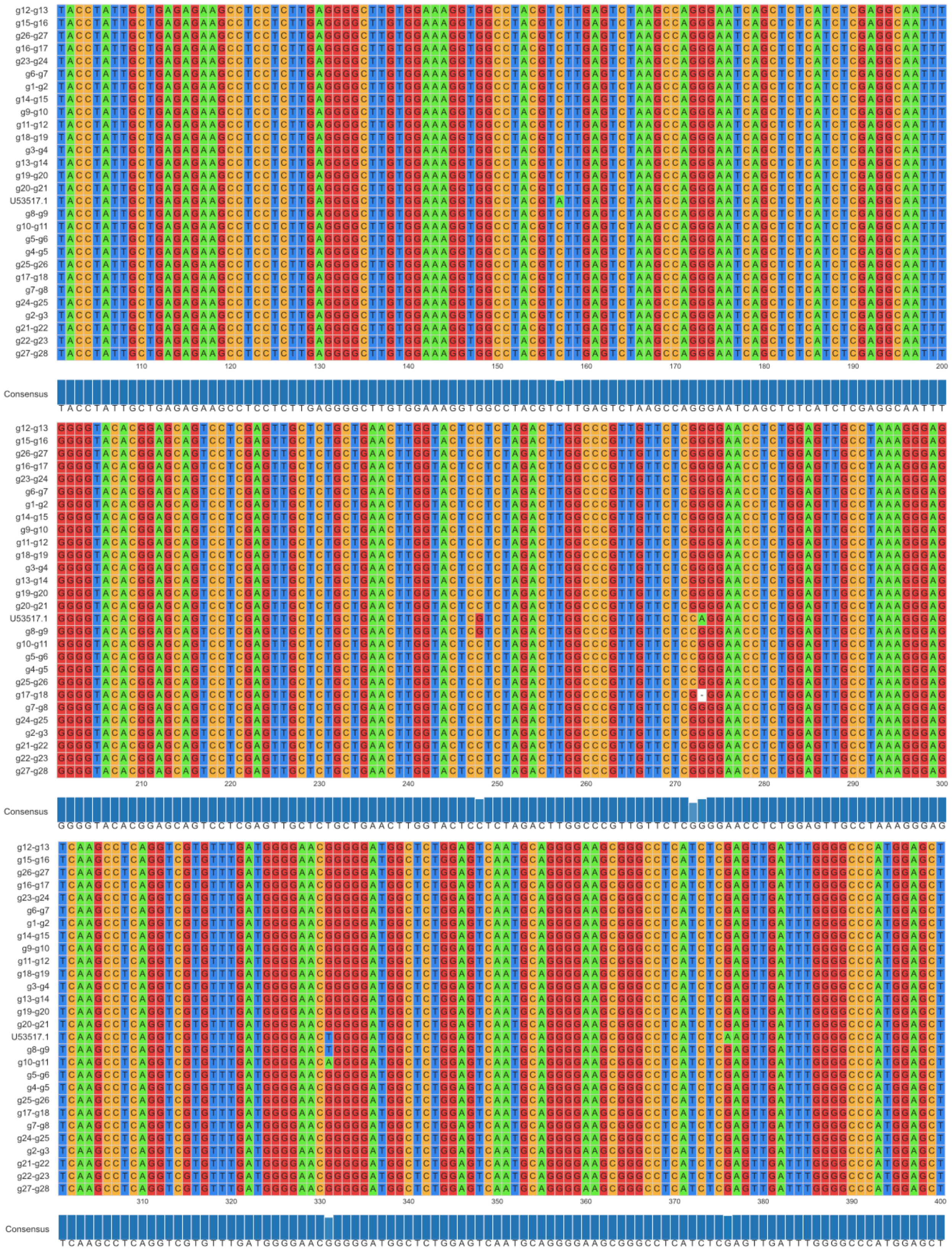
g12-g13
g15-g16
g26-g27
g16-g17
g23-g24
g6-g7
g1-g2
g14-g15
g9-g10
g11-g12
g18-g19
g3-g4
g13-g14
g19-g20
g20-g21
U53517.1
g8-g9
g10-g11
g5-g6
g4-g5
g25-g26
g17-g18
g7-g8
g24-g25
g2-g3
g21-g22
g22-g23
g27-g28
110 120 130 140 150 160 170 180 190 200
Consensus
TACCTATTGCTGAGAGAAGCCTCCTCTTGAGGGGCTTGTGGAAAGGTGGCCTACGTCTTGAGTCTAAGCCAGGGAATCAGCTCTCATCTCGAGGCAATTT
210 220 230 240 250 260 270 280 290 300
Consensus
GGGGTACACGGAGCAGTCCTCGAGTTGCTCTGCTGAACTTGGTACTCCTCTAGACTTGGCCCGTTGTTCTCGGGGAACCTCTGGAGTTGCCTAAAGGGAG
310 320 330 340 350 360 370 380 390 400
Consensus
TCAAGCCTCAGGTCGTGTTTGATGGGGAACGGGGGATGGCTCTGGAGTCAATGCAGGGGAAGCGGGCCTCATCTCGAGTTGATTTGGGGCCCATGGAGCT

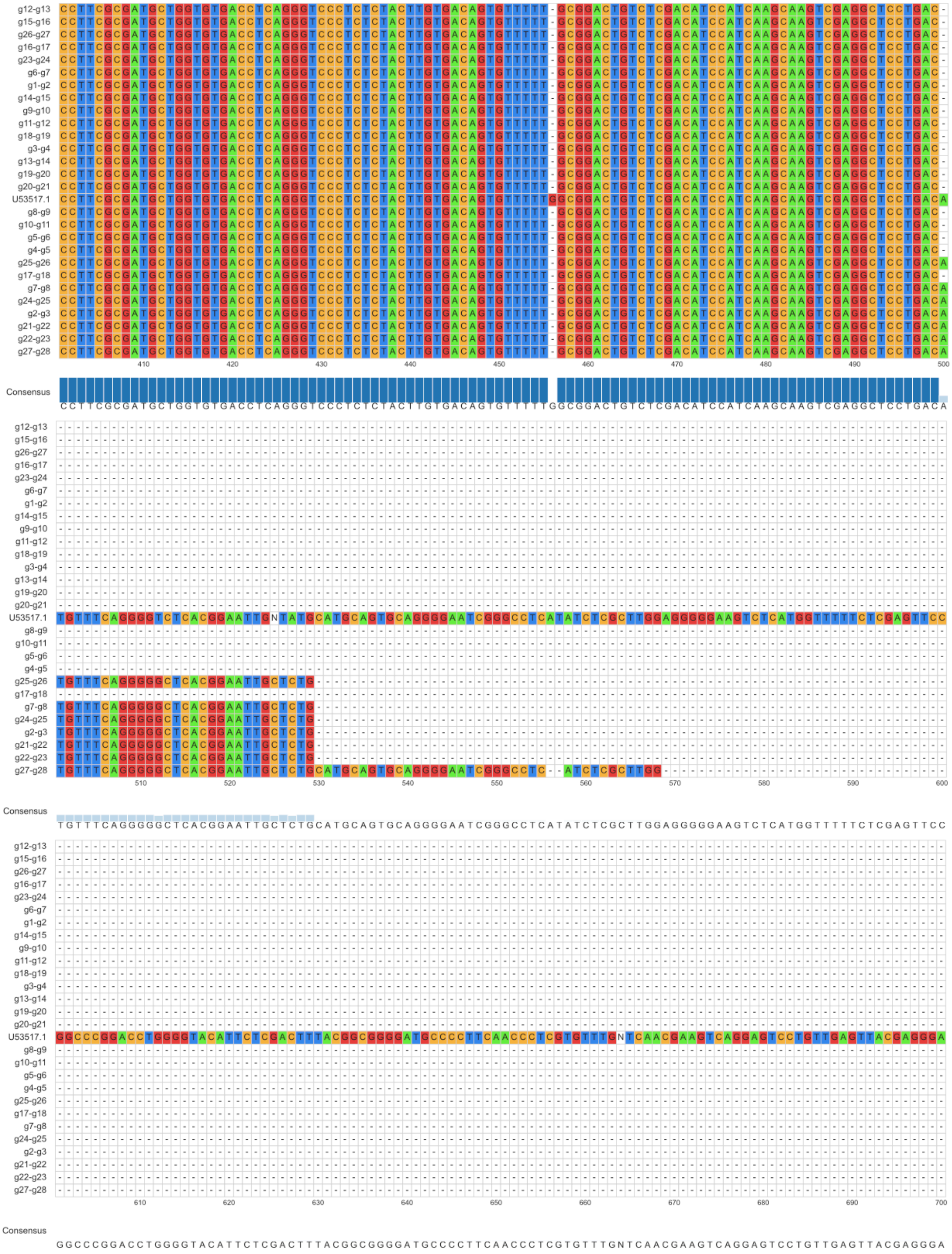
g12-g13
g15-g16
g26-g27
g16-g17
g23-g24
g6-g7
g1-g2
g14-g15
g9-g10
g11-g12
g18-g19
g3-g4
g13-g14
g19-g20
g20-g21
U53517.1
g8-g9
g10-g11
g5-g6
g4-g5
g25-g26
g17-g18
g7-g8
g24-g25
g2-g3
g21-g22
g22-g23
g27-g28
410 420 430 440 450 460 470 480 490 500
Consensus
CCTTCGCGATGCTGGTGTGACCTCAGGGTCCCTCTCTACTTGTGACAGTGTTTTTGGCGGACTGTCTCGACATCCATCAAGCAAGTCGAGGCTCCTGACA
510 520 530 540 550 560 570 580 590 600
Consensus
TGTTTCAGGGGGCTCACGGAATTGCTCTGCATGCAGTGCAGGGGAATCGGGCCTCATATCTCGCTTGGAGGGGGAAGTCTCATGGTTTTTCTCGAGTTCC
610 620 630 640 650 660 670 680 690 700
Consensus
GGCCCGGACCTGGGGTACATTCTCGACTTTACGGCGGGGATGCCCCTTCAACCCTCGTGTTTGNTCAACGAAGTCAGGAGTCCTGTTGAGTTACGAGGGA

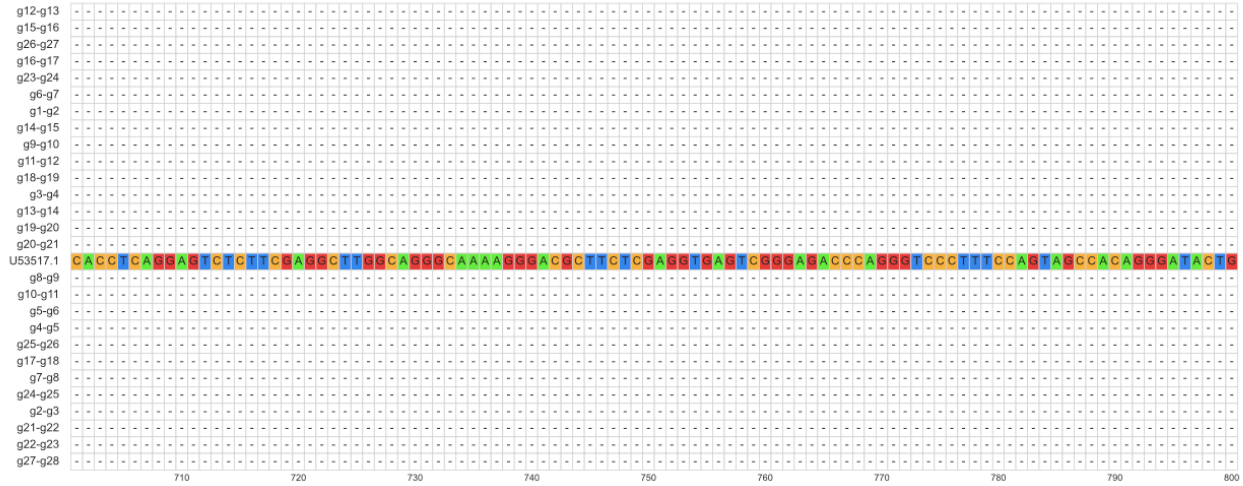


Consensus

C A C C T C A G G A G T C T C T T C G A G G C T T G G C A G G G C A A A A G G G A C G C T T C T C G A G G T G A G T C G G G A G A C C C A G G G T C C C T T T C C A G T A G C C A C A G G G A T A C T G

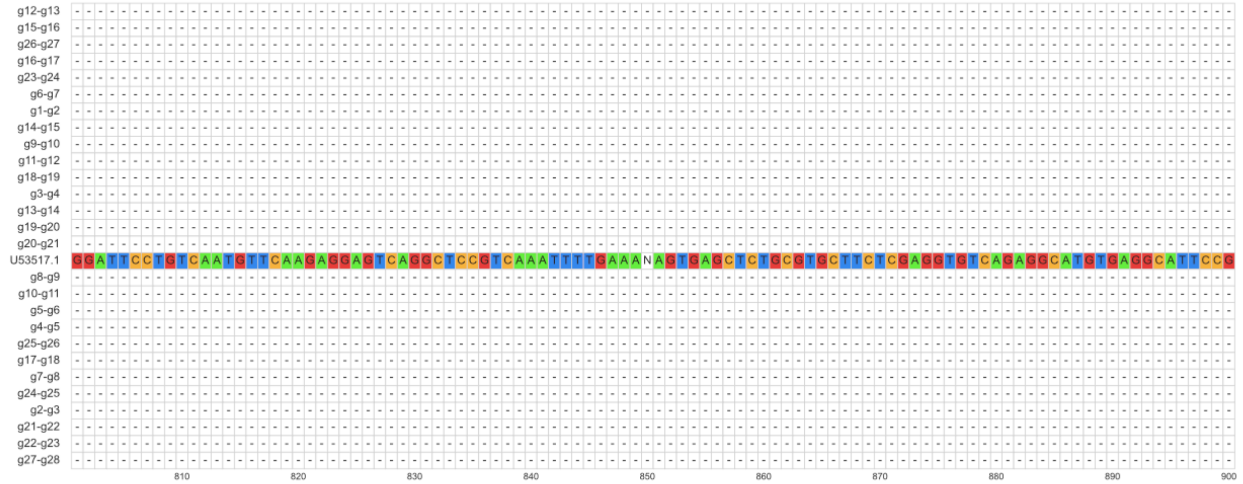


Consensus

G G A T T C C T G T C A A T G T T C A A G A G G A G T C A G G C T C C G T C A A A T T T T G A A A N A G T G A G C T C T G C G T G C T T C T C G A G G T G T C A G A G G C A T G T G A G G C A T T C C G

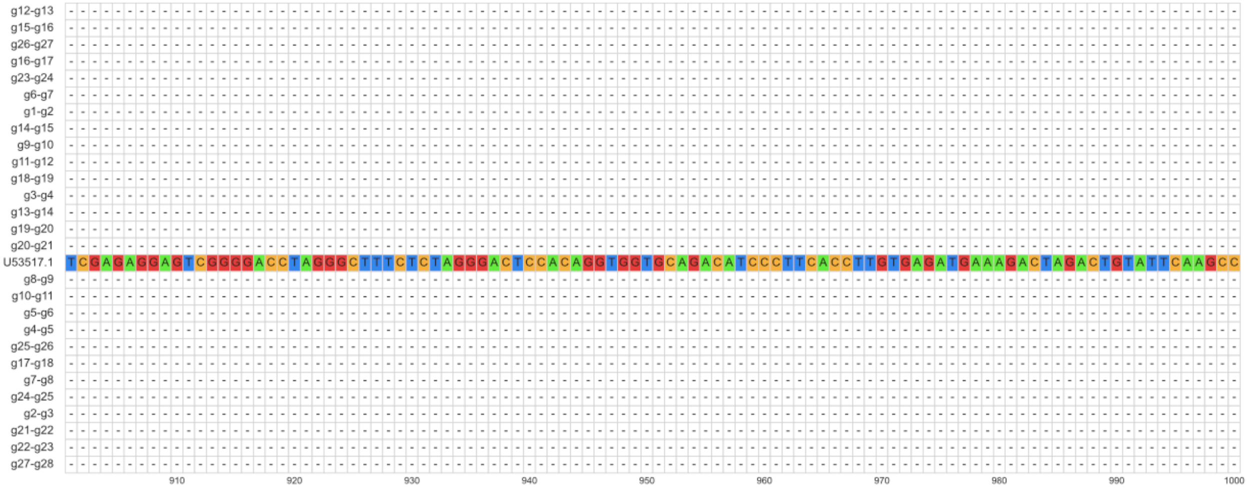

| | | | | | | | | | | | | | | | | | |
|---|---|---|---|---|---|---|---|---|---|---|---|---|---|---|---|---|---|
| g12-g13 | - | - | - | - | - | - | - | - | - | - | - | - | - | - | - | - | - |
| g15-g16 | - | - | - | - | - | - | - | - | - | - | - | - | - | - | - | - | - |
| g26-g27 | - | - | - | - | - | - | - | - | - | - | - | - | - | - | - | - | - |
| g16-g17 | - | - | - | - | - | - | - | - | - | - | - | - | - | - | - | - | - |
| g23-g24 | - | - | - | - | - | - | - | - | - | - | - | - | - | - | - | - | - |
| g6-g7 | - | - | - | - | - | - | - | - | - | - | - | - | - | - | - | - | - |
| g1-g2 | - | - | - | - | - | - | - | - | - | - | - | - | - | - | - | - | - |
| g14-g15 | - | - | - | - | - | - | - | - | - | - | - | - | - | - | - | - | - |
| g9-g10 | - | - | - | - | - | - | - | - | - | - | - | - | - | - | - | - | - |
| g11-g12 | - | - | - | - | - | - | - | - | - | - | - | - | - | - | - | - | - |
| g18-g19 | - | - | - | - | - | - | - | - | - | - | - | - | - | - | - | - | - |
| g3-g4 | - | - | - | - | - | - | - | - | - | - | - | - | - | - | - | - | - |
| g13-g14 | - | - | - | - | - | - | - | - | - | - | - | - | - | - | - | - | - |
| g19-g20 | - | - | - | - | - | - | - | - | - | - | - | - | - | - | - | - | - |
| g20-g21 | - | - | - | - | - | - | - | - | - | - | - | - | - | - | - | - | - |
| U53517.1 | A | C | T | G | C | A | A | G | G | A | A | A | C | C | C | G | G |
| g8-g9 | - | - | - | - | - | - | - | - | - | - | - | - | - | - | - | - | - |
| g10-g11 | - | - | - | - | - | - | - | - | - | - | - | - | - | - | - | - | - |
| g5-g6 | - | - | - | - | - | - | - | - | - | - | - | - | - | - | - | - | - |
| g4-g5 | - | - | - | - | - | - | - | - | - | - | - | - | - | - | - | - | - |
| g25-g26 | - | - | - | - | - | - | - | - | - | - | - | - | - | - | - | - | C |
| g17-g18 | - | - | - | - | - | - | - | - | - | - | - | - | - | - | - | - | - |
| g7-g8 | - | - | - | - | - | - | - | - | - | - | - | - | - | - | - | - | C |
| g24-g25 | - | - | - | - | - | - | - | - | - | - | - | - | - | - | - | - | C |
| g2-g3 | - | - | - | - | - | - | - | - | - | - | - | - | - | - | - | - | C |
| g21-g22 | - | - | - | - | - | - | - | - | - | - | - | - | - | - | - | - | C |
| g22-g23 | - | - | - | - | - | - | - | - | - | - | - | - | - | - | - | - | C |
| g27-g28 | - | - | - | - | - | A | G | G | G | G | A | A | G | T | C | T | C |

1010

Consensus

A C T G C A X G G X A A X X C X C
